# Conserved core RNAi machinery in trematode-vectoring snails indicates gene silencing potential in the absence of classical systemic and amplification effectors

**DOI:** 10.64898/2026.07.10.737666

**Authors:** Damilare O Famakinde, Ciaran Lonergan, Geoffrey N Gobert, Duncan Wells, Paul McVeigh

## Abstract

RNA interference (RNAi) is a widely exploited reverse-genetics tool with potential uses for disease control. Successful RNAi has been reported in trematode-vectoring snails, but the composition of RNAi effector-encoding gene complements, a key driver for RNAi efficiency, remain unstudied in these species. Using bioinformatics and comparative genomics, we searched for orthologues of 115 RNAi effector sequences in genomes or transcriptomes of four snail vectors: *Biomphalaria glabrata*, *B. pfeifferi*, *Bulinus truncatus*, and *Lymnaea staginalis*. Gene expression patterns of selected RNAi effectors were then examined across developmental stages and tissues of the model *B. glabrata* snail. At least 74 RNAi-related proteins were conserved across all four species, including core components known to be essential for gene silencing. Classical systemic RNAi-deficient (SID) genes that facilitate systemic RNAi in other systems were absent, suggesting that alternative pathways may compensate for dsRNA uptake and transport. Core effectors of secondary RNAi amplification and heritable RNAi were not detected. Expressions of *Dicer-1*, *Argonaute-2*, and the exonuclease *Eri-1* did not vary significantly with snail size. A putative RNAi-inhibiting Staufen orthologue showed elevated expression in the ovotestis, while another putative cholesterol-interacting gene was overexpressed in the trunk tissue and may partly contribute to RNAi import. Altogether, our results present the most comprehensive overview of RNAi pathway effectors in major intermediate snail hosts for trematodes. The findings underscore the likely broad potential for RNAi use in trematode intermediate hosts as an experimental tool and potential control method.

## Introduction

Pulmonate freshwater snails (Gastropoda: Mollusca) play significant roles in the transmission of parasitic helminths, most notably flukes (Trematoda: Platyhelminthes) of global medical and veterinary importance (Lu et al., 2018; Skala et al., 2020). They serve as intermediate hosts supporting the asexual multiplication cycle of the parasites. Given the significance of the diseases they transmit and their importance as a model in biological research, *Biomphalaria*, *Bulinus*, and *Lymnaea* species are among the most well studied trematode-transmitting snails.

While functional genomics elucidates the associations between the genetic makeup and phenotypic traits of an organism, RNA interference (RNAi) has been a powerful tool in this process. It silences the expression of specific genes to study their functions through observable loss-of-function mutations (Fire et al., 1998). Although RNAi is a natural gene regulatory system that is evolutionarily conserved among most eukaryotes, its mechanisms can be triggered artificially to suppress endogenous expression of target genes through in vivo or in vitro delivery of synthetic long double-stranded RNAs (dsRNAs, typically ≥ 100 nucleotides in length), short interfering RNAs (siRNAs, ∼ 22 nt), or microRNAs (miRNAs, ∼ 22 nt) that are complementary to the target gene sequence. When introduced into the cell, long dsRNAs are first processed into siRNA duplexes by Dicer (DCR) enzymes. The small silencing RNAs (referred here as siRNAs and miRNAs) within the cell are then loaded into the RNA interference silencing complex (RISC), where they guide an Argonaute (AGO) protein to recognise and bind target mRNA. This binding induces cleavage via the siRNA pathway or translational repression via the miRNA pathway to directly target mRNA. Both pathways ultimately cause degradation of the target mRNA (Wilson and Doudna, 2013).

RNAi has permitted the study of gene functions in many genetically intractable systems, including intermediate snail hosts of helminths (Xu et al., 2023; Baron et al., 2013; Allan et al., 2017; Smith et al., 2021). Methodologically advanced applications of RNAi also demonstrate high potential as a biocontrol tool for insect pests and aquacultural diseases (Charoonnart et al., 2023; Ghosh et al., 2018). The efficiency of RNAi, however, varies between organisms due to inter- and intra-specific differences in the essential effector proteins. These instances are evident among various groups of organisms including protozoans (Baum et al., 2009), nematodes (Nuez and Felix, 2012), arachnids (Nganso et al., 2020), and insects (Arraes et al., 2021). Therefore, to exploit the full potential of RNAi technology, species-specific profiling and functional screening of the pathway effector compositions is critical. Currently, in disease-transmitting snails, description of the RNAi effectors is fragmentary (Huang et al., 2021; Queiroz et al., 2017; Formaggioni et al., 2024). While the available studies provided some fundamental insights into the snail RNAi biology, compositional diversity of the snail RNAi effector components has not been captured. Consequently, the limitations of snail RNAi techniques as tools for experimentation or disease control is not yet understood.

To provide basic insight into RNAi biology that will guide future functional genomics in snail vectors, this paper aims to determine the diversity of RNAi effector components and extrapolate the consequences of those components to RNAi competence in selected species of *Biomphalaria* (*B. glabrata*, *B. pfeifferi*), *Bulinus* (*Bu. truncatus*), and *Lymnaea* (*L. stagnalis*) snails, using bioinformatics, computational, and gene expression approaches. While we identified core RNAi components that primarily mediate gene silencing in these snails, the absence of other important effectors known to enhance RNAi efficiency provides new mechanistic understanding of the snail RNAi pathways.

## Materials and Methods

### Curation of query sequences and reciprocal BLAST searches

Known RNAi effectors from the well-studied *Caenorhabditis elegans*, *Drosophila melanogaster*, and *Homo sapiens* models were identified systematically from literature up until April 2024 and grouped based on their functional roles in the RNAi pathway. Other model organisms belonging to Order Rhabditida, Digenea, Coleoptera, Hemiptera, Orthoptera, and Decapoda were included for further validation of the effector orthologues. The comprehensive literature searches were conducted using search engines and scholarly databases including PubMed, Web of Science, Scopus, and Google Scholar, using general search terms such as “RNAi effectors *Drosophila*”, “RNAi effectors *C. elegans*”, “RNAi export *C. elegans*” and more specific terms such as “sid-1 human” with Boolean operators where appropriate. Additional full-text articles were retrieved by manual review of the reference lists of relevant articles.

Query protein sequences of the identified effectors were retrieved from online resources, including WormBase for *C. elegans* and other nematodes (http://www.wormbase.org/) (Sternberg et al., 2024), WormBase ParaSite for platyhelminths (https://parasite.wormbase.org/index.html) (Howe et al., 2017), FlyBase for *D. melanogaster* (https://flybase.org/) (Öztürk-Çolak et al., 2024), UniProt for *H. sapiens* (https://www.uniprot.org/) (The Uniprot Consortium, 2024), and NCBI for other sequences (https://www.ncbi.nlm.nih.gov/protein) (Goldfarb et al., 2025). The protein sequences were used as search strings to identify orthologous RNAi proteins in the snails. Accessions of all query proteins used are provided in the Supplementary Material 1. For the four snail species, genome assembly completeness was analysed using BUSCO (Benchmarking Universal Single-Copy Orthologs) v5.8.0 (Simao et al., 2015) on Galaxy (https://usegalaxy.eu/). For *B. glabrata*, *B. pfeifferi*, and *Bu. truncatus* with annotated genome assemblies available through NCBI GenBank database, reciprocal BLASTp searches were performed against the NCBI non-redundant (nr) protein database v. 2.15.0, with default parameters. In cases where a protein orthologue was confirmed in only one of the snail species, the snail sequence was used as a query for other snail species. Since a publicly available *L. stagnalis* transcriptome was not available at the time of research, a transcriptome was generated in-house by us using previously published data of other authors. The adapter-trimmed, high-quality RNA sequencing (RNA-Seq) reads (SRA Bioproject PRJNA420639) were mapped to an *L. stagnalis* genome (https://www.ncbi.nlm.nih.gov/datasets/genome/GCA_900036025.1/), using HiSAT2 v2.1.0 (Kim et al., 2015), with default settings. Samtools v1.9 was used to convert SAM files to BAM files (Li et al., 2009). New GTFs were merged into a single matrix, and reads which uniquely mapped to a gene were counted using StringTie v1.3.6 (Pertea et al., 2015). The resulting FASTA file was used as a BLAST search database. The RNAi protein queries were searched using tBLASTn against the *L. stagnalis* transcriptome. BLAST searches were performed via the NI-HPC facility (https://www.ni-hpc.ac.uk/). Hits were converted to protein sequence using the Expasy Translate tool (Artimo et al., 2012), followed by reciprocal tBLASTn of the longest open reading frames (ORFs) on NCBI.

### Conserved domain/motif annotation and three-dimensional structure prediction

For further identification of the snail orthologues and to resolve any ambiguity after applying the above parameters, conserved functional domains of selected snail orthologues were assessed by searching the sequences against multiple protein family databases including Pfam, NCBIfam, SMART, CDD, Prosite, and Panther, using *hmmscan* algorithm of the Hidden Markov models (Finn et al., 2015) and InterProScan (Blum et al., 2025), adopting default settings in all cases.

To predict functional role in dsRNA import, in silico characterisation of SID-1 (systemic RNA interference-defective 1)-like orthologues from cholesterol uptake protein 1 (CHUP-1), a SID-1 paralogue in *C. elegans* (Whangbo et al., 2017), was done by first assessing the presence or absence of the two mirroring transmembrane cholesterol-recognition motifs; the CRAC (Cholesterol-Recognition Amino acid Consensus) and CARC (inverse CRAC) motifs, characterised by L/V-X_(1–5)_-Y-X_(1–5)_-R/K and K/R-X_(1–5)_-Y/F-X_(1–5)_-L/V, respectively (Fantini et al., 2016), followed by scrutiny of the N-terminal extracellular microdomains (Whangbo et al., 2017). Topologies of the transmembrane motifs were predicted using DeepTMHMM v1.0.42 (https://dtu.biolib.com/DeepTMHMM/) (Hallgren et al., 2022) and Phobius (https://phobius.sbc.su.se/) (Kall et al., 2007), and were visualised using Protter web server v1.0 (https://protter.ethz.ch/start/) (Omasits et al., 2014).

For DCRs, multidomain analysis was performed by assessing the two RNase III catalytic domains and the tandemly arrayed C-terminal dsRBD (dsRNA-binding domain), the PAZ (Piwi-Argonaute-Zwille) and Platform domains, the N-terminal helicase domain, and the Domain of Unknown Function DUF283 (Dicer dimerisation domain) (Ciechanowska et al., 2021; Zapletal et al., 2023). The two functionally distinct DCR proteins in *D. melanogaster* (*Dm*DCR-1 and *Dm*DCR-2) were included as model, and conservation of the pre-miRNA 5’-end binding pockets within the Platform–PAZ region (Lee et al., 2023; Park et al., 2011) was analysed in the snail orthologues. For AGO orthologues, catalytic triad DDX or tetrad DEDX (where X is either Asp (D) or His (H) amino acid) of the Piwi domain (Nakanishi et al., 2012) was considered for predicting the slicer activity. Staufen (STAU-1/2) proteins were assessed by their characteristic dsRNA-binding domain repeats (RBDs) (Yoon et al., 2018) and the presence/absence of N-terminal signal peptide (Koo and Palli, 2024) was predicted using SignalP 6.0 model (Teufel et al., 2022).

AlphaFold3 Server (Abramson et al., 2024) was used to predict three-dimensional protein fold structures. AlphaFold files were converted to protein data bank (PDB) files with predicted local distance difference test (pLDDT) confidence scores assigned to the B-factor field using Biopython (v1.85) (Cock et al., 2009), before further analysis and visualisation using ChimeraX v1.9 (Meng et al., 2023). Structural regions with very low prediction confidence (plDDT < 50) were trimmed out.

### Multiple sequence alignments and phylogenetic analysis

Alignments of multiple protein sequences were performed using Clustal Omega v1.2.4 (Sievers and Higgins, 2018) and visualised in Jalview v2.11.4.1 (Procter et al., 2021). WebLogo v3.7 (Crooks et al., 2004) was used to generate graphical representations of patterns within some multiple sequence alignments. Phylogenetic trees were constructed using MEGA11 software v11.0.13 (Tamura et al., 2021). Multiple full-length or partial protein sequences were subjected to the plug-in MUSCLE sequence alignment programme and evolutionary relationships were inferred using Maximum Likelihood (ML) clustering with 1,000 non-parametric bootstrap replicates. The non-bilaterian *Nematostella vectensis* (sea anemone) and *Amphimedon queenslandica* (sponge) were used as outgroups. Best-fitting substitution models for ML clustering were selected based on lowest Bayesian Information Criterion (BIC) and corrected Akaike Information Criterion (AICc) scores, while also considering sequence divergence and composition across datasets. Bootstrap consensus trees were visualised and edited with FigTree v1.4.4 (http://tree.bio.ed.ac.uk/).

### Protein–protein interaction (PPI) network and gene ontology (GO) analysis

Putative *B. glabrata* RNAi effector protein sequences were mapped to STRING v12.0 database (Szklarczyk et al., 2025) to generate interaction networks and GO terms. PPI networks were built with 70% confidence and clustered using k-means algorithm. Clustered networks were loaded into Cytoscape v3.10.3 (Shannon et al., 2003) for further annotation, analysis, and visualisation. Gene ontology (GO) enrichments for biological process (BP), cellular component (CC), and molecular function (MF) were obtained with enrichment false discovery rate (FDR) ≤ 0.05 and *P*-values corrected for multiple testing for each category using the Benjamini–Hochberg procedure (Benjamini and Hochberg, 1995).

### Gene expression analysis

#### Snail husbandry

*Biomphalaria glabrata* snails (uninfected NMRI strains) were used in this study. The snails were separated from the main aquarium and reared in aerated, transparent 1L plastic containers filled with diluted 1x artificial river water (10x ARW contained 5L H_2_O, 2.78g CaCl_2_, 6.15g MgSO_4_.7H_2_O, 2.1g NaHCO_3_, 215mg K_2_SO_4_ and 250µL FeCl_3_.6H_2_O, at pH 7.0) and kept at 25 _, under 12:12 light/dark photoperiod. The snails were fed tropical fish food flakes (Pets at Home, UK). Both the rearing water and the food were changed three times weekly.

#### Experimental design

Gene transcripts of six putative RNAi effectors in *B. glabrata* (*Ago2*, *Dcr1*, *Eri1*, *Sid1L*, *Chc1*, and *Stau*) were selected for mRNA expression evaluation, based on their primary relevance in the RNAi machinery. Expression levels of some key effectors such as Dicers and Argonautes may vary significantly with the age of the organism (Sanz-Ros et al., 2023). To examine whether snail size influences expression levels of three major genes, *Ago2*, *Dcr1*, and *Eri1*, whole snail tissues of juveniles and adult snails, which were further categorised into four subgroups based on shell size (diameter), were analysed. The first juvenile group (J1) were 3-4 mm in size, the second juvenile group (J2) were 6-7 mm in size, the first adult group (A1) were 10-11 mm in size, and the second adult group (A2) were 13 mm in size. Each group had five biological replicates. In another assay, gene expression profile of *Sid1L*, *Chc1*, and *Stau* genes were analysed in various isolated tissues rather than whole soft body. Five adult snails (13 mm) were sacrificed and anatomically dissected to isolate the head-foot region (HF), ovotestis (OV), hepatopancreas (digestive gland) (HP/DG), and albumen gland (AG) from each snail. The remaining tissue, here termed ‘trunk’ T (consisting of the mantle lobe, lung, pseudobranch, intestine, stomach, heart, and kidney) was also assessed for gene expression (Figure 8A).

#### RNA isolation, cDNA synthesis, and quantitative PCR

Total RNA was extracted from snail tissues using TRIzol (Invitrogen), followed by DNase treatment using DNA-*free*™ kit (Invitrogen). RNA samples were quantified spectrophotometrically before and after DNase treatment, using a DeNovix DS-11 FX instrument. First-strand cDNA was synthesised by reverse transcription (RT) of 1.8 µg of each DNase-treated RNA sample using High-Capacity RNA-to-cDNA™ Kit (Applied Biosystems). The reverse-transcriptase PCR (RT-PCR) conditions were 37 _ for 1 hour, 95 _ for 5 min, and 4 _ until storage. For quantitative PCR (qPCR), oligonucleotide primers (Metabion) were designed using Primer-BLAST (https://www.ncbi.nlm.nih.gov/tools/primer-blast/) and Primer3Plus (https://www.primer3plus.com/). Supplementary Material 1 contains the list of all primer sequences used. All primer sets were tested on endpoint PCR with 1.5 µL of cDNA template and electrophoresed on 2% agarose gel in 1x Tris-acetate-EDTA (TAE) buffer to confirm amplicon size and evaluate amplification specificity. Gene expression levels in snail cDNA samples were subsequently assessed real-time on qPCR. Each qPCR reaction was set up in three technical replicates, each having a total reaction volume of 10 µL, containing 1.5 µL of cDNA, 5.0 µL of SensiFAST SYBR No-ROX mix 2x (Meridian Bioscience), 0.5 µL of the F primer, 0.5 µL of R primer, and 2.5 µL of molecular grade water. Negative no-template controls were included for each sample by replacing cDNA with molecular grade water. qPCR was run on a Qiagen Rotor-Gene Q instrument by 40 cycles of 95 _ for 10 sec, 60 _ for 15 sec and 72 _ for 20 sec, after an initial hold for 10 min at 95 _. Thermal cycling results were processed on Rotor-Gene Q series software v2.3.1. *Biomphalaria glabrata* β-Actin gene was selected as endogenous control for normalisation, as it has shown stable expression in *B. glabrata* of different ages and sizes (Xu et al., 2023). Relative gene expression was calculated using Pfaffl’s *E*^-ΔCt^ method (Pfaffl, 2001).

#### qPCR data analysis and visualisation

All qPCR data were statistically analysed, compared among groups, and visualised using R software v4.3.2 (R Core Team, 2025). Data that assumed normal distribution (Shapiro–Wilk’s *P* > 0.05) were subjected to one-way ANOVA for multiple comparisons, followed by Tukey’s HSD test. Otherwise, non-parametric Kruskal–Wallis test was performed for multiple comparisons among groups, and Wilcoxon Rank Sum (Mann–Whitney U) test for comparison between two independent variables. Non-parametric multiple comparisons were followed up by Dunn’s *post-hoc* test with Bonferroni correction. In all cases, *P* values ≤ 0.05 were considered statistically significant.

## Results

### Comparative BUSCO analyses showed uneven genome completeness between snail species

Based on BUSCO estimates, *B. glabrata*, *B. pfeifferi*, and *Bu. truncatus* genomes showed highly complete genomes, although higher gene duplication was found in *Bu. truncatus* (Table 1). Under the same parameters, the *L. staginalis* genome showed much lower BUSCO scores for completeness. Relative to other genomes, many more pan-eukaryotic genes may be fragmented or missing in the *L. staginalis* genome (Table 1).

**Table 1.** Comparative genome completeness estimates for the snail species based on BUSCO analysis.

| Species | GenBank | C (%) | S (%) | D (%) | F (%) | M (%) | *n* | BUSCO<br>(Benchmarking Universal Single-Copy Orthologs) |
| --- | --- | --- | --- | --- | --- | --- | --- | --- |
| *Biomphalaria glabrata* | GCA_947242115.1 | 96.9 | 95.7 | 1.2 | 2.4 | 0.8 | 255 |  |
| *Biomphalaria pfeifferi* | GCA_030265305.1 | 96.1 | 95.7 | 0.4 | 2.7 | 1.2 | 255 |  |
| *Bulinus truncatus* | GCA_021962125.1 | 96.5 | 67.5 | 29.0 | 2.7 | 0.8 | 255 |  |
| *Lymnaea stagnalis* | GCA_900036025.1 | 73.3 | 71.8 | 1.6 | 21.2 | 5.5 | 255 |  |

categories were reported as: C (Complete), subdivided into S (Complete and single-copy) and D (Complete and duplicated); F (Fragmented); and M (Missing). *n* indicates the total number of BUSCO groups searched. All assemblies were evaluated under the same parameters (eukaryota\_odb10 lineage, euk\_genome\_met mode, and metaeuk gene predictor) to ensure comparability.

### Snail vector genomes encode RNAi pathway components

A total of 115 RNAi effector proteins were collated, comprising 47 uptake and transport proteins, 11 DCR and DCR-associated proteins, 4 AGOs (excluding worm-specific AGOs), 11 RISC proteins, 7 RNAi amplification effectors, 15 RNAi inhibitors, and 20 nuclear RNAi effectors (Table 2). Out of these, 83 orthologues were found in *L. stagnalis*, 76 in *B. glabrata*, 75 in *Bu. truncatus*, and 74 in *B. pfeifferi* (Table 2, Figure 2). Each snail species possesses at least one orthologue in each functional group of the RNAi effectors (Table 2). Nevertheless, all the snails examined lack orthologues of the RNA-dependent RNA polymerases (RdRPs) (RRF-1 and EGO-1) and associated proteins (RDE-10 and RDE-11) (Figure 2), which are key players in secondary RNAi amplification. Similarly, core components responsible for heritable or transgenerational RNAi in *C. elegans* (the nuclear RNAi defective NRDE-1, NRDE-2, NRDE-3 and NRDE-4) were absent from the snails (Figure 2). While these absences may limit RNAi potency, other preserved effector components are sufficient for successful RNAi.

**Figure 1.**
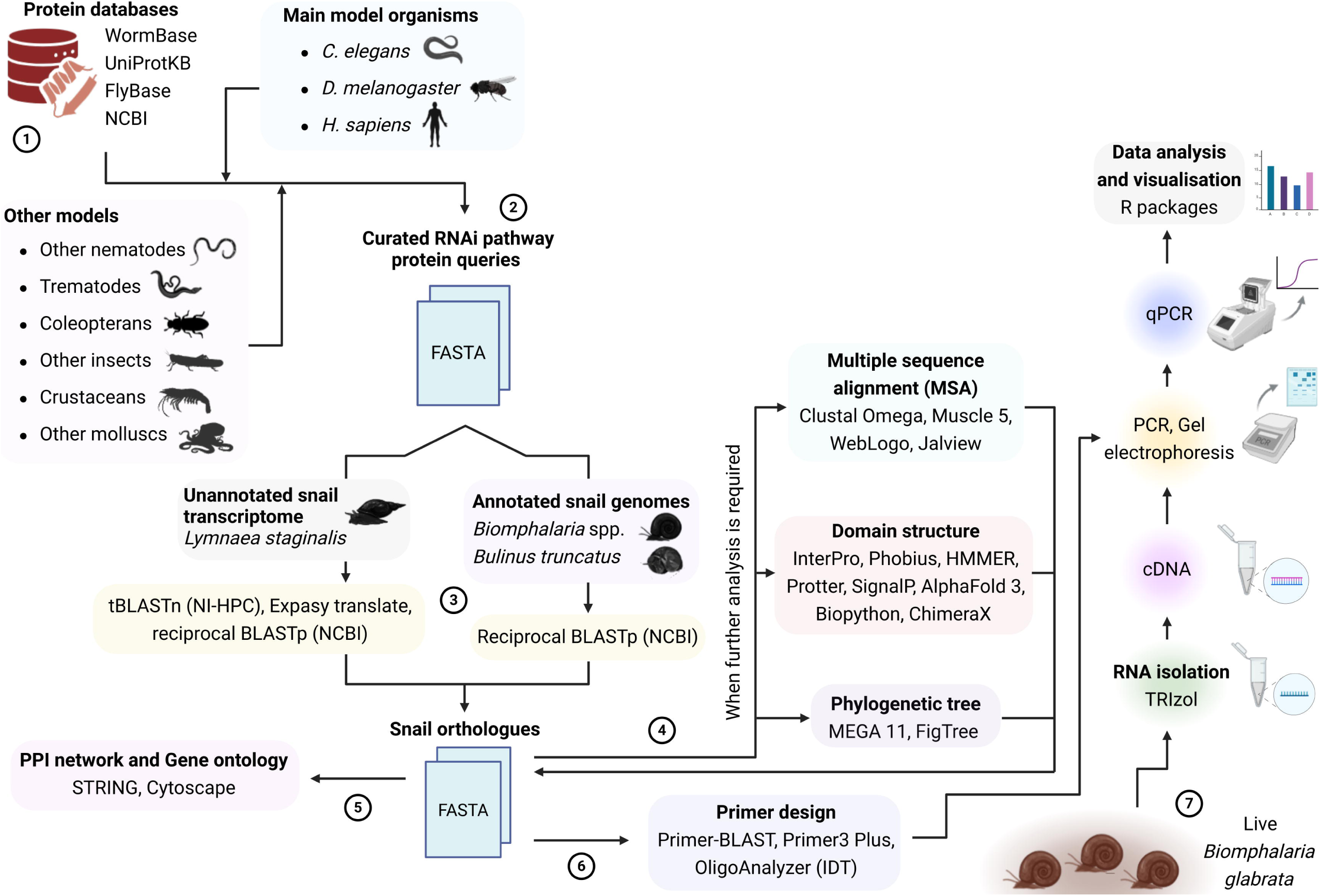
Methodological overview of the identification and gene expression analysis of RNAi effector components in snail vectors of parasitic flukes.

**Figure 2.**
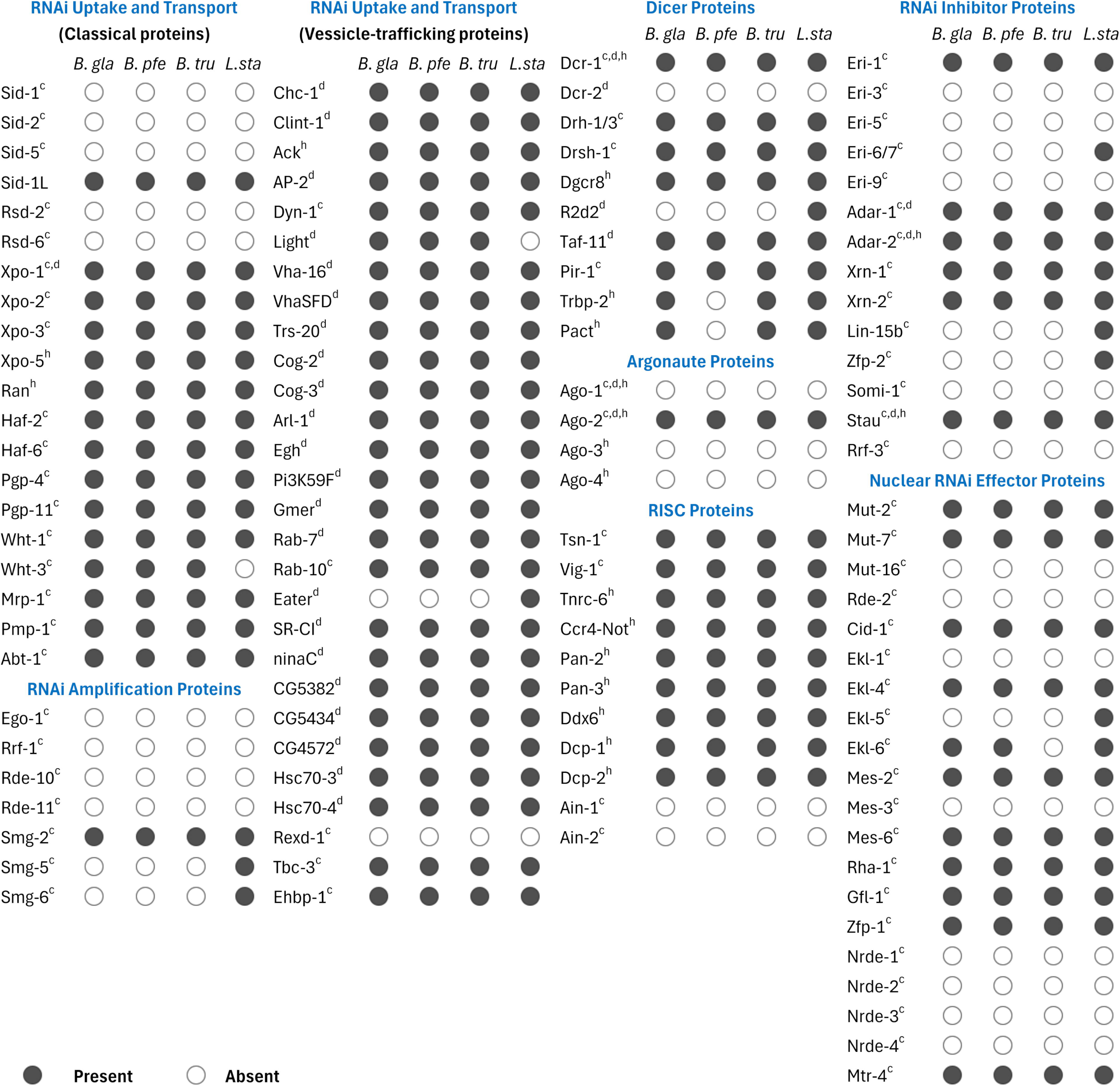
A compendium of predicted RNAi effector components in *Biomphalaria* (*B. gla, B. pfe*), *Bulinus* (*B. tru*), and *Lymnaea* (*L. sta*) snails. Query sequences used were retrieved from *C. elegans* (^c^), *D. melanogaster* (^d^), or *H. sapiens* (^h^). In *Biomphalaria* spp., WHT-1 and WHT-3 protein orthologues are encoded by a single gene. Human TRBP-2 and PACT shared single orthologous proteins in *B. glabrata* and *Bu. truncatus*. In *L. staginalis*, Drh-1 and Drh-3 returned separate sequences. Full accessions and sequences are available in the Supplementary Material 1.

**Table 2.** Functional grouping and diversity of snail RNAi effector proteins.

| Effector protein category | No. queries | *B. glabrata* | *B. pfeifferi* | *Bu. truncatus* | *L. stagnalis* |
| --- | --- | --- | --- | --- | --- |
| RNAi uptake and transport | 47 | 40 | 39 | 40 | 40 |
| Dicer and DCR-associated | 11 | 8 | 6 | 8 | 9 |
| Argonautes (AGOs) | 4 | 1 | 1 | 1 | 1 |
| RISC | 11 | 9 | 9 | 9 | 9 |
| RNAi amplification | 7 | 1 | 1 | 1 | 3 |
| Nuclear RNAi | 20 | 11 | 11 | 10 | 12 |
| RNAi inhibition | 15 | 6 | 6 | 6 | 9 |
| Total | 115 | 76 | 74 | 75 | 83 |

### RNAi uptake and spreading may be mediated by a vesicle-trafficking pathway, rather than the transmembrane SID-dependent pathway

Reciprocal BLASTp screens indicated that snail SID-1-like sequences displayed higher sequence similarities with *C. elegans* CHUP-1, and human SID-1-like orthologue (*Hs*SIDT-1) (Figure 3A). Although *B. truncatus* SID-1 shared slightly higher percent sequence identity with the *C. elegans* SID-1 (*Ce*SID-1) orthologue (Figure 3A), best match (higher bit score of 169 with 81% query cover and a lower E-value of 2e-47) was found between *B. truncatus* SID-1-like orthologue and *C. elegans* CHUP-1. Further transmembrane sequence analysis identified six putative cholesterol-binding CRAC motifs in each of the two *Biomphalaria* species, and three CRAC motifs in *Bu. truncatus* and *L. stagnalis* (Figure 3B; Table 3). Each snail sequence also possessed multiple mirrored transmembrane CARC motifs (Table 3). Conserved CRAC and CARC motifs were also found across other molluscan SID-1-like proteins, including slugs, limpets, bivalves, and cephalopods (Table 3). Interestingly, single CRAC (transmembrane 1, TM1) and CARC (TM9) motifs were found within the *Ce*SID-1 sequence (Figure 3C; Table 3), suggesting that the structural framework for cholesterol binding is conserved in both SID-1 and CHUP-1 proteins. However, the characteristic N-terminal microdomains of *Ce*SID-1 were not observed in any of the molluscan or non-molluscan SID-1-like orthologues examined (Supplementary Material 1). Similarly to SID-1, orthologues of other *C. elegans* dsRNA-transporting SID proteins (SID-2 and SID-5) appeared absent from snail genomes (Figure 2). These results suggest that the canonical SID-dependent RNAi transport pathway, well characterised in *C. elegans*, may be absent from gastropod lineages.

**Figure 3.**
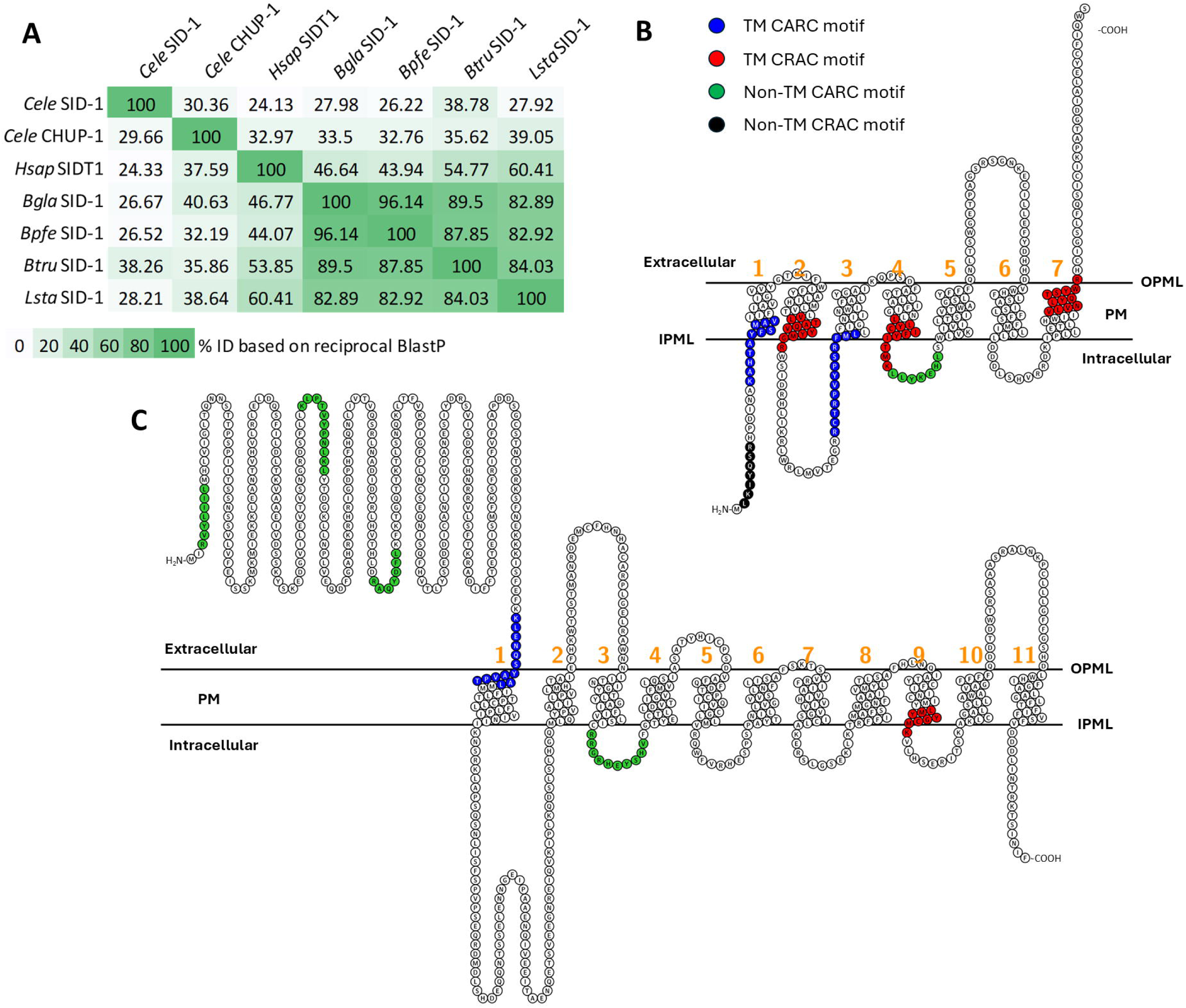
Characterisation of snail SID-1-like proteins **A.** Heatmap matrix for reciprocal BlastP showing percent identity between orthologues. **B.** Predicted topology of *Bu. truncatus* SID-1 showing the TM and non-TM CRAC/CARC motifs. **C.** Predicted topology of *C. elegans* SID-1 secondary structure showing single TM CRAC and CARC motifs. The first 19 amino acid residues at the N-terminus of *C. elegans* SID-1 protein (M1–N19) were predicted to constitute a signal peptide domain. In figures B and C, numeric annotations in orange colour indicate number of TM domains. CRAC, cholesterol-recognition amino acid consensus; CARC, inverse CRAC motif; IPML, inner plasma membrane layer; OPML, outer plasma membrane layer; PM, plasma membrane; TM, transmembrane.

**Table 3.** Amino acid residues of the putative transmembrane CRAC and CARC motifs in selected SID/CHUP-1-like sequences.

| Protein | Transmembrane CRAC motifs | Transmembrane CARC motifs |
| --- | --- | --- |
| *Cele*SID-1 | L674-K681 | K308-L320 |
| *Cele*CHUP-1 | L440-R448 | K265-V274, K370-L378, K636-L643 |
| *Bgla*SID-1 | L528-R534, L577-R585, L651-R660, V707-R717, V761-K769, L786-K798 | K505-L514, K667-V677, K738-L750 |
| *Bpfe*SID-1 | L477-R483, L526-R534, L600-R609, L656-R666, V709-K717, L735-K745 | K454-L463, K616-V626, K687-L699 |
| *Btru*SID-1 | V55-R65, L134-K144, V247-R258 | K15-V25, R86-L98 |
| *Lsta*SID-1 | L61-R70, V117-R127, L196-K206 | K77-V87, R148-L160 |
| *Acal*SID-1 | L579-K585, L628-R636, L702-R711, L759-R768, L837-K847 | K556-L565, K718-V728, R789-V800 |
| *Cvir*SID-1 | L421-K427, L468-R476, L543-R552, L560-R569 | K273-V279, R398-L407, K559-V570, R630-L642 |
| *Lgig*SID-1 | L56-K62, L173-R182, L309-K318 | K33-L42, R260-L272 |
| *Mgal*SID-1 | L488-K494, L610-R619 | K265-V271, K465-L474, K626-V637 |
| *Ovul*SID-1 | L517-K523, L639-R648, V695-R705, L774-K784 | K494-L503, K729-L738 |
| *Hsap*SIDT-1 | L584-R593, L643-R652, L722-K732 | K302-L311, K540-L552, R677-L686, R707-L714 |

CRAC, cholesterol-recognition amino acid consensus; CARC, inverse CRAC motif; K, lysine; L, leucine; R, arginine; V, valine; *Cele*, *Caenorhabditis elegans* (nematode); *Bgla*, *Biomphalaria glabrata* (gastropod), *Bpfe*, *Biomphalaria pfeifferi* (gastropod); *Btru*, *Bulinus truncatus* (gastropod); *Lsta*, *Lymnaea stagnalis* (gastropod); *Acal*, *Aplysia californica* (gastropod slug); *Cvir*, *Crassostrea virginica* (bivalve); *Lgig*, *Lottia gigantea* (limpet); *Mgal*, *Mytilus galloprovincialis* (bivalve); *Ovul*, *Octopus vulgaris* (cephalopod); *Hsap*, *Homo sapiens* (mammal). Supplementary Material 1 contains the multiple sequence alignments of the transmembrane CRAC/CARC motifs and the accession numbers of all protein sequences.

Protein components of an alternative, SID-independent, RNAi-importing clathrin-mediated endocytosis (CME) pathway were expectedly preserved in the intermediate host snails (Figure 2). These was reflected in clathrin heavy chain 1 (CHC-1) and associated proteins, such as clathrin adapter protein AP-50/AP-2, clathrin interactor CLINT-1 (the mammalian ENTH (epsin N-terminal homology)-domain-containing orthologue of *C. elegans* RSD-3), dynamin-1 (DYN-1), vacuolar ATPase VHA-16, VhaSFD (a VHA-16 subunit), RAB-7 (a Ras-like GTPase), ARF72A/ARL-1 (an ADP-ribosylation factor), a non-receptor tyrosine kinase ACK (the mammalian orthologue of *C. elegans* SID-3), scavenger receptors (Sr-CI and Eater), and protein chaperones (HSC70-3 and HSC70-4). Snail orthologues corresponding to vesicle proteins that facilitate cellular export of silencing RNAs, REXD-1, TBC-3, EHBP-1 and RAB-10, were also present (Figure 2). These results suggest that the import of exogenous silencing RNAs and their systemic spreading in vector snails may be mediated majorly via clathrin-dependent, vesicle-trafficking pathway given the absence of SID-1-like sequences in our datasets.

### Snail Dicer (DCR) and Argonaute (AGO) proteins contain functional domains and amino acid residues for miRNA/siRNA biogenesis and processing

Sequences similar to DCR and AGO proteins from humans, *D. melanogaster*, and *C. elegans* were identified in the snails (Figure 1). Single orthologues of DCR proteins were found in all snails, except *Bu. truncatus* that returned two putative hits (Figure 4A). Similarly, two putative AGO sequence hits were found in *B. glabrata* (Figure 5A).

**Figure 4.**
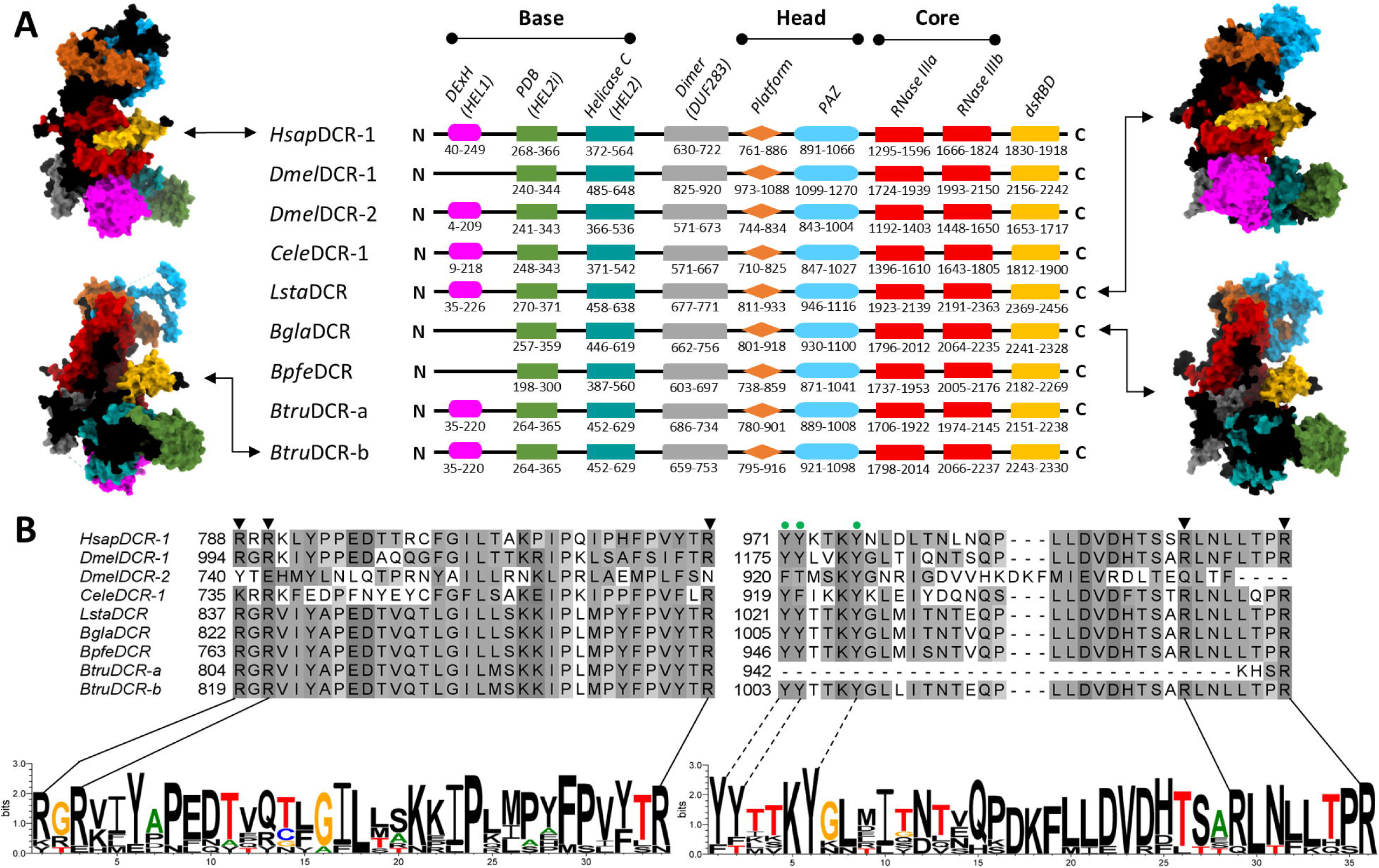
Comparative domain structures of Dicer in *H. sapiens*, *C. elegans*, *D. melanogaster*, and the studied snail species. **A.** Cartoon representation of the domains with their amino acid sequence ranges. The ‘head’ consists of the Platform-PAZ domain, the ‘base’ is made up of the helicase domain, while the ribonulcease domain forms the ‘core’. InterProScan and Pfam accessions: **DExH**, DEAD box helicase domain (IPR011545/PF00270); **PBD**, Dicer partner-binding domain (IPR048513/PF20930); **Helicase C**, helicase conserved C-terminal domain (IPR001650/PF00271); **Dimer**, Dicer dimerisation domain (IPR005034/PF03368); **PAZ** (Piwi-Argonaute-Zwille) domain (IPR003100/PF02170); **RNase III**, Ribonuclease III (IPR000999/PF00636); **dsRBD**, dsRNA-binding domain (IPR014720/PF20932). AlphaFold three-dimensional (3D) protein folding architectures, showing the domains in same colour with cartoon representation, for *H. sapiens* (model), *B. gblabrata*, *Bu. truncatus*, and *L. stagnalis* Dicers are presented on the either side. **B.** Hypothetical motifs for 5’-end recognition pockets in the Platform–PAZ region of the studied snail species. *Bgla*DCR (R822, R824, R855, R1030 and R1037); *Bpfe*DCR (R763, R765, R796, R971 and R978); *Btru*DCR-a (R804, R806, R837 and R945); *Btru*DCR-b (R819, R821, R852, R1028 and R1035); *Lsta*DCR (R837, R839, R870, R1049 and R1056). Conserved tyrosine (Y) residues of the 3’-end pockets (Lee et al., 2023) within the captured peptides are highlighted in dashed lines.

**Figure 5.**
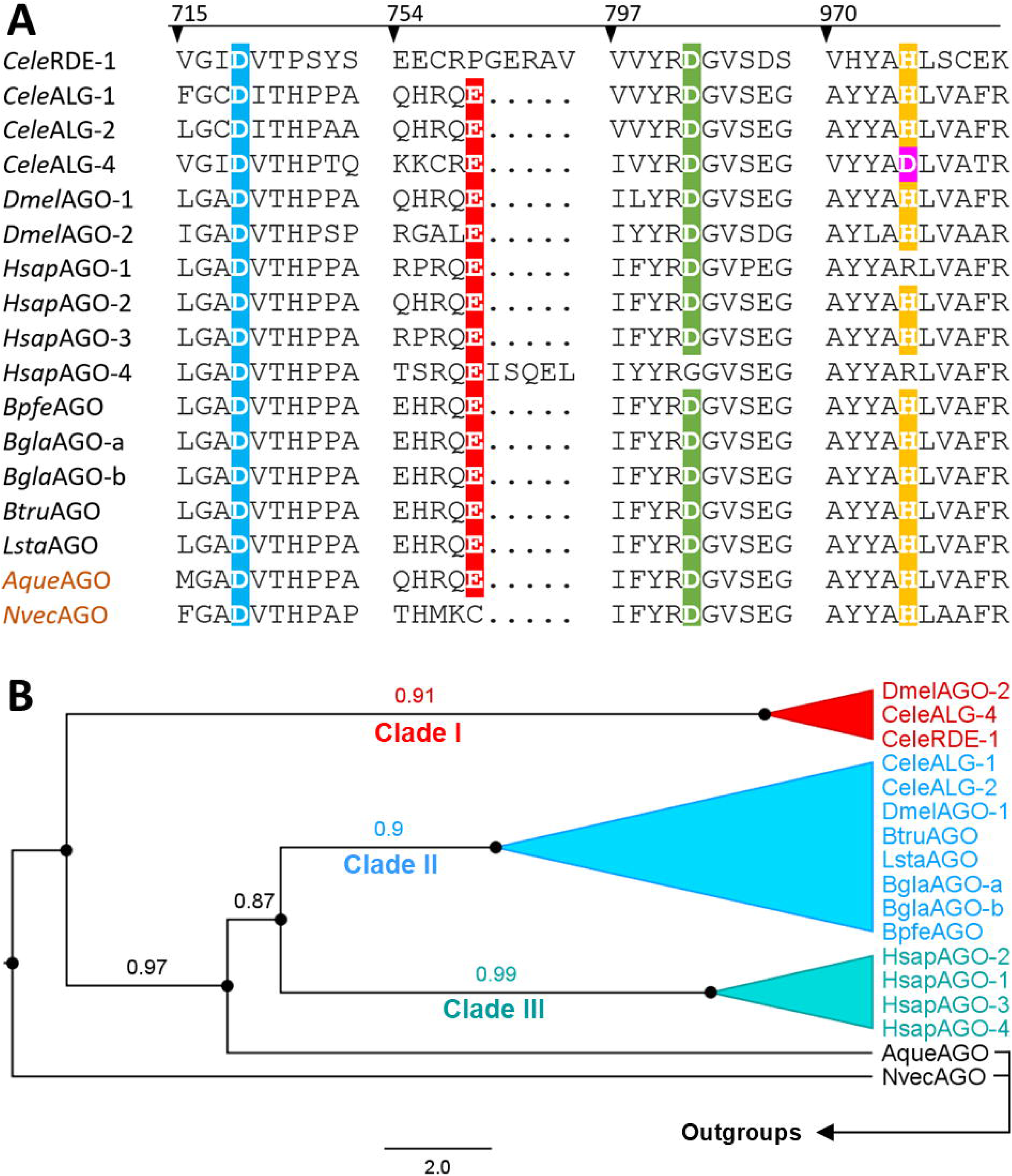
Argonaute (AGO) proteins in intermediate host snails **A**. Multiple alignments of the Piwi domain sequences highlighting conserved catalytic residues. **B.** Maximum likelihood phylogeny reconstructed from the PAZ/Mid/Piwi regions. Sequences represent freshwater snails *Biomphalaria glabrata* (*Bgla*AGO), *Biomphalaria pfeifferi* (*Bpfe*AGO), *Bulinus truncatus* (*Btru*AGO), and *Lymnaea stagnalis* (*Lsta*AGO); *D. melanogaster* (*Dmel*AGO), *C. elegans* (*Cele*AGO), *H. sapiens* (*Hsap*AGO), and two selected outgroups (*Amphimedon queenslandica* and *Nematostella vectensis*). Outgroups were used to root the tree, enabling polarisation of bilaterian AGO diversification relative to early-branching metazoans. Alignment positions were filtered using partial deletion at 80% site coverage and phylogenetic inference was performed under the LG+G model. Branch labels represent bootstrap support values ranging between 0 and 1. Values closer to 1 denote higher confidence in the branching relationship. This analysis involved 17 amino acid sequences and there was a total of 582 positions in the final dataset. Scale bar indicates the expected number of amino-acid substitutions per site.

The functional divergence of DCR enzymes is well established in *D. melanogaster*, where *Dm*DCR-1 and *Dm*DCR-2 specialise in miRNA and siRNA biogenesis, respectively, through distinct substrate preferences and ATP requirements. Guided by these known structural features, we examined the architecture of snail DCR orthologues. Our analysis revealed that the overall domain organisation of the snail sequences closely resembles that of *Hs*DCR (mi/siRNA), *Ce*DCR (mi/siRNA), and *Dm*DCR-2 (siRNA only). Notably, in *B. glabrata* and *B. pfeifferi*, the DExH (HEL1) lobe was absent from the helicase N-terminus, a configuration reminiscent of ATP-independent *Dm*DCR-1 (miRNA only) (Figure 4A). Further inspection of the Platform–PAZ region, however, demonstrated perfect conservation of both pre-siRNA 3’-end and pre-miRNA 5’-end recognition motifs across all snail orthologues (Figure 4B).

At the amino sequence level, all snail AGO sequences shared higher similarities with *D. melanogaster* AGO-1/2 isoforms. Among the four human AGO isoforms (*Hs*AGO 1-4), the snail orthologues also showed higher reciprocal similarities with *Hs*AGO-2. When the Piwi domains were further examined, all snail AGO sequences encoded the conserved DEDH catalytic motif (Figure 5A), supporting their potential to mediate siRNA-guided mRNA cleavage activity. Maximum likelihood (ML) phylogeny constructed using human, *D. melanogaster*, *C. elegans*, and snail sequences showed three distinct clades, grouping snail AGOs, *D. melanogaster* AGO-1 (*Dm*AGO-1), and *C. elegans* ALG-1 and ALG-2 within the same clade (Clade II) (Figure 5B).

### Putative RNAi inhibitor proteins, including Staufen, are well conserved in the snail vectors

Six orthologues of known RNAi inhibitors were encoded in *Biomphalaria* and *Bulinus* genomes. In *L. stagnalis*, nine putative RNAi inhibitors were respectively present (Table 2). Out of these, six putative RNAi inhibitors: Adenosine deaminases ADAR-1 and ADAR-2, enhanced RNAi protein ERI-1, exoribonucleases XRN-1 and XRN-2, and Staufen STAU-1/2, were found across the snail datasets (Figure 2). Further domain characterisation of the STAU proteins revealed that an N-terminal signal peptide is preserved only in coleopteran STAU orthologues (StauC) (*Leptinotarsa decemlineata* M1-G21, *Pr* = 0.53; *Tribolium castaneum* M1-G16, *Pr* = 0.79; *Diabrotica virgifera virgifera* M1-S19, *Pr* = 0.98). Conversely, an RBD4 domain found within other STAU sequences was absent (Figure 6A). Phylogenetic ML analysis similarly showed distinct branching of StauC sequences (bootstrap value = 1) from other STAU sequences, which were also clustered by species (Figure 6B).

**Figure 6.**
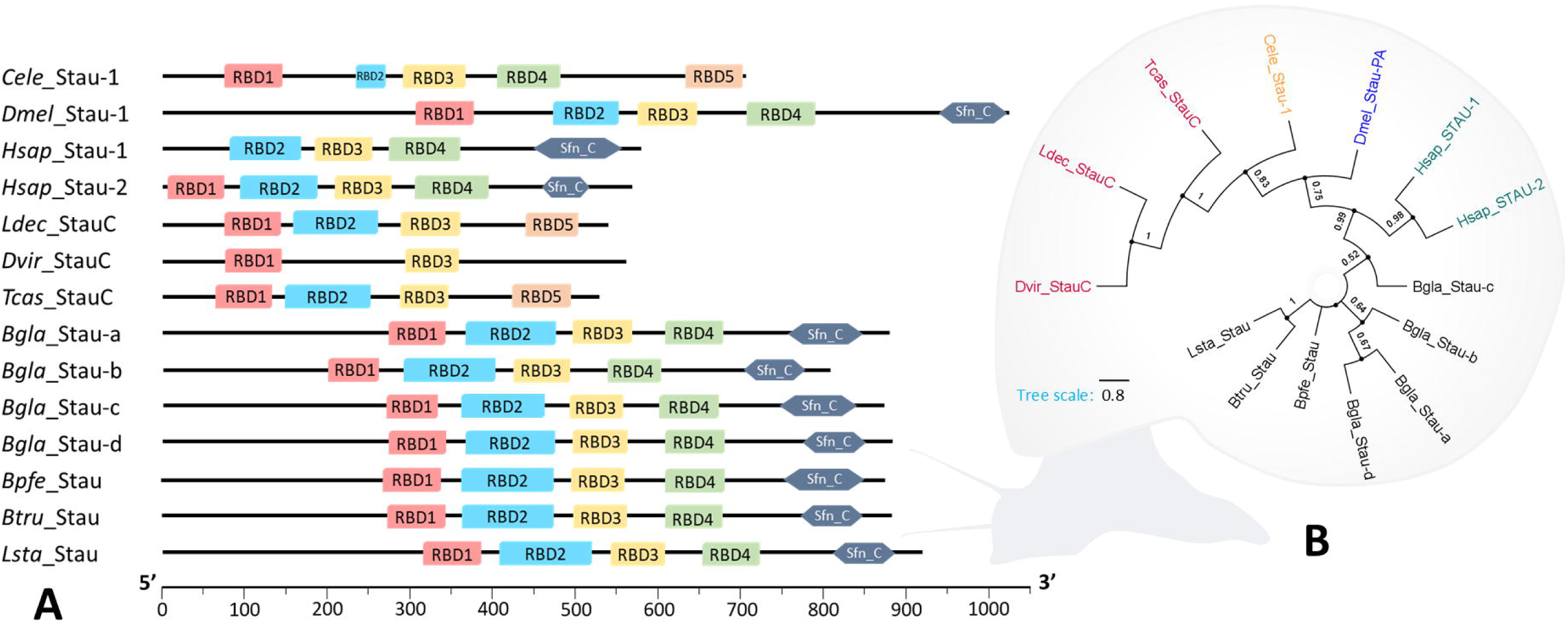
Characterisation of Staufen (Stau) proteins in intermediate host snails of helminths. **A.** Domain profile of Staufen proteins among human (Hsap), coleopteran insects (*Dvir*, *Ldec*, *Tcas*), *Caenorhabditis elegans* (*Cele*), *Drosophila melanogaster* (*Dmel*), and snail vectors (*Bgla*, *Bpfe*, *Btru*, *Lsta*). Domain annotations and coordinates were predicted based on multiple sequence alignments and InterProScan searches and were plotted to scale. **B.** A maximum likelihood cladogram constructed based on full-length sequences of Stau proteins from all species. Tree was inferred using LG+G model and all positions with less than 80% site coverage were eliminated. Bootstrap values lower than 50% were hidden from the tree. This analysis involved 14 amino acid sequences and there was a total of 437 positions in the final dataset. Bgla, *Biomphalaria glabrata*; Bpfe, *Biomphalaria pfeifferi*; Btru, *Bulinus truncatus*; Dvir, *Diabrotica virgifera vergifera*; Hsap, *Homo sapiens*; Ldec, *Leptinotarsa decemlineata*; Lsta, *Lymnaea stagnalis*; Tcas, *Tribolium castaneum*. Hsap_Stau-1 (577aa isoform) and Hsap_Stau-2 (570aa isoform) were used.

### PPI network and GO enrichments supported functional snail orthologues

A protein–protein interaction (PPI) network constructed using *B. glabrata* protein sequences resulted in 15 clusters (Figure 7A) and showed enrichment *P*-value < 1.0e-16. Clathrin-1 (CHC-1) showed biological interactions with AP-2, DYN-1, and CLINT-1. Interactions between DCR-1 and TRBP-2, DRSH-1 and DGCR8, RAN and exportins (XPOs), as well as AGO-2 and deadenylation complexes and decapping enzymes (e.g. TNRC6 and DDX6, respectively) were also represented (Figure 7A). Gene ontology (GO) enrichments also showed strong involvement of the snail protein orthologues in the biological process of gene silencing by RNA (*signal* = 4.29, *FDR* = 9.83e-17), with marked molecular functions in dsRNA binding (*signal* = 3.29, *FDR* = 3.55e-12) and ribonuclease activity (*signal* = 1.82, *FDR* = 1.31e-07). In the cellular components, putative snail proteins associated with the RISC complex (*signal* = 3.14, *FDR* = 9.90e-11) and the processing bodies (P-bodies; *signal* = 2.53, *FDR* = 8.51e-10) showed the strongest signals (Figure 7C, Supplementary Material 2). Six protein orthologues were predicted to constitute the RISC complex (DCR-1, AGO-2b, TSN-1, TRBP-2, XPO-5, and DCP-2), while eight P-body protein orthologues (DDX6, AGO-2b, TNRC6, PAN-2, PAN-3, CCR4-NOT, XRN-1, and DCP-2) were predicted (Figures 7B and 7C, Supplementary Material 2).

**Figure 7.**
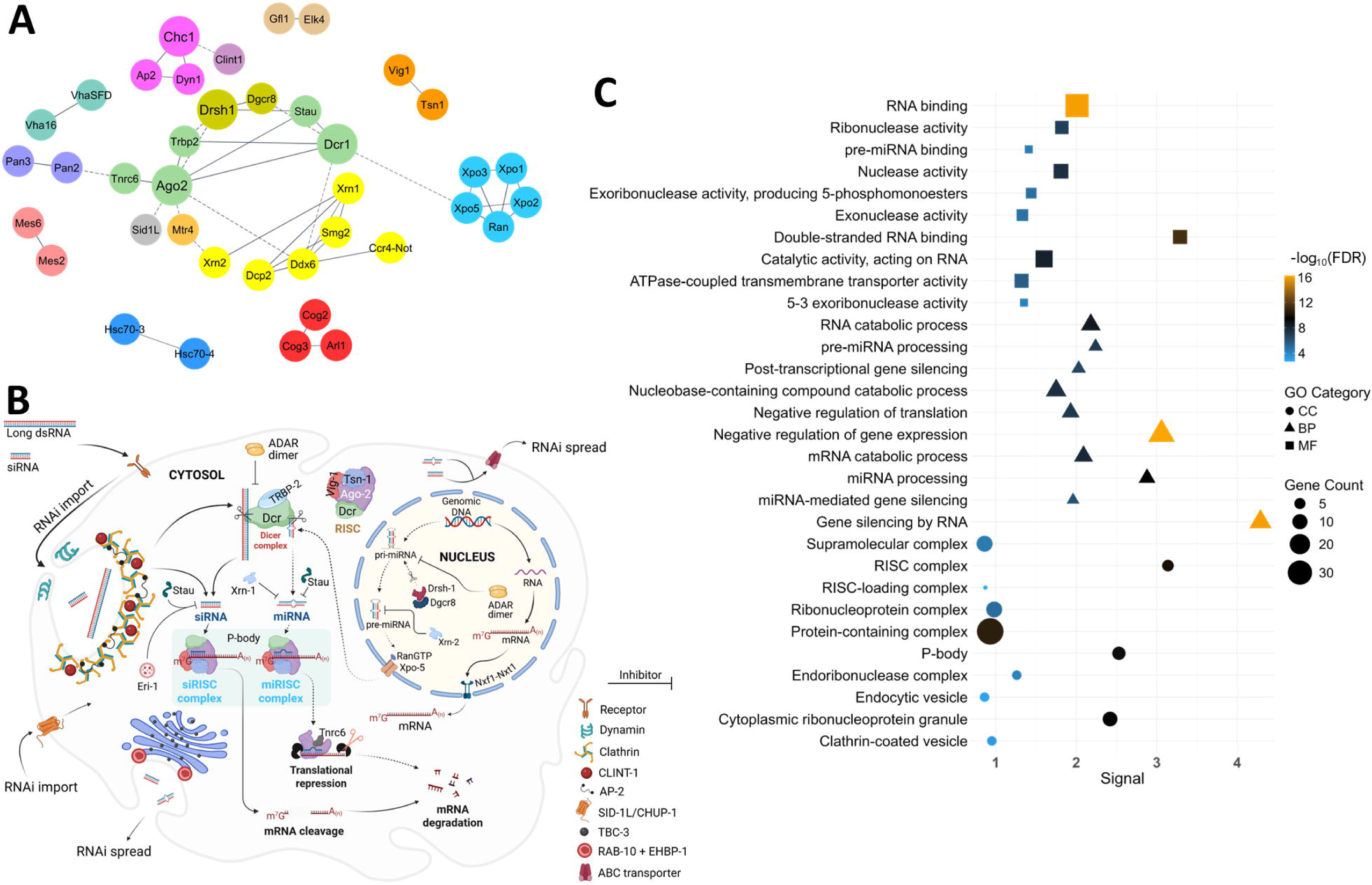
Protein–protein interaction (PPI) and gene ontology (GO) analyses **A.** PPI network of putative RNAi complements in *Biomphalaria glabrata*. Only connected nodes are shown. Straight unbroken edges between nodes indicate direct functional interactions while dashed edges indicate indirect functional associations. **B.** Hypothetical model for mechanistic RNAi pathways in a single cell of *Biomphalaria or Bulinus* spp. In the miRNA pathway (traced with dotted arrows), primary miRNA (pri-miRNA) is first transcribed from genomic DNA by RNA polymerase II. Drosha associates with DGCR8 (referred to as Pasha in flies and nematodes) and cleaves pri-miRNA to generate precursor miRNA (pre-miRNA) which is exported from the nucleus, primarily by Exportin (XPO)-5, into the cytoplasm where it is diced into smaller ∼22-nt miRNA duplexes by Dicer protein. The miRNA duplex is incorporated into the RNA-induced silencing complex (RISC) (now known as miRISC) where it is unwound into two separate strands by Argonaute 2 (AGO-2). The guide (antisense) strand, through AGO-2, locates and pairs insufficiently with a cognate cytosolic mRNA sequence, while AGO-2 mediates translational repression, and hence decay, of the cognate mRNA by recruiting TNRC6, a GW182 protein. The siRNA pathway (traced with bold arrows) is triggered by the introduction of exogenous long dsRNA into the cell. Long dsRNA substrate is diced into ∼22-nt siRNA duplexes. Short ∼22-nt siRNAs introduced directly into the cell does not undergo Dicer cleavage. The siRNA duplex is loaded into the RISC, now called siRISC, and follows the same unwinding and pairing process. The guide siRNA strand however forms a full complementary pairing with the cognate mRNA sequence, culminating in degradative cleavage of the mRNA sequence. P-bodies are cytoplasmic ribonucleoprotein granules that act as sites for mRNA translational repression or degradation. Negative RNAi regulators (RNAi inhibitors) may interfere with the overall process at specific stages. **C.** GO enrichment plot showing predicted biological process (BP), cellular components (CC), and molecular function (MF) terms associated with the identified *B. glabrata* proteins sequences. False discovery rate (FDR) values show significance level of enrichment. Signal is the weighted harmonic mean between the observed versus expected ratio and -log (FDR), which ensures more intuitive ordering of enriched terms. Gene count is the number of protein-coding genes in each enrichment term. For each GO category, the top 10 terms ranked by their signals were shown. List of genes under each GO term is available in the Supplementary Material 2.

### Expression levels of Ago2, Dcr1, and Eri1 genes were independent of snail size

Agarose gel analysis showed the expected amplicon size for each specifically amplified primer set (Figure 8B). Based on the snail size, the expression of *Dcr1*, *Ago2*, and *Eri1* suggested differences in the average relative mRNA expression levels, but these differences were not significant (*P* > 0.05) between groups (Figures 8 C-E).

**Figure 8.**
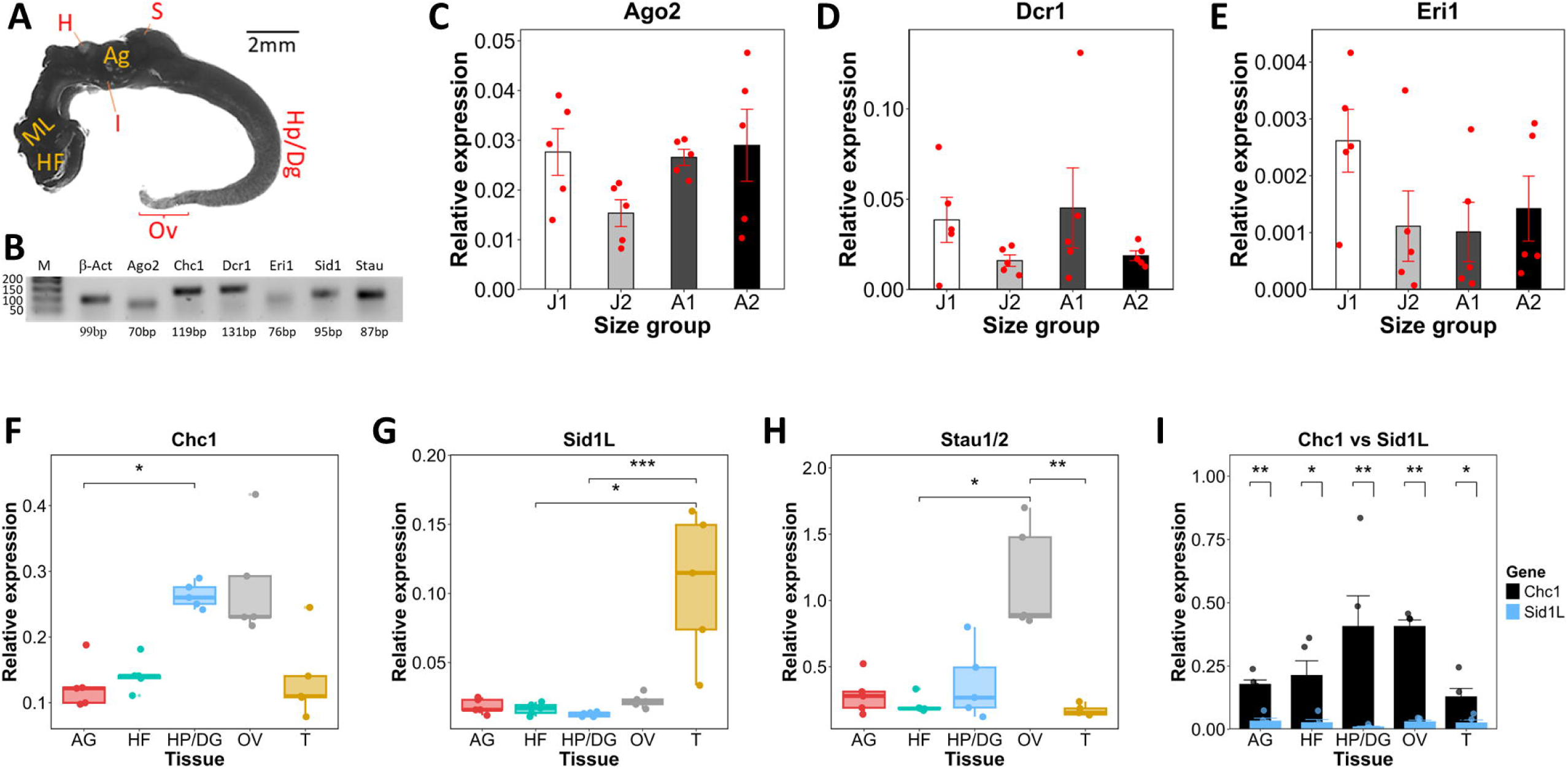
Gene expression analysis of selected putative RNAi genes in *Biomphalaria glabrata*. **A.** Anatomy of whole *Biomphalaria glabrata* soft body. HF, head-foot region; ML, mantle lobe; H, heart; AG, albumen gland; S, stomach; Hp/Dg, hepatopancreas/digestive gland; Ov, ovotestis; trunk (T, a multi-organ section containing the mantle lobe, lung, pseudobranch, intestine, stomach, heart, and kidney). **B.** Agarose gel image of real-time PCR products. **C.** Relative mRNA expression level of Argonaute 2, *Ago2* **D.** Dicer 1, *Dcr1* and **E.** enhancer of RNA interference, *Eri1*, in *B. glabrata* juvenile and adult snails. Comparisons of relative mRNA expression of **F.** clathrin heavy chain 1, *Chc1*, **G.** systemic RNA interference defective protein 1-like, *Sid1L*, **H.** staufen, *Stau1/2*, and **I.** Chc1 vs Sid1L, in *B. glabrata* HF, OV, AG, HP/DG and T tissue lysates. * *P* ≤ 0.05, ** *P* ≤ 0.01, and *** *P* ≤ 0.001.

### Enriched expression of Chc1, Sid1-like, and Stau1/2 genes were tissue-specific

Tissue-specific expressions of *Chc1*, *Sid1L*, and *Stau* were assessed to determine their localisation and extrapolate their roles in the snail RNAi. In all isolated tissues, expression of the endocytic pathway *Chc1* gene was significantly higher than *Sid1L* (*P* = 0.008 in AG, HP, and OV; *P* = 0.016 in HF and T) (Figure 8I), notably in the hepatopancreas and the ovotestis (Figure 8F). *B. glabrata Sid1L* was found more highly expressed in the trunk area than other tissues isolated, especially in comparison with the hepatopancreas (*P* = 0.0009) and the head-foot region (*P* = 0.0399) (Figure 8G). Out of the four *B. glabrata* STAU orthologues, one transcript (XM_056016125.1) examined for expression analysis showed higher expression in the ovotestis than other tissues, most significantly relative to the trunk region (*P* = 0.0036) and the head-foot tissue (*P* = 0.0399) (Figure 8H).

## Discussion

This study delineates the diversity of RNAi effectors in aquatic parasite-transmitting snails and provides insight into the scope and constraints of their functional roles. The reduced BUSCO completeness for the *L. staginalis* genome suggests that the genome does not contain as many pan-eukaryotic genes as other snail genomes examined in this study. In the original study where the genome assembly was assessed under a mollusc-specific lineage (mollusca_odb10), BUSCO completeness scored 91.8% (Koene et al., 2024). This indicates that the BUSCO outcome was strongly influenced by gene prediction settings, including the lineage choice. Although RNAi effectors are considered pan-eukaryotic, differences in the genome completeness of the snails did not impact the recovery of RNAi effector proteins. Interestingly, the number of RNAi effector proteins identified in *L. stagnalis* was comparable to but higher than in the *Biomphalaria* and *Bulinus* species examined. Two factors may have contributed to this pattern. First, searches for RNAi effector proteins were performed using the *L. staginalis* transcriptome, which may capture genes that are fragmented or ‘missing’ but not biologically absent in the genome assemblies. Second, phylogenetic divergence between lymnaeid (*Lymnaea*) and planorbid (*Biomphalaria* and *Bulinus*) snails might have caused family-specific loss or expansion of RNAi gene sets. Altogether, the results explain the highly conservative nature of the core RNAi effectors, even when genome completeness is less robust.

While we found that the snails lack classical orthologues of dsRNA-importing SID-1 protein, they may instead employ the clathrin-mediated endocytic (CME) pathway for dsRNA internalisation, as has been found in other organisms including insects (Xiao et al., 2015), mites (Zhou et al., 2023), and fungi (Wytinck et al., 2020). However, this alternative pathway may be slower and may require higher concentrations of dsRNA to initiate a systemic RNAi response (Feinberg and Hunter, 2003; Kobayashi et al., 2012). Importantly, CME is only one of the several core endocytic trafficking pathways in eukaryotes. It is highly selective in its endocytosed cargo, repelling many other substances that are imported via the non-CME pathways (Kumari et al., 2010).

Snail SID1-like orthologues showed closer evolutionary proximities to *Ce*CHUP-1 (also known as TAG-130) and *Hs*SIDT-1, both which are more exclusively involved in cholesterol uptake, rather than dsRNA uptake (Valdes et al., 2012; Mendez-Acevedo et al., 2017). In *C. elegans*, SID-1 is ubiquitously expressed in non-neuronal tissues, especially in the cell periphery of externally exposed tissues where it readily imports exogenous dsRNAs (Winston *et al*., 2002). Conversely, the paralogous *Ce*CHUP-1 is densely localised in the nematode’s intestine where most dietary cholesterol uptake occurs (Valdes et al., 2012). In this study, *Sid1L* gene also showed localised and high expression in the trunk section, which included the snail’s intestine and other organs listed previously. In *B. glabrata*, however, the digestive gland–gonad (DGG) complex, encompassing hepatopancreas and ovotestis, form the main site for cholesterol uptake (Shetty et al., 1990). In these organs, we observed no increased expression of *Sid1L* gene, suggesting that while the snail SID1L/CHUP1L orthologues are structurally competent of cholesterol uptake, their main activities are limited to certain organs and tissues outside the DGG complex. A major limitation here is that we were unable to achieve organ-specific resolution of the trunk area where *Sid1L* was significantly overexpressed. A similar study in *Litopenaeus vannamei* shrimp found high expression of *Sid1L* gene in the gills, while CME mainly facilitated uptake of ingested dsRNA into the hepatopancreatic cells (Maruekawong et al., 2022). This provides the most proximate insight as gills in shrimp are analogous to pseudobranch in *Biomphalaria*, which partly constitutes the snail trunk tissue. Moreover, we similarly found in the snail hepatopancreas (and ovotestis) a significant enrichment of *Chc1*, the main effector of the CME pathway. Previously, fluorescence microscopy showed hepatopancreas and ovotestis as the major sites for RNAi uptake in *B. glabrata* (Knight et al., 2011). Findings from the present study, reinforced by the observations from previous works, further support the clathrin-mediated cellular uptake of silencing RNAs in *B. glabrata*, mainly through the hepatopancreas and ovotestis.

Interestingly, more recent works showed that the cholesterol-trafficking *Hs*SIDT-1 protein facilitates dsRNA uptake via its phospholipase D (PLD) activity (Sun et al., 2024), at low pH (Zheng et al., 2024), although knockdown of clathrin in the same human embryonic kidney cell line (HEK293T) most markedly attenuated dsRNA uptake (Sun et al., 2024). Mammalian studies further revealed that CHUP1-like proteins may specialise in importing only cholesterol-conjugated dsRNAs (Brück et al., 2015; Mendez-Acevedo et al., 2017), a method that has also significantly enhanced RNAi delivery in *B. glabrata* (Portet et al., 2021). Together, this evidence suggests that intermediate snail hosts import external silencing RNAs potentially via both CME- and CHUP1L/SID1L-dependent pathways, as similarly noted in *Leptinotarsa decemlineata* beetle (Cappelle et al., 2016) and *L. vannamei* shrimp (Maruekawong et al., 2022), in a likely tissue-specific manner.

Dicer (DCR) as well as AGO proteins vary in their domain, microdomain, and amino acid structures. These architectural variations commonly underscore evolutionary specialisation of their isoforms toward distinct small RNA pathways (Nakanishi et al., 2012; Park et al., 2011). While the Platform–PAZ region of DCR enzymes consistently conserves its 3’-end binding pockets required for pre-siRNA processing, the 5’-end binding pockets that is critical for pre-miRNA processing is not conserved in siRNA-restricted DCRs (e.g. in *Dm*DCR-2) (Park et al., 2011; Lee et al., 2023). Results from this study however indicate that each snail species examined retained the structural requirements in their DCR to mediate both miRNA and siRNA biogenesis. This is potentially in an ATP-independent manner for *Biomphalaria* species, as DExH (HEL1) moiety of the helicase domain that facilitates ATP hydrolysis was not found. Such DCRs may be inefficient in cleaving blunt-ended siRNA precursors (dsRNAs without 2-nt 3’ terminal overhangs) (Donelick et al., 2020).

Prior to this study, phylogenetic analysis suggested an evolutionary loss of siRNA-processing AGO proteins in molluscs (Formaggioni et al., 2024). Our phylogenetic inference similarly suggested that snail AGOs evolved within the miRNA-specific AGO clade. Although phylogenetic analyses did not recover clear cleavage-competent siRNA AGOs in the snails, multiple sequence alignments revealing full conservation of the Piwi catalytic residues, coupled with many successful reports of dsRNA/siRNA-based silencing in snails (Xu et al., 2023; Portet et al., 2021) show otherwise. These findings suggest that AGO-2-mediated mRNA cleavage in the snails may be evolutionarily divergent, yet mechanistically canonical.

Across the groups of organisms examined, STAU protein exhibits functional plasticity in RNA metabolism. STAU-1 and STAU-2 are negative regulators of RNAi (Ren et al., 2016; Ehses et al., 2022), except coleoptera-specific StauC proteins that promote dsRNA-to-siRNA processing (Yoon et al., 2018). Unlike the putatively cytosolic snail STAU orthologues, N-terminal signal peptides found in StauC sequences indicate their typical localisation in the endoplasmic reticulum, which suggests an ability to process dsRNA prior to secretion for inter- or intra-cellular transport (Koo and Palli, 2024). Conversely, the fourth dsRNA-binding domain (RBD4) found only in non-coleopteran STAU sequences may be involved in the RNAi inhibition process. Many metazoans including *C. elegans* (LeGendre et al., 2013), *D. melanogaster* (Micklem et al., 2000), zebrafish (Ramasamy et al., 2006), frog (Allison et al., 2004; Bilogan and Horb, 2012), and mouse (Cao et al., 2016) exhibit high and stable expression of STAU-1 and STAU-2 in the oocytes, ovaries, and embryos. We similarly found the highest expression of *Stau* in the snail ovotestis. Considering both the function and the pattern of expression, it may be suggested that STAU-1/2 shields key developmental transcripts from the cell’s own RNAi machinery.

The adenosine deaminases acting on RNAs (ADARs), found in all snail species examined, are also RNAi inhibitors that may play contrasting roles in RNAi processing. These proteins, typically containing one or multiple dsRNA-binding domains, an A-deaminase domain, and one or more Z-binding domains, can edit dsRNAs by converting adenosine (A) residues into inosines (I). This A-to-I editing antagonises RNAi process by altering sequence regions that are required for small RNA precursor binding and/or cleavage by Drosha and Dicer proteins (Nishikura, 2010). While the RNA editor role of ADAR is facilitated by its ADAR/ADAR homodimerisation, ADAR/DCR heterodimerisation may rather define its role as an RNAi promoter (Ota et al., 2013).

The generation of secondary endogenous siRNAs (endo-siRNAs), following the introduction of initial RNAi triggers, drives a more robust RNAi response in some organisms. RNA-dependent RNA polymerases (RdRPs), majorly RRF-1 and EGO-1, and their upstream proteins RDE-10 and RDE-11, facilitate the production of endo-siRNAs (Zhang and Ruvkun, 2012), none of which were present in the snails. The phylogenetic distribution of RdRPs among metazoans is sporadic. They appear to have been lost in numerous lineages including gastropods (Pinzon et al., 2019). However, an RdRP-independent RNAi amplification mechanism mediated by endogenous reverse transcriptases has been suggested in *Drosophila* (Tassetto et al., 2017), and may represent an alternative mechanism for endo-siRNA production in many other organisms.

Core components responsible for transgenerational RNAi in *C. elegans* are the nuclear RNAi defective NRDE-1, NRDE-2, NRDE-3 and NRDE-4. The NRDE-3, a secondary AGO, binds and shuttles secondary siRNAs from the cytoplasm into the nucleus where it associates with other NRDE proteins (NRDE-2, -4 and -1, sequentially) to mediated transcriptional silencing of nascent mRNA through histone methylation. This way, the silencing effect is heritable by the progeny (Burton et al., 2011). Our in silico analysis recovered no orthologues of the core nuclear RNAi effectors in the snails examined, and no evidence for heritable RNAi has been published from the phylum Mollusca. Substantiating all findings from the in silico data, the significantly low *P*-value of the PPI network indicates that the putative protein orthologues found in the snails could interact to form functional RNAi units. The PPI networks were consistent with the common functional grouping of the RNAi effectors, as well as the general mechanistic RNAi pathways.

Argonaute-2 (*Ago-2*) and *Dcr-1* are two major dsRNA-processing genes, while *Eri1* is a major RNAi-inhibiting gene. Our qPCR analyses of these genes between snail developmental stages displayed stochastic variation among size-matched replicates. This indicated that RNAi efficiency is not influenced primarily by snail size. Shell size itself has imperfect correlation with snail’s age. Age-related differences, along with microenvironmental heterogeneity such as individual responses to stress, could contribute to temporal fluctuations in mRNA expression, even under uniform rearing conditions. Importantly, these observations remain consistent with existing experimental evidence. In *B. glabrata*, for example, RNAi-mediated knockdown has been successfully achieved across snails of different sizes, ranging from juveniles to adults (Portet et al., 2021; Xu et al., 2023).

Overall, our study has contributed new mechanistic insights into the RNAi biology of these snail species relevant to helminth disease transmission. We showed that RNAi effectors diversity between these snail vectors is consistent, suggesting that similar mechanisms and degree of susceptibility to exogenous gene silencing occur across the species. Despite lineage-specific absence of some effector components, our findings underscore the wide potential for RNAi use in trematode intermediate host snails as an experimental tool. These approaches can be used for functional genomics or as a future RNAi-based disease control strategy. Further work, including spatial and functional validation of these putative RNAi effectors, will strengthen the mechanistic framework established here. The development of RNAi approaches that can be manipulated more reliably and efficiently in parasite-vectoring snails, will substantially enhance our understanding of these important organisms.

## Data availability

All codes and data files used in data analysis are available on GitHub, at https://github.com/DamiF25/Snail_RNAi_effectors.

## Acknowledgements

We are grateful to Dr Gabriel Rinaldi for kindly providing the initial snail populations in the year 2020.

## Author’s contribution

PM, DOF, and CL conceived and designed the study. DOF, CL, and PM performed bioinformatic analyses. DOF performed gene expression analysis. DOF collated all data for analysis and visualisation. DOF wrote the initial draft. CL, DOF, DW, GNG and PM revised the initial draft and approved its final form.

## References

Abramson J, Adler J, Dunger J, Evans R, Green T, Pritzel A, Ronneberger O, Willmore L, Ballard AJ, Bambrick J, Bodenstein SW, Evans DA, Hung C, O’neill M, Reiman D, Tunyasuvunakool K, Wu Z, Žemgulytė A, Arvaniti E, Beattie C, Bertolli O, Bridgland A, Cherepanov A, Congreve M, Cowen-Rivers AI, Cowie A, Figurnov M, Fuchs FB, Gladman H, Jain R, Khan YA, Low CMR, Perlin K, Potapenko A, Savy P, Singh S, Stecula A, Thillaisundaram A, Tong C, Yakneen S, Zhong ED, Zielinski M, Žídek A, Bapst V, Kohli P, Jaderberg M, Hassabis D and Jumper JM (2024). Accurate structure prediction of biomolecular interactions with AlphaFold 3. Nature, 630, 493–500. doi: 10.1038/s41586-024-07487-w

Allan ERO, Tennessen JA, Bollmann SR, Hanington PC, Bayne CJ and Blouin MS (2017). Schistosome infectivity in the snail, Biomphalaria glabrata, is partially dependent on the expression of Grctm6, a Guadeloupe Resistance Complex protein. PLoS Negl Trop Dis, 11(2), e0005362. doi: 10.1371/journal.pntd.0005362

Allison R, Czaplinski K, Git A, Adegbenro E, Stennard F, Houliston E and Standart N (2004). Two distinct Staufen isoforms in *Xenopus* are vegetally localized during oogenesis. RNA, 10(11), 1751–63. doi: 10.1261/rna.7450204

Arraes FBM, Martins-De-Sa D, Noriega Vasquez DD, Melo BP, Faheem M, De Macedo LLP, Morgante CV, Barbosa J, Togawa RC, Moreira VJV, Danchin EGJ and Grossi-De-Sa MF (2021). Dissecting protein domain variability in the core RNA interference machinery of five insect orders. RNA Biol, 18(11), 1653–1681. doi: 10.1080/15476286.2020.1861816

Artimo P, Jonnalagedda M, Arnold K, Baratin D, Csardi G, De Castro E, Duvaud S, Flegel V, Fortier A, Gasteiger E, Grosdidier A, Hernandez C, Ioannidis V, Kuznetsov D, Liechti R, Moretti S, Mostaguir K, Redaschi N, Rossier G, Xenarios I and Stockinger H (2012). ExPASy: SIB bioinformatics resource portal. Nucleic Acids Research, 40(W1), W597–603. doi: 10.1093/nar/gks400

Baron OL, Van West P, Industri B, Ponchet M, Dubreuil G, Gourbal B, Reichhart JM and Coustau C (2013). Parental transfer of the antimicrobial protein LBP/BPI protects *Biomphalaria glabrata* eggs against oomycete infections. PLoS Pathog, 9(12), e1003792. doi: 10.1371/journal.ppat.1003792

Baum J, Papenfuss AT, Mair GR, Janse CJ, Vlachou D, Waters AP, Cowman AF, Crabb BS and De Koning-Ward TF (2009). Molecular genetics and comparative genomics reveal RNAi is not functional in malaria parasites. Nucleic Acids Res, 37(11), 3788–3798. doi: 10.1093/nar/gkp239

Benjamini Y and Hochberg Y (1995). Controlling the false discovery rate: A practical and powerful approach to multiple testing. J R Stat Soc Series B Stat Methodol, 57, 289–300. doi:

Bilogan CK and Horb ME (2012). *Xenopus* staufen2 is required for anterior endodermal organ formation. Genesis, 50(3), 251–9. doi: 10.1002/dvg.22000

Blum M, Andreeva A, Florentino LC, Chuguransky SR, Grego T, Hobbs E, Pinto BL, Orr A, Paysan-Lafosse T, Ponamareva I, Salazar GA, Bordin N, Bork P, Bridge A, Colwell L, Gough J, Haft DH, Letunic I, Llinares-López F, Marchler-Bauer A, Meng-Papaxanthos L, Mi H, Natale DA, Orengo CA, Pandurangan AP, Piovesan D, Rivoire C, Sigrist CJA, Thanki N, Thibaud-Nissen F, Thomas PD, Tosatto SCE, Wu CH and Bateman A (2025). InterPro: the protein sequence classification resource in 2025. Nucleic Acids Res, 53(D1), D444–D456. doi: 10.1093/nar/gkae1082

Brück J, Pascolo S, Fuchs K, Kellerer C, Glocova I, Geisel J, Dengler K, Yazdi AS, Röcken M and Ghoreschi K (2015). Cholesterol modification of p40-specific small interfering RNA enables therapeutic targeting of dendritic cells Journal of Immunology, 195(5), 2216–2223. doi: 10.4049/jimmunol.1402989

Burton NO, Burkhart KB and Kennedy S (2011). Nuclear RNAi maintains heritable gene silencing in *Caenorhabditis elegans*. Proc Natl Acad Sci U S A, 108(49), 19683–19688. doi: 10.1073/pnas.1113310108

Cao Y, Du J, Chen D, Wang Q, Zhang N, Liu X, Liu X, Weng J, Liang Y and Ma W (2016). RNA-binding protein Stau2 is important for spindle integrity and meiosis progression in mouse oocytes. Cell Cycle, 15(19), 2608–2618. doi: 10.1080/15384101.2016.1208869

Cappelle K, De Oliveira CF, Van Eynde B, Christiaens O and Smagghe G (2016). The involvement of clathrin-mediated endocytosis and two Sid-1-like transmembrane proteins in double-stranded RNA uptake in the Colorado potato beetle midgut. Insect Mol Biol, 25(3), 315–23. doi: 10.1111/imb.12222

Charoonnart P, Taunt HN, Yang L, Webb C, Robinson C, Saksmerprome V and Purton S (2023). Transgenic microalgae expressing double-stranded RNA as potential feed supplements for controlling white spot syndrome in shrimp aquaculture. Microorganisms, 11(8), 1893. doi: 10.3390/microorganisms11081893

Ciechanowska K, Pokornowska M and Kurzynska-Kokorniak A (2021). Genetic Insight into the Domain Structure and Functions of Dicer-Type Ribonucleases. Int J Mol Sci, 22(2). doi: 10.3390/ijms22020616

Cock PJA, Antao T, Chang JT, Chapman BA, Cox CJ, Dalke A, Friedberg I, Hamelryck T, Kauff F, Wilczynski B and De Hoon MJL (2009). Biopython: freely available Python tools for computational molecular biology and bioinformatics. Bioinformatics, 25(11), 1422–1423. doi: 10.1093/bioinformatics/btp163

Crooks GE, Hon G, Chandonia J and Brenner SE (2004). WebLogo: a sequence logo generator. Genome Res, 14, 1188–1190. doi: 10.1101/gr.849004

Donelick HM, Talide L, Bellet M, Aruscavage PJ, Lauret E, Aguiar E, Marques JT, Meignin C and Bass BL (2020). In vitro studies provide insight into effects of Dicer-2 helicase mutations in *Drosophila melanogaster*. RNA, 26(12), 1847–1861. doi: 10.1261/rna.077289.120

Ehses J, Schlegel M, Schroger L, Schieweck R, Derdak S, Bilban M, Bauer K, Harner M and Kiebler MA (2022). The dsRBP Staufen2 governs RNP assembly of neuronal Argonaute proteins. Nucleic Acids Res, 50(12), 7034–7047. doi: 10.1093/nar/gkac487

Fantini J, Di Scala C, Evans LS, Williamson PT and Barrantes FJ (2016). A mirror code for protein-cholesterol interactions in the two leaflets of biological membranes. Scientific Reports, 6, 21907. doi: 10.1038/srep21907

Feinberg EH and Hunter CP (2003). Transport of dsRNA into cells by the transmembrane protein SID-1. Science, 301(5639), 1545–1547. doi: 10.1126/science.1087117

Finn RD, Clements J, Arndt W, Miller BL, Wheeler TJ, Schreiber F, Bateman A and Eddy SR (2015). HMMER web server: 2015 update. Nucleic Acids Res, 43(W1), W30–8. doi: 10.1093/nar/gkv397

Fire A, Xu S, Montgomery MK, Kostas SA, Driver SE and Mello CC (1998). Potent and specific genetic interference by double-stranded RNA in *Caenorhabditis elegans*. Nature, 391, 806–811. doi: 10.1038/35888

Formaggioni A, Cavalli G, Hamada M, Sakamoto T, Plazzi F and Passamonti M (2024). The evolution and characterization of the RNA interference pathways in Lophotrochozoa. Genome Biol Evol, 16(5), evae098. doi: 10.1093/gbe/evae098

Ghosh SKB, Hunter WB, Park AL and Gundersen-Rindal DE (2018). Double-stranded RNA oral delivery methods to induce RNA interference in phloem and plant-sap-feeding hemipteran insects. Journal of Visualized Experiments, (135). doi: 10.3791/57390

Goldfarb T, Kodali VK, Pujar S, Brover V, Robbertse B, Farrell CM, Oh DH, Astashyn A, Ermolaeva O, Haddad D, Hlavina W, Hoffman J, Jackson JD, Joardar VS, Kristensen D, Masterson P, Mcgarvey KM, Mcveigh R, Mozes E, Murphy MR, Schafer SS, Souvorov A, Spurrier B, Strope PK, Sun H, Vatsan AR, Wallin C, Webb D, Brister JR, Hatcher E, Kimchi A, Klimke W, Marchler-Bauer A, Pruitt KD, Thibaud-Nissen F and Murphy TD (2025). NCBI RefSeq: reference sequence standards through 25 years of curation and annotation. Nucleic Acids Res, 53(D1), D243–D257. doi: 10.1093/nar/gkae1038

Hallgren J, Tsirigos KD, Pedersen MD, Almagro Armenteros JJ, Marcatili P, Nielsen H, Krogh A and Winther O (2022). DeepTMHMM predicts alpha and beta transmembrane proteins using deep neural networks. doi: 10.1101/2022.04.08.487609

Howe KL, Bolt BJ, Shafie M, Kersey P and Berriman M (2017). WormBase ParaSite - a comprehensive resource for helminth genomics. Mol Biochem Parasitol, 215, 2–10. doi: 10.1016/j.molbiopara.2016.11.005

Huang S, Yoshitake K, Asaduzzaman M, Kinoshita S, Watabe S and Asakawa S (2021). Discovery and functional understanding of miRNAs in molluscs: a genome-wide profiling approach. RNA Biol, 18(11), 1702–1715. doi: 10.1080/15476286.2020.1867798

Kall L, Krogh A and Sonnhammer EL (2007). Advantages of combined transmembrane topology and signal peptide prediction - the Phobius web server. Nucleic Acids Res, 35(Web Server issue), W429–32. doi: 10.1093/nar/gkm256

Kim D, Langmead B and Salzberg SL (2015). HISAT: a fast spliced aligner with low memory requirements. Nat Methods, 12(4), 357–60. doi: 10.1038/nmeth.3317

Knight M, Miller A, Liu Y, Scaria P, Woodle M and Ittiprasert W (2011). Polyethyleneimine (PEI) mediated siRNA gene silencing in the *Schistosoma mansoni* snail host, *Biomphalaria glabrata*. PLoS Negl Trop Dis, 5(7), e1212. doi: 10.1371/journal.pntd.0001212

Kobayashi I, Tsukioka H, Komoto N, Uchino K, Sezutsu H, Tamura T, Kusakabe T and Tomita S (2012). SID-1 protein of *Caenorhabditis elegans* mediates uptake of dsRNA into *Bombyx* cells. Insect Biochem Mol Biol, 42(2), 148–54. doi: 10.1016/j.ibmb.2011.11.007

Koene JM, Jackson DJ, Nakadera Y, Cerveau N, Madoui MA, Noel B, Jamilloux V, Poulain J, Labadie K, Da Silva C, Davison A, Feng ZP, Adema CM, Klopp C, Aury JM, Wincker P and Coutellec MA (2024). The genome of the simultaneously hermaphroditic snail Lymnaea stagnalis reveals an evolutionary expansion of FMRFamide-like receptors. Sci Rep, 14(1), 29213. doi: 10.1038/s41598-024-78520-1

Koo J and Palli SR (2024). StaufenC facilitates utilization of the ERAD pathway to transport dsRNA through the endoplasmic reticulum to the cytosol. Proc Natl Acad Sci U S A, 121(26), e2322927121. doi: 10.1073/pnas.2322927121

Kumari S, Mg S and Mayor S (2010). Endocytosis unplugged: multiple ways to enter the cell. Cell Res, 20(3), 256–75. doi: 10.1038/cr.2010.19

Lee YY, Lee H, Kim H, Kim VN and Roh SH (2023). Structure of the human DICER-pre-miRNA complex in a dicing state. Nature, 615(7951), 331–338. doi: 10.1038/s41586-023-05723-3

Legendre JB, Campbell ZT, Kroll-Conner P, Anderson P, Kimble J and Wickens M (2013). RNA targets and specificity of Staufen, a double-stranded RNA-binding protein in *Caenorhabditis elegans*. J Biol Chem, 288(4), 2532–45. doi: 10.1074/jbc.M112.397349

Li H, Handsaker B, Wysoker A, Fennell T, Ruan J, Homer N, Marth G, Abecasis G, Durbin R and Genome Project Data Processing S (2009). The sequence alignment/map format and SAMtools. Bioinformatics, 25(16), 2078–9. doi: 10.1093/bioinformatics/btp352

Lu XT, Gu QY, Limpanont Y, Song LG, Wu ZD, Okanurak K and Lv ZY (2018). Snail-borne parasitic diseases: an update on global epidemiological distribution, transmission interruption and control methods. Infect Dis Poverty, 7(1), 28. doi: 10.1186/s40249-018-0414-7

Maruekawong K, Namlamoon O and Attasart P (2022). Systemic gene silencing from oral uptake of dsRNA in *Litopenaeus vannamei* requires both clathrin-mediated endocytosis and LvSID-1. Aquaculture, 548. doi: 10.1016/j.aquaculture.2021.737557

Mendez-Acevedo KM, Valdes VJ, Asanov A and Vaca L (2017). A novel family of mammalian transmembrane proteins involved in cholesterol transport. Sci Rep, 7(1), 7450. doi: 10.1038/s41598-017-07077-z

Meng EC, Goddard TD, Pettersen EF, Couch GS, Pearson ZJ, Morris JH and Ferrin TE (2023). UCSF ChimeraX: tools for structure building and analysis. Protein Sci, 32(11), e4792. doi: 10.1002/pro.4792

Micklem DR, Adams J, Grunert S and St Johnston D (2000). Distinct roles of two conserved staufen domains in oskar mrna localization and translation. EMBO J, 19(6), 1366–1377. doi: 10.1093/emboj/19.6.1366

Nakanishi K, Weinberg DE, Bartel DP and Patel DJ (2012). Structure of yeast Argonaute with guide RNA. Nature, 486(7403), 368–74. doi: 10.1038/nature11211

Nganso BT, Sela N and Soroker V (2020). A genome-wide screening for RNAi pathway proteins in Acari. BMC Genomics, 21(1), 791. doi: 10.1186/s12864-020-07162-0

Nishikura K (2010). Functions and regulation of RNA editing by ADAR deaminases. Annu Rev Biochem, 79, 321–49. doi: 10.1146/annurev-biochem-060208-105251

Nuez I and Felix MA (2012). Evolution of susceptibility to ingested double-stranded RNAs in Caenorhabditis nematodes. PLoS One, 7(1), e29811. doi: 10.1371/journal.pone.0029811

Omasits U, Ahrens CH, Muller S and Wollscheid B (2014). Protter: interactive protein feature visualization and integration with experimental proteomic data. Bioinformatics, 30(6), 884–886. doi: 10.1093/bioinformatics/btt607

Ota H, Sakurai M, Gupta R, Valente L, Wulff BE, Ariyoshi K, Iizasa H, Davuluri RV and Nishikura K (2013). ADAR1 forms a complex with Dicer to promote microRNA processing and RNA-induced gene silencing. Cell, 153(3), 575–89. doi: 10.1016/j.cell.2013.03.024

Öztürk-Çolak A, Marygold SJ, Antonazzo G, Attrill H, Goutte-Gattat D, Jenkins VK, Matthews BB, Millburn G, Dos Santos G, Tabone CJ and The Flybase Consortium (2024). FlyBase: updates to the *Drosophila* genes and genomes database. Genetics, 227(1), iyad211. doi: 10.1093/genetics/iyad211

Park JE, Heo I, Tian Y, Simanshu DK, Chang H, Jee D, Patel DJ and Kim VN (2011). Dicer recognizes the 5’ end of RNA for efficient and accurate processing. Nature, 475(7355), 201–5. doi: 10.1038/nature10198

Pertea M, Pertea GM, Antonescu CM, Chang TC, Mendell JT and Salzberg SL (2015). StringTie enables improved reconstruction of a transcriptome from RNA-seq reads. Nat Biotechnol, 33(3), 290–5. doi: 10.1038/nbt.3122

Pfaffl MW (2001). A new mathematical model for relative quantification in real-time RT-PCR. Nucleic Acids Res, 29(9), e45. doi: 10.1093/nar/29.9.e45

Pinzon N, Bertrand S, Subirana L, Busseau I, Escriva H and Seitz H (2019). Functional lability of RNA-dependent RNA polymerases in animals. PLoS Genet, 15(2), e1007915. doi: 10.1371/journal.pgen.1007915

Portet A, Galinier R, Lassalle D, Faille A, Gourbal B and Duval D (2021). Hemocyte siRNA uptake is increased by 5’ cholesterol-TEG addition in *Biomphalaria glabrata*, snail vector of schistosome. PeerJ, 9, e10895. doi: 10.7717/peerj.10895

Procter JB, Carstairs GM, Soares B, Mourao K, Ofoegbu TC, Barton D, Lui L, Menard A, Sherstnev N, Roldan-Martinez D, Duce S, Martin DMA and Barton GJ (2021). Alignment of biological sequences with Jalview. Methods Mol Biol, 2231, 203–224. doi: 10.1007/978-1-0716-1036-7_13

Queiroz FR, Silva LM, Jeremias WJ, Baba EH, Caldeira RL, Coelho PMZ and Gomes MS (2017). Differential expression of small RNA pathway genes associated with the *Biomphalaria glabrata/Schistosoma mansoni* interaction. PLoS One, 12(7), e0181483. doi: 10.1371/journal.pone.0181483

R Core Team. (2025). The R Project for Statistical Computing [Online]. Veinna, Austria. Available: https://www.R-project.org/ [Accessed 03 January 2026].

Ramasamy S, Wang H, Quach HN and Sampath K (2006). Zebrafish Staufen1 and Staufen2 are required for the survival and migration of primordial germ cells. Dev Biol, 292(2), 393–406. doi: 10.1016/j.ydbio.2006.01.014

Ren Z, Veksler-Lublinsky I, Morrissey D and Ambros V (2016). Staufen negatively modulates microRNA activity in *Caenorhabditis elegans*. G3 (Bethesda), 6(5), 1227–37. doi: 10.1534/g3.116.027300

Sanz-Ros J, Mas-Bargues C, Romero-Garcia N, Huete-Acevedo J, Dromant M and Borras C (2023). MicroRNA biogenesis pathway alterations in aging. Extracell Vesicles Circ Nucl Acids, 4(3), 486–501. doi: 10.20517/evcna.2023.29

Shannon P, Markiel A, Ozier O, Baliga NS, Wang JT, Ramage D, Amin N, Schwikowski B and Ideker T (2003). Cytoscape: a software environment for integrated models of biomolecular interaction networks. Genome Res, 13(11), 2498–2504. doi: 10.1101/gr.1239303

Shetty PH, Fried B and Sherma J (1990). Sterols in the plasma and digestive gland-gonad complex of *Biomphalaria glabrata* snails, fed lettuce versus hen’s egg yolk, as determined by GLC. Comp Biochem Physiol B, 96(4), 791–4. doi: 10.1016/0305-0491(90)90233-J

Sievers F and Higgins DG (2018). Clustal Omega for making accurate alignments of many protein sequences. Protein Sci, 27(1), 135–145. doi: 10.1002/pro.3290

Simao FA, Waterhouse RM, Ioannidis P, Kriventseva EV and Zdobnov EM (2015). BUSCO: assessing genome assembly and annotation completeness with single-copy orthologs. Bioinformatics, 31(19), 3210–2. doi: 10.1093/bioinformatics/btv351

Skala V, Walker AJ and Horak P (2020). Snail defence responses to parasite infection: The *Lymnaea stagnalis-Trichobilharzia szidati* model. Dev Comp Immunol, 102, 103464. doi: 10.1016/j.dci.2019.103464

Smith M, Yadav S, Fagunloye OG, Pels NA, Horton DA, Alsultan N, Borns A, Cousin C, Dixon F, Mann VH, Lee C, Brindley PJ, El-Sayed NM, Bridger JM and Knight M (2021). PIWI silencing mechanism involving the retrotransposon nimbus orchestrates resistance to infection with Schistosoma mansoni in the snail vector, Biomphalaria glabrata. PLoS Negl Trop Dis, 15(9), e0009094. doi: 10.1371/journal.pntd.0009094

Sternberg PW, Auken KV, Wang Q, Wright A, Yook K, Zarowiecki M, Arnaboldi V, Becerra A, Brown S, Cain S, Chan J, Chen WJ, Cho J, Davis P, Diamantakis S, Dyer S, Grigoriadis D, Grove CA, Harris T, Howe K, Kishore R, Lee R, Longden I, Luypaert M, Müller H, Nuin P, Quinton-Tulloch M, Raciti D, Schedl T, Schindelman G and Stein L (2024). WormBase 2024: status and transitioning to Alliance infrastructure. Genetics, 227(1), iyae050. doi: 10.1093/genetics/iyae050

Sun CR, Xu D, Yang F, Hou Z, Luo Y, Zhang CY, Shan G, Huang G, Yao X, Chen Y, Li Q and Zhou CZ (2024). Human SIDT1 mediates dsRNA uptake via its phospholipase activity. Cell Res, 34(1), 84–87. doi: 10.1038/s41422-023-00889-x

Szklarczyk D, Nastou K, Koutrouli M, Kirsch R, Mehryary F, Hachilif R, Hu D, Peluso ME, Huang Q, Fang T, Doncheva NT, Pyysalo S, Bork P, Jensen LJ and Von Mering C (2025). The STRING database in 2025: protein networks with directionality of regulation. Nucleic Acids Res, 53(D1), D730–D737. doi: 10.1093/nar/gkae1113

Tamura K, Stecher G and Kumar S (2021). MEGA11: Molecular Evolutionary Genetics Analysis Version 11. Mol Biol Evol, 38(7), 3022–3027. doi: 10.1093/molbev/msab120

Tassetto M, Kunitomi M and Andino R (2017). Circulating immune cells mediate a systemic RNAi-based adaptive antiviral response in *Drosophila*. Cell, 169, 314–325. doi: 10.1016/j.cell.2017.03.033

Teufel F, Almagro Armenteros JJ, Johansen AR, Gislason MH, Pihl SI, Tsirigos KD, Winther O, Brunak S, Von Heijne G and Nielsen H (2022). SignalP 6.0 predicts all five types of signal peptides using protein language models. Nat Biotechnol, 40(7), 1023–1025. doi: 10.1038/s41587-021-01156-3

The Uniprot Consortium (2024). UniProt: the Universal Protein Knowledgebase in 2025. Nucleic Acids Res, gkae1010, 1–9. doi: 10.1093/nar/gkae1010

Valdes VJ, Athie A, Salinas LS, Navarro RE and Vaca L (2012). CUP-1 is a novel protein involved in dietary cholesterol uptake in *Caenorhabditis elegans*. PLoS One, 7(3), e33962. doi: 10.1371/journal.pone.0033962

Whangbo JS, Weisman AS, Chae J and Hunter CP (2017). SID-1 domains important for dsRNA import in *Caenorhabditis elegans*. G3 (Bethesda), 7(12), 3887–3899. doi: 10.1534/g3.117.300308

Wilson RC and Doudna JA (2013). Molecular mechanisms of RNA interference. Annual Review of Biophysics, 42, 217–239. doi: 10.1146/annurev-biophys-083012-130404

Wytinck N, Sullivan DS, Biggar KT, Crisostomo L, Pelka P, Belmonte MF and Whyard S (2020). Clathrin mediated endocytosis is involved in the uptake of exogenous double-stranded RNA in the white mold phytopathogen *Sclerotinia sclerotiorum*. Sci Rep, 10(1), 12773. doi: 10.1038/s41598-020-69771-9

Xiao D, Gao X, Xu J, Liang X, Li Q, Yao J and Zhu KY (2015). Clathrin-dependent endocytosis plays a predominant role in cellular uptake of double-stranded RNA in the red flour beetle. Insect Biochem Mol Biol, 60, 68–77. doi: 10.1016/j.ibmb.2015.03.009

Xu S, Zhang Y-W-Q, Habib MR, Li S-Z, Yuan Y, Ke WH, Jiang N, Dong H and Zhao Q-P (2023). Inhibition of alternative oxidase disrupts the development and oviposition of *Biomphalaria glabrata* snails. Parasites Vectors, 16, 73. doi: 10.1186/s13071-022-05642-8

Yoon JS, Mogilicherla K, Gurusamy D, Chen X, Chereddy S and Palli SR (2018). Double-stranded RNA binding protein, Staufen, is required for the initiation of RNAi in coleopteran insects. Proc Natl Acad Sci U S A, 115(33), 8334–8339. doi: 10.1073/pnas.1809381115

Zapletal D, Kubicek K, Svoboda P and Stefl R (2023). Dicer structure and function: conserved and evolving features. EMBO Rep, 24(7), e57215. doi: 10.15252/embr.202357215

Zhang C and Ruvkun G (2012). New insights into siRNA amplification and RNAi. RNA Biol, 9(8), 1045–9. doi: 10.4161/rna.21246

Zheng L, Yang T, Guo H, Qi C, Lu Y, Xiao H, Gao Y, Liu Y, Yang Y, Zhou M, Nguyen HC, Zhu Y, Sun F, Zhang C-Y and Ji X (2024). Cryo-EM structures of human SID-1 transmembrane family proteins and implications for their low-pH-dependent RNA transport activity. Cell Res, 34, 80–83 doi: 10.1038/s41422-023-00893-1

Zhou H, Wan F, Jian Y, Guo F, Zhang M, Shi S, Yang L, Li S, Liu Y and Ding W (2023). Chitosan/dsRNA polyplex nanoparticles advance environmental RNA interference efficiency through activating clathrin-dependent endocytosis. Int J Biol Macromol, 253(Pt 4), 127021. doi: 10.1016/j.ijbiomac.2023.127021

