## Supplementary material 1 for "Conserved core RNAi machinery in trematode-vectoring snails indicates gene silencing potential in the absence of classical systemic and amplification effectors"

**Table 1.** Accessions to human, *Drosophila*, *Caenorhabditis* RNAi protein query sequences used

| Protein | *C. elegans* (WormBase) | *D. melanogaster* (FlyBase) | *H. sapiens* (UniProt) |
|---|---|---|---|
| SID-1 | CE30331 | | Q9NXL6-1 |
| SID-2 | CE33908 | | |
| SID-5 | CE02645 | | |
| CLINT-1 | CE03062 | | Q14677-1 |
| CHUP-1 | CE26397* | | |
| SID-3/ACK | CE33506 | | Q07912-1 |
| SMG-2 | CE28367 | | |
| SMG-5 | CE25134 | CG8954-PA* | |
| SMG-6 | CE32984 | CG6369* | |
| DYN-1 | | CG18102-PB | |
| CHC-1 | | CG9012-PA | |
| Light | | CG18028-PC | |
| ARF72A | | CG6025-PA | |
| Eater | | CG6124-PC | |
| SR-CI | | CG4099-PA | |
| RRF-3 | CE45624 | | |
| RRF-1 | CE27141 | | |
| RSD-2 | CE40653 | | |
| RSD-6 | CE32384 | | |
| EGO-1 | CE27140 | | |
| RDE-1 | CE28243 | | |
| RDE-10 | CE24371 | | |
| RDE-11 | CE36970 | | |
| ABT-1 | CE48446 | | |
| PGP-4 | CE44238 | | |
| PGP-11 | CE53819 | | |
| HAF-2 | CE51253 | | |
| HAF-6 | CE39850 | | |
| WHT-1 | CE52537 | | |
| WHT-3 | CE51368 | | |
| MRP-1 | CE29412 | | P33527-1* |
| PMP-1 | CE02546 | | |
| DCR-1 | CE47418 | CG4792-PA | Q9UPY3-1 |
| DCR-2 | | CG6493-PA | |
| DRH-1 | CE16988 | | |
| DRH-3 | CE36120 | | |
| DRSH-1 | CE29039 | | |
| DGCR8 | CE48538 | | Q8WYQ5-1 |
| PIR-1 | CE48135 | | |
| MES-2 | CE28067 | | |
| MES-3 | CE11046 | | |
| MES-6 | CE24796 | | |
| NRDE-1 | CE31415 | | |
| NRDE-2 | CE18165 | | Q9H7Z3* |
| NRDE-3 | CE04789 | | |
| NRDE-4 | CE52211 | | |
| ADAR-1 | CE32459 | | P55265-1 |
| ADAR-2 | CE25120 | CG12598-PA | P78563-1 |
| ADAR-3 | | | Q9NS39-1 |
| ALG-1 | CE31525 | | |
| PACT | | | O75569-1 |
| TNRC6A | | CG31992-PB | Q8NDV7-1 |
| PAN-2 | | | Q504Q3-1 |
| PAN-3 | | | Q58A45-1 |
| DDX-6 | | | P26196 |
| CCR4-NOT | | | A5YKK6-1 |
| DCP1 | | | Q9NPI6-1 |
| DCP2 | | | Q8IU60-1 |
| RAB-10 | CE14114 | | |
| EHBP-1 | CE05711 | | |
| STAU-1 | CE11107 | CG5753-PA | O95793-1 |
| STAU-2 | | | Q9NUL3-1 |
| EGH | | CG9659-PC | |
| VHA-16 | | CG32090-PA | |
| VHA-SFD | | CG17332-PA | |
| COG-2 | | CG6177-PA | |
| COG-3 | | CG3248-PA | |
| GMER | | CG3495-PA | |
| RAN | | | P62826 |
| TRS-20 | | CG5161-PA | |
| Pi3K59F | | CG5373-PA | |
| RAB-7 | | CG5915-PA | |
| AP-50 | | CG7057-PA | |
| NinaC | | CG5125-PB | |
| CG5382 | | CG5382 | |
| CG5434 | | CG5434 | |
| CG4572 | | CG4572 | |
| Hsc70-3 | | CG4147-PB | |
| Hsc70-4 | | CG4264-PE | |
| XPO-1 | CE23467 | CG13387-PA | |
| XPO-2 | CE27001 | | |
| XPO-3 | CE29726 | | |
| XPO-5 | | | Q9HAV4 |
| AIN-1 | CE03967 | | |
| AIN-2 | CE27563 | | |
| TSN-1 | CE02626 | | |
| MTR-4 | CE06562 | | |
| VIG-1 | CE11254 | | |
| R2D2 | CE00630 | CG7138-PA | |
| TRBP-2 | | | Q15633-1 |
| MUT-2 | CE11740 | | |
| MUT-7 | CE00370 | | |
| MUT-16 | CE40346 | | |
| RDE-2 | CE37005 | | |
| CID-1 | CE02015 | | |
| EKL-1 | CE05687 | | |
| EKL-4 | CE29836 | | |
| EKL-5 | CE44333 | | |
| EKL-6 | CE51593 | | |
| RHA-1 | CE39177 | | |
| GFL-1 | CE12388 | | |
| ZFP-1 | CE25003 | | |
| LIN-15B | CE15381 | | |
| SOMI-1 | CE36495 | | |
| TAF-11 | | CG4079-PA | |
| AGO-1 | | CG6671-PC | Q9UL18 |
| AGO-2 | CE28243 | CG7439-PB | Q9UKV8-1 |
| AGO-3 | | | Q9H9G7-1 |
| AGO-4 | | | Q9HCK5 |
| ALG-2 | CE32063 | | |
| XRN-1 | CE42924 | | |
| XRN-2 | CE42702 | | |
| REXD-1 | CE42034 | | |
| TBC-3 | CE35519 | | |
| ZFP-2 | CE00981 | | |
| ERI-3 | CE32072 | | |
| ERI-9 | CE01165 | | |
| ERI-1 | CE17214 | | |
| ERI-5 | CE39434 | | |
| ERI-6/7 | ACH90516.1 (NCBI) | | |

*Potentially non-RNAi protein but used for further analysis

**Table 2**. *Biomphalaria glabrata* primer sequences used for qPCR assays

| Gene | Transcript accession no. | Primer sequences (5'→3') | Amplicon size (bp) | Design tool |
|---|---|---|---|---|
| Ago2 | XM_013213009.2 | F: ATGATGGATGTGACGGGCAG<br>R: CTTGGGCTTTGGTCCGTCC | 70 | Primer-BLAST |
| Dcr1 | XM_056023171.1 | F: ACCCCTCTGATCCGTCATCA<br>R: GTCAGCAGCATCAGGACCTT | 131 | Primer3Plus |
| Sid1L | XM_056010244.1 | F: TGTTTTGGCTACTCCTGGTAATGA<br>R: CTCCACTGGACTCTGTAGATGC | 95 | Primer-BLAST |
| Chc1 | XM_013224298.2 | F: CACTGGGCAATGGAAGGAGA<br>R: ACTAGCAAGAGCCACTGCTG | 119 | Primer-BLAST |
| Eri1 | XM_056010161.1 | F: CACAGTGTGTTGAGGTGCGG<br>R: AGTTTTCGAGCCCATCACCC | 76 | Primer-BLAST |
| Stau | XM_056016125.1 | F: AGTGCCTGGACTGCTTCATC<br>R: GACTGGGAGCTTGCAGATGG | 87 | Primer-BLAST |
| β-Act | XM_013228447.1 | F: GCTTCCACCTCTTCATCTCTTG<br>R: GAACGTAGCTTCTGGACATCTG | 99 | Primer3Plus |

F, forward sequence; R, reverse sequence.

**Table 3.** Accessions to putative RNAi effector protein sequences in *Biomphlalaria glabrata*, *Biomphalaria pfeifferi*, and *Bulinus truncatus* snails

| **RNAi effector protein** | **NCBI accession no.** | | |
|---|---|---|---|
| | ***B. glabrata*** | ***B. pfeifferi*** | ***Bu. truncatus*** |
| Sid-1L/Chup-1L | XP_055866219.1 | KAK0069211.1 | KAH9507595.1 |
| Clint-1 | XP_013083372.2 | KAK0058365.1 | KAH9491911.1 |
| Ack | XP_013070020.2 | KAK0063636.1 | KAH9513559.1 |
| Chc-1 | XP_013079752.1 | KAK0047708.1 | KAH9502444.1 |
| Xpo-1 | XP_013069551.1 | KAK0059781.1 | KAH9496126.1 |
| Xpo-2 | XP_055884665.1 | KAK0052093.1 | KAH9496808.1 |
| Xpo-3 | KAI8781818.1 | KAK0064135.1 | KAH9494515.1 |
| Xpo-5 | XP_055878613.1 | KAK0042291.1 | KAH9505187.1 |
| Light/VPS41 | XP_013095699.1 | KAK0051675.1 | KAH9500097.1 |
| Vha-16 | XP_055891493.1 | KAK0041102.1 | KAH9509959.1 |
| VhaSFD | XP_055871714.1 | KAK0066067.1 | KAH9513776.1 |
| Trs-20 | XP_013066166.1 | KAK0050522.1 | KAH9492369.1 |
| Cog-2 | XP_055879772.1 | KAK0059790.1 | KAH9496136.1 |
| Cog-3 | XP_013085383.2 | KAK0059752.1 | KAH9512324.1 |
| Arf72A/Arl-1 | XP_055900743.1 | KAK0055341.1 | KAH9510156.1 |
| Haf-2 | XP_055878829.1 | KAK0043502.1 | KAH9507216.1 |
| Haf-6 | KAI8760177.1 | KAK0043972.1 | KAH9523570.1 |
| Pi3K59F | KAI8788980.1 | KAK0054667.1 | KAH9524584.1 |
| Gmer | XP_013096171.2 | KAK0052592.1 | KAH9503670.1 |
| Rab-7 | XP_013063134.1 | KAK0062410.1 | KAH9500806.1 |
| Rab-10 | XP_055884483.1 | KAK0062868.1 | KAH9504142.1 |
| Dyn | XP_055898036.1 | KAK0062499.1 | KAH9498991.1 |
| Tbc-3 | XP_055882966.1 | KAK0056700.1 | KAH9505195.1 |
| Pgp-4 | XP_055864038.1 | KAK0066802.1 | KAH9508579.1 |
| Pgp-11 | XP_055864045.1 | KAK0058015.1 | KAH9520292.1 |
| Wht-1 | XP_055873622.1 | KAK0042817.1 | KAH9524963.1 |
| Wht-3 | KAI8768363.1 | KAK0042817.1 | KAH9524970.1 |
| Pmp-1 | XP_055886984.1 | KAK0069404.1 | KAH9504675.1 |
| Mrp-1 | KAI8797123.1 | KAK0051767.1 | KAH9514937.1 |
| CG5382 | XP_013082575.2 | KAK0043496.1 | KAH9507220.1 |
| CG5434 | KAI8732962.1 | KAK0062380.1 | KAH9518917.1 |
| CG4572 | KAI8744072.1 | KAK0053040.1 | KAH9507790.1 |
| Ap-50/AP-2 | XP_055864660.1 | KAK0069834.1 | KAH9513477.1 |
| NinaC | XP_055869732.1 | KAK0049982.1 | KAH9524024.1 |
| Hsc70-3 | KAI8778920.1 | KAK0063581.1 | KAH9500761.1 |
| Hsc70-4 | XP_013082114.1 | KAK0067577.1 | KAH9498720.1 |
| Ehbp-1 | KAI8747180.1 | KAK0057021.1 | KAH9499140.1 |
| Ran | KAI8749373.1 | KAK0041501.1 | KAH9498686.1 |
| Abt-1 | KAI8796754.1 | KAK0056586.1 | KAH9488682.1 |

| **RNAi effector protein** | | **NCBI accession no.** | | |
|---|---|---|---|---|
| | | ***B. glabrata*** | ***B. pfeifferi*** | ***Bu. truncatus*** |
| Dcr-1 | a | XP_055879146.1 | KAK0058462.1 | KAH9519444.1 |
| | b | | | KAH9519447.1 |
| Drh-1/3 | | KAI8768888.1 | KAK0060216.1 | KAH9492753.1 |
| Drsh-1 | | KAI8778695.1 | KAK0063359.1 | KAH9500714.1 |
| Dgcr8 | | XP_055861605.1 | KAK0054976.1 | KAH9524193.1 |
| Taff-11 | | KAI8739574.1 | KAK0040952.1 | KAH9504782.1 |
| Pir-1 | | XP_013096378.2 | KAK0046578.1 | KAH9514491.1 |
| Trbp2 | | XP_013088062.1 | KAK0047136.1 | KAH9524692.1 |
| Smg-2 | | XP_013074693.1 | KAK0057905.1 | KAH9520158.1 |
| Ago-2 | a | XP_013068463.2 | KAK0055590.1 | KAH9494476.1 |
| | b | KAI8776705.1 | | |
| Tsn-1 | | KAI8757997.1 | KAK0053358.1 | KAH9499055.1 |
| Vig-1 | | XP_013081752.2 | KAK0069929.1 | KAH9523375.1 |
| Tnrc6 | | XP_013075548.1 | KAK0043145.1 | KAH9519740.1 |
| Ccr4-Not | | XP_055890769.1 | KAK0046373.1 | KAH9515531.1 |
| Pan-2 | | KAI8770369.1 | KAK0057343.1 | KAH9513218.1 |
| Pan-3 | | XP_055893130.1 | KAK0059555.1 | KAH9513810.1 |
| Ddx6 | | XP_013066850.1 | KAK0042408.1 | KAH9504297.1 |
| Dcp1 | | XP_055884034.1 | KAK0051889.1 | KAH9512487.1 |
| Dcp2 | | XP_055864573.1 | KAK0060280.1 | KAH9524493.1 |
| Mut-2 | | XP_013063605.2 | KAK0057656.1 | KAH9503294.1 |
| Mut-7 | | XP_013073446.2 | KAK0050820.1 | KAH9498986.1 |
| Cid-1 | | KAI8728918.1 | KAK0068679.1 | KAH9515295.1 |
| Elk-4 | | KAI8790464.1 | KAK0043116.1 | KAH9499113.1 |
| Elk-6 | | XP_055892149.1 | KAK0062717.1 | |
| Mes-2 | | KAI8761384.1 | KAK0044138.1 | KAH9498123.1 |
| Mes-6 | | KAI8791814.1 | KAK0053562.1 | KAH9525029.1 |
| Rha-1 | | KAI8771043.1 | KAK0041774.1 | KAH9505328.1 |
| Gfl-1 | | KAI8737473.1 | KAK0061368.1 | KAH9499362.1 |
| Zfp-1 | | KAI8756278.1 | KAK0049027.1 | KAH9500895.1 |
| Mtr-4 | | XP_013074790.2 | KAK0052105.1 | KAH9489916.1 |
| Eri-1 | | XP_055866136.1 | KAK0060180.1 | KAH9500731.1 |
| Adar-1 | a | XP_013065626.2 | KAK0049375.1 | KAH9494790.1 |
| | b | XP_055899947.1 | KAK0049597.1 | KAH9488909.1 |
| Adar-2-like | | XP_055860200.1 | KAK0049885.1 | KAH9505413.1 |
| Xrn-1 | | KAI8736246.1 | KAK0063715.1 | KAH9508699.1 |
| Xrn-2 | | KAI8740342.1 | KAK0050342.1 | KAH9502793.1 |
| Stau | a | XP_055872100.1 | KAK0066309.1 | KAH9490575.1 |
| | b | XP_055872101.1 | | |
| | c | KAK6960359.1 | | |
| | d | XP_055872099.1 | | |

### FASTA I

### Additional protein sequences used

#### Sid-1-like sequences

>*Acal*SID-1 (NCBI: XP_012944272.1)
MFNKAVVNHSGICTVALLLLFYELTAMVRCTEHLPQDSYDNSEIGNSLKKSHFRNAVVNGKNVIVESFKVSDDTHRSVGDVSSVDDRRSDEGFIAVSDQVASWSQ
PRFPVETVIVDAEFDQNYGNWTVNRTSQILFVYNYTETVNKTTAVRIRVTCSNATENYPVMFVARQQEGILSWRVPLEIGKGYSFWSVSRTLCPLDREHRQIIVKE
QLIYLTVSTMAFNNRDFNLTAQLLQDFELEHARAKTFSVAASRPKYYMYTFPENVETVLLKVTSPTETCMTVSVQTVKCPVFDLDTNVEYEGKHQTMSSQAAMFL
EKSDYPDMNSFYVVFITKPNNEDCEGFVEFSPLLPPGHASEVVKTVTVEVKETISDSQYDKAVFATLAVFLCFYLIAVVIGCVFAGCGLRKDIGDLTEEEREAKQLLFS
NISPLLPEQNVNSDYIQALHRPLAGVLDSGTSEEQAAILAQVSESEEDNFLNRTDSPGGYGAIPTNRGASKELGPVHPRGAASNSDSSEAYVNENDIDFLTDADSEK
EVFRTKTALFVSDLARKKRSKLSKLYKLYHWNLGTIAIFYGLPVAQLVLTYQKILRQTGNQDLCYYNFACAHPLGSYISSFNNVFSNIGYVLLGILFIIIVYYRSWHNRRVI
EKYGDLERKYGIPQHFGLFYAMGLALVMEGIMSSCYHICPNYSNFQFDTSFMYVIACLCMLKIYQARHPDINAKAHVAYFVMAVIIFLAVVGVVYGNTGLWVVYA
VVHIVLTIFLTAQIYYMGRWKVDRYIFKRFFLVIVSDCRRCSRPVYPNRFVMLLVGNVVNWAFALYGAIERPSDFASYLLAVFIGNLLLYCVFYVIMKLLSKERLSWLVI
VVILTSMVTWGGSLYFFFQHLTSWQKTPAGSRAGNRECILLEFYDSHDVWHFLSAISLFFSFLILLLLDDDVSLKRRDQIPVF

>*Lgig*SID1 (NCBI: XP_009055461.1)
MLSDADEDKRIVRTKTALFVSDLARKNHKKVSKNYKIYHWNLLTIGIFYGLPVIQLVFNYQKVLHQTGDEDICYYNFYCAHPLGVLSSFNNVFSNIGYVLLGILFIIIARR
RSFVRNNKEVAKLYGIPQHFGLYYAMGLSLILEGVMSACYHVCPNNSNFQFDTSFMYIIACLNMLKIYQTRHPDINAKAHTTYLSMAFIIFIAVIGVYYGTNIFWILYG
LVHMLVSLIISAQIYFMGMWKMDCGIFKRLFWLFRNDCLRCAKPIYPNRFVLLLIGNVINWSFAIYGVLNTPIDFATYLLAIFIGNLMLYVLFYIIMKLFSGEKIHILAIS
CIVITMITWAAALYFFFAHLTSWSVTPAKSREGNRPCILLEFYDAHDIWHFLSAIGMFFGFLIILTLDDDLVLKRRDQIPVF

>*Mgal*SID-1 (NCBI: VDI62648.1)
MEIRFQTSSASILSRLYSQKLSVIDGHFGNSYTNQTNENSTIIYKFEYKEIENKTTAVRISIKLLNTADEDFPLLFVARQQRGVISWQLPLPLENIYEYEYASRTLCPLDQQ
HRKADMKQVMYAEISSMSVKNLNFSLKASKISDFELKPGEVKTLSLTPSEPQYYMYTFQEYRFCYFKVESETGNCPVFDLDKDVEFSGTYQTVTKQGAITVEKKKY
MEESFYIVLVLKPKDYECNGIDEIQLPGEERTKHLTLRVYGTISSSKYYIGIVGALAIFGGFYLIAFMIGCCYHGCQKNRGVWIIEEEKSEGEGETDDLLNRSNTLPGSY
GATTGNESDVMDRSLHEIQTPSNTTVDIDSVDSSIDFLPDADKEKDVFRTKVRTPSNTTVDTDSVDSSIDFLPDADKEKDVFRTKVRTPSNTTVDTDSVDSSIDFLP
DADKEKDVFRTKTILFVVDLARKSRKKVAKSYSLFCRNLAAISIFYGLAVLQLVLTYQKVLNVTGNQDICYYNFDCAHPLGSLTSFNNVFSNVGYILLGILFLILVARRDVI
YKKAVQNDRKIGKELGIPQHFGLFYAMGLALVMEGLMSACYHVCPNYSNFQFDTSFMYIIACLCMLKIYQSRHPDISAKAHTSYMFMACVIFIAVIGV

>*Cvir*SID-1 (NCBI: XP_022322232.1)
MLHFSLVRFSIILVVYGLLQGSYATDANFSQTYSDIINTTVEAIYKFTYTEMRNRTTAVRITTALQSDDEKATMKHPVIVVVRQQQSTLSWTMPLYSKNSDGAFDSDT
VSRTLCPLDASHRPEKPLEQSFFVDVSTFSKQPLNFTLRAIKMESFEISAGVAKEIYFTPSEPQFYMYTFPEDVDSVLVHATSKDEKCAVMSIQTIECPVYDLNVNVEF
EGKYQTMTKQAAIQIQRTDYPEGAFYVVFVLKPLDSACSSGLETLSRVKKTISSDKYYWGMVFAVAVFGAFYLIAFLLGVIYHGCSKHRGIWMIPEYQDIKKDQDSG
HDGGDDLLSTVIRDNTSSQATNTSNSMDNHSVSSLDSEDVDFLKDADEEKDVFRTKTELFVYDLSRKKYKKQARNYAIYWRNLLTIAIFYGLPVLQLVFTYQKVLNV
TGNLDICYYNFNCAHPWGLVSSFNNIFSNIGYVSLGFLFLFLVYRRKLIYNMAVHREKMMKKEVGIPQHFGLFFGMGLALIMEGVMSACYHVCPNYSNFQFDTSF
MYIIACLCMLKIYQTRHPDISAKAHTSYLLMACVIFIAVVGVIYGDNIFWILFACVYMLFYLILSVHIYFMGRWSIDRGICRRICVAVRYDMCRCRRPMYPDRMLLLVI
GNLINWGIAVYGALKHPSDFATFLLAIFIGNLMIYCSFYIFMKLRYKEKINKLPLVIIIMTVLVWAVALYFFFFHLTTWQLTPSRSREGNRDCLLLEFYDAHDIWHFLSSI
ALFFSFLVLLTLDDDRFSTRRDQIHVIVNIDPMLKSIGKLFGRKRSSKKGEKGTVESAENDDSHEPENIAKHGPGARECRIAKTPVHEDAVVHLSAVEPGLCLSCSKDK
SAALYDYENHKLVDRWTGHERELTKILNGRTSNSIFSASRDKSVKMWRSGNPNFLQEFNDHDLVVTAIDLNHDDSWLCSGSRDNHVKLWDVNTAQCFLESSIHR
NLVTDVKFVPHSHLLVQTGEDKEVRLFDTRTMQVVHSFPRKHYIQMACDVSADGLYCVTCSNGFSGKGCEATLWDIRGRKIVHEYREHGEAVESCIFLPSINERQII
ATTSRDTTVRLWDMNTRECLSCFPIAGGGPITSIICYDDLSLCVSTFNQGIISLQLSPLDYSLRQTGQF

>*Ovul*SID-1 (NCBI: CAI9737789.1)
MYNRIRLIFSSQYQIIFLCILMNARQIYCNDYQANEFVSRTDTKEEQHNSTPSKQLSLTKHIKYDVSQTRDPYDYEIIKKPGEIPTTYVHNNATFGKEYNETVNQNKQL
EFIYTYEESENETKALRVSTSSSSAEEKYPIIFVVRQQEGVISWTVPFFLDFKYEFFHVSRTLCPLDQEHRANISKHQKIIIEVSSQVKESSDFLLQTSIVKNFKLKLDVEK
QVTVTPAEPLYYMFTFPDNVESVVVTGQSNDTSCMTISIQDIKCPIYDLDRNIRFHGKFQTMSKNSAIVVQKNLYSQNAFYVVLVLMLNNLKCSASDSEKVFPMGS
VRTKDVTILVEETISVAHYSTAIIASVGLFLSFYIVAGLALLIYNKCDKKPEFPEISELERERFLWGAAGLIPSDSDSDEKSALPQGYQTQRYGSINERRNDNVSAPLLNP
MSPSATTEDTSSVNSSLDTEDIDFLNDAEEEKDVFRTKTALFVSDLSRKGRKKLEKYGKVYNWNLWTIVIFYGLPVIQLVITYQKVLNVTGNEDTCYYNFKCAMPYKV
LSSFNNVFSNIGYILLGMLFLLLVWRRDLIHREIVKKYGKAEELYGIPQPYGLFYAMGLGLCMEGVMSACYHVCPTYSNFQFDTSFMYIIACLCMLRLYQTRHPDINA
KAHTAYLYMAFIIFIAVIGVVYQTPVFWIVFAAIHFVTTLVLSVQIYYMGRWKIDLGIFNRMRLMLCSDFIQCAKPMYKDRLILLIISNIMNWTFAVYGAIKKPSDFASY
LLAIFIGNLLFYLFFYLIMKLRSKETIKVWTRFYILMTGILWGFALYFFLKHLTSWQLSPAMSRDKNRECILLGFFDDHDIWHFLSASSMFFSFMILLTLDDDLQLIRRDR
IPVF

#### Coleoptera-specific Staufen (StauC) protein sequences

>*Ld*StauC (Kim et al 2021)
MLELTIHFNVLLVFLVAFCGGYSEEGIRQAFGAKSCIQASLSSLDLHSFGVKESQNSIIGNIKSARGAENTDKSSLSRLNELVQFNEIDYYFKLEKEEGLPHDKVFTVSL
TLGTETYIGEGKSLKKAKQNVAAIALHETRYETPPVKEPSENEESLTPTVMLNNLGAKLGIGITYYLIDKEKQHILNSNLVVSENSKKSYVQKLNDSIYSNKNLRMKK
DIENTKGPFKIKVQVGDQVFSGYAHSIQAARHQAASNALDFLIKNKDSLDLDCLKEGSEEQCKKAKQNLKSPVSLVYESAQMRKLDVEFEIIKEFGPPHKKTFVTEC
RVGHLTTTGEGRSKKASKKAAAEDMLEKMSELEPIPQEVQVKSMLKDKKKKNKKKKIIKNKLDEISMTVGNVIDSVVGFGKDILADKKGDDKTDKNSSDGPKSKK
SKKSENLKQTYQDQLLEMSNALNFEISYADFEEGSKHFSLLSLHINPEYLCFGEGSNKMQSRNKAADKGLDLLGKMGLFDILNDQKTVPLERDTKEAVHHVLEHHI
SQDKDEL

>*Tc*StauC (NCBI: XP_975506.3)
MFRFLVCAVILKWSFGSEFVNDLQSVFGDKHCVDASLISIPSNIDTYVMFSSESQNNVTKTPEKNIVATIYELAQFNEIECSYTLISEEGPPHAKIFKVSLHLGDELYE
GSGSSLKKAQQAAATLALQDTFYSHPPEKINTPQEKSLTPTVALNNVAAKLGIAVSYFLLNGKEEQTIDSYGAVDESYGKSKSSFPKWNATFDPLKVPLEDAQISGP

FAVRVKAGDSVFEGFGQTIQAARHDAASKALKDMEKRALLKDHVCSREDSVEDCKKLKDDLKSPITRVHEAAQLNNLALEFNVISESGKPHQRNFVTECKLGDYV
TEGEGRSKKESKRVAAEKMLQFLPQLHDTYNEKAIISGLQNKQKKKKNKKNKVKRENVDTMMNSINTMVHSVLGTVSVASSDESSDSSTQPKRGASSSNSPYAS
PKTKLLEIGNKLNINVEFTDLIKQDNQYYTLVSLGLKPPHVCLGKAAVQAKSHEDAALRGLQLLRKSGILDKTGFAEESKQVEDVFEVGNEEQYRILDDFEEK

>*Dvv*StauC (NCBI: XP_050501881.1)
MKNQLVFFLHIIFITTVTSKTHSVKDLKNVFKDELCIEGNINSIYSKKTDTVVVKSTKTEGLLNLTSVGTVQESEKPVLSRLNELVGFNQIDYYFKLENEEGPPHNRLFT
VELTLGKEKYIGKGKSLKKAKQEAATKALEKTEYDFPEIKNNSPEDIEELTPTVLLNNIASKLGVGVTYYLLDKNKEEILHSNIIMTDKKKSYLQRLNASIYNDKKLRKKK
EIESTKGPFRLKLKFADYTFFTTSHSIQGGRHEVAVQALEYLVKNKDSLDIACLQEGSEAECKQNKDYLKSPISKVYEEAEKRKMSIECEVIKESGMSHKKKFKTECRL
GDIITEGEGFSKKESKRDAAINMLAKIAELEPLPIEEEARNFIKNDKKQKNRKKNKKNKLIKTKFDEIGMMLDKVGESIKTMTTNIFGEEDQPTDSNDSSGYDAKPK
EVKSNKKSHKQQPRTKQSSFQDELLELSNILGFGISYIDISEKDKHASLLSLYTNPEYICFGEGKTESDARNSAADNGLDLLEKIGIYDLFQEQKETNLERETKEGVRIA
MKHIVSEKKEEL

RBD1 RBD2 RBD3 RBD5

#### *Amphimedon queenslandica* Argonaute 2

>XP_019851058.1
MSRIGTPIDSSHWSPMDRYTSASLPPLPVPAHSLPPPVTLPPVTLGLPPFPGPRGGGGGPVIPTFPPPSTLPSPSITSSPLTPITPQTPSSITPGSILPWTPSSASLHPTL
SLQPPPRPNFGQIGRPIGLRANHFQVKIPTSTLYHYDVAIHPDKCPRRVNREIIEALIQTRKDYFEEQHPVFDGKKNLYSRKPLPGIGRDRVEITVTLGGDGNRERAFK
VSVKYVAQVNLALLDSVLRGESLAPIPFESIQALDVVMRHLPSMTYTPVGRSFFAPPEGEPYTLGNGREVWFGFHQSIRPSMWKMMMNIDVSATAFYKRQCVLD
FVHEVLDLDSDVIRRPLSDSQRLRFAKEIKGLKVEVTHTGPIRRKYRVCNVTRRPASAQTFPLQLENGDVFDCSVVQYFKEKYHIDLQYPFLPCLQVGQEKKHTYLPL
EVCDLVPGQRCIKKLSEMQTSRMIKATSRTAPDRETEINRLVARANFNADPYVQDFGISVDTKMVTVTGRVLPPPKLQYGGKARVQALPDRGVWDMRGKQFHF
GVEVSVWAIIIFTSVKQCPEEKLRNFVFQLRKISQDAGMPFRRDPQFVRYIQDRLGVERVEAVEPLFRQLLTEMEGLQLILVVLPGKTPVYAEVKRVGDTLLGVATQC
VQTRNVNRTSPQTLSNLCLKINVKLGGINSIIVPNMRPPIFREPVIFMGADVTHPPAGDEKKPSIAALVASMDAHPSRYSATVRIQQHRQELISELAAMVREMLIEF
YKSTRFKPQRIIFYRDGVSEGQFLQVLSHELASIRLACRKLEDGYQPGISFIVVQKRHHTRLFCSDDRDKVGKSGNIPAGTTVDVGITHPTEFDFFLCSHAGVQGTSR
PSHYHVLWDDNGFTADDLQCLTYQLCHTYVRCTRSVSYPAPAYYAHLVAFRARYHLQDREDRSSGDGSSASQQSEEPRSPALMAQAVKIHEGMCNVMYFA

#### *Nematostella vectensis* Argonaute 2 (NCBI)

>AGW15595.1
MPKKSKGRGRGRGRGNPHHEKQQTLVGQQATSRNNEHKQLPKPTQTQQETSLSQQPYTCSSAAEAQGPLTPPNNTQGGSLTNAEPAQADTLGQKFETQLNLSP
QSGKEQGAIPKTGARLKGNSLAPGSQGNGQFSSKNLLAQMQRTPRSASGSQNEASSKSQQAHNNQSQAAAHQQTQAGPQQTPARSQQTPAGPQQTQAGP
QQTPAGPQQTQAGPQQTQEGPQQTPARSQQTPAIIGSTTTADEHLARHRQENMELPKRPGYGREGRKIKLRANFFQVTLPNVDFLYHYDLEISPEKAPVSVCRDV
VDAAIKHGEFKGVFNGCKPAFDGRRNLYCREPLPLKSEEASLKVTLPGTDGGKERKFTLKIKEAGLVSIKELDQFLNGEFRGKVPQDAIQGMDIVLRQMPSMKFTA
VGRCFFPPPNGHCHDLGGGCELWTGFYQSVRPSQWKTMLLNIDVSSKGFQKSMPVIDFMLEILRQDRNRVLDTRWVMDERDKKKLTTEIKGLRVETTHIKRKFT
VMGLSHPAFNNRFRLEDNTETTVEAYFRNKYNTSLRYPHLPCLLVGQKKNSVPMEVCNMIPTQRKRLTDEQTAAMIRKTAKPANERQRDINQWVDELATASDQY
LKNEYGMRINKQMVAIEGRVLPAPELTLGGNPQGSALTPSDGAWDMRGKSFFEARTVEVWALVCFSHPKWCPKEKLEGFARQMGNVCRSEGMRMNPVPCRA
EYASRVQEVEGIFGKLLHDFNSLQLIVVALPDRGNKDVYNEVKRVGDTVLGIPTQCVQMKQFTMAKPQVCSNIAMKINGKLGGTNHVIADSLKATITDDKGNGIF
NSLVIIFGADVTHPAPGDMASLLSPPYVASLNRNASRYCARVRPQTHMKCKQAQEIIVDLADMVKELMIEFYKENKRKPVRIIFYRDGVSEGQFQQVLLEEVRAVQ
QACAMLEKDYQPLITFVAVQKRHHARLFAEEGRDARGKSRNVPAGTTVDTVICHPFEFDFYLCSHAGIQGTSRPTHYHVLYDDNGFSADKLQALTYQLCHTFARCT
RSVSMPAPAYYAHLAAFRARGLVNAGRGAANGSEESQMKTYYETIKVCKRMEASITLPKRNSRSTVRGENEPLSELVKAAKGIPVKSVEWQRSSSGKDRRVAKIVE
WQRASSGKGRRVAKGVEWQRASSGKGRRVA

### FASTA II

#### *Lymnaea staginalis* RNAi protein sequences – Putative homologues

**>Sid-1L/Chup-1L**
MLQKKHGDLEKHYGIPQHFGLFYAMGMALLMEGIMSACYHVCPNYSNFQFDTSFMYIIACLNMLKIYQFRHPDINAKAHTAYFSMAVIIFIAVLGVVYGTSVLWIF
YALIHMLMSLVLSAQVYYMGRWRFDRHIFKRLFRVILTEGRRCTRPIYPSRFILLLFGNLINWAFALYGAIKQPSDFATYLLAIFIGNLLLYCMFYIVMKLLYKESLSWLVI
LVILTSMVTWAGSLYFFFQNLTSWGDTPAGSRARNSECILMEFYDAHDVWHFLSAISLFFSFMILLLLDDDLSLKRRDKLPVF
**>Smg-2**
MKLMHGDELKLRYVGELQKPWSGVGHVIKIPDNHSDEIAIELKSNAGVPETCTHNFVLDFVWKSTSFDRMQSALKTFAVDETSVWGYIYHKLLGHEVEDIALKCT
LPKRFSAPGLPELNHSQVYAVKTVLQRPLSLIQGPPGTGKTVTSATVVYHMVKQNNGPVLVCAPSNIAVDQLTEKIHKTGLKVVRLCAKSREAIDSPVSFLALHNQIR
NMDTLPELNKLQQLKDETGELSSGDEKRYRSLKKQCEKELLQAADVICCTCVGAGDPRLAKLQFSSVLIDESTQATEPECMIPVILGCRQLILVGDHCQLGPVVMCK
KAARAGLSQSLFERLVVLGIRPIRLQVQYRMHPALSSFPSNIFYEGSLQNGVTAADRTRPGLDFPWPQPDKPMFFYSTFGTEEISSSGTSYLNRTEAANIEKLATKLLR
AGVKPEQIGIITPYEGQRAYQVQHMQYNGSLNKKLYMV
**>Smg-5**
MKKRAEPSNGSKSDTDRAKRVYRAALESIKRLDSINKEKKAYREVFRQDAVGLRNKLKDHCERLMFFNPIEYGRKAEEVLWRRVFYDVIQVVKHSKDHVRHHGSV
ETAYRTHLDAAAGYYQHLVFRLQREFSLCLAGVIDYHVLPDPRTGRRNDTSSIKAINPSVQEWAQRACHRCLICLGDISRYICDVDPMSIPTTADRYYHQAFMLFPEI
GMPHNQLGTLAGSRYSGCEAAYHYVRCLACEKPFDGARGNLTRLFEKNTKRFYELNRPQARDLPPDEQRHQDIKRFFTRFLKLLEIFHSSTNNIEMNELQQLCQHT
LQDFNLCMFYEPQHQSREGHQSGWQDSAEDEEDGEEFADPGDPSSFQYLDNGIVFKIVCCLICTIHLLEKSGSSNITAATAFLLALLSHILNHVVIRLQGSLEDMEH
PNKILIAGSSLSTENAHIEESSGSEDDSNQQNATKNGSTKTDSKKSRASTLRNLRRRRRRRQLSESSEEMSDLSEASDLSEGNEDAEDQIVSSDSEDDALASYFDHGS
DSDLSEAEDAADNDAKANTNNANSGTPDEHSKWPPPGNSLPNGDIKHIWSNGTGDKSLAHISSELFSSSLMFLGQDLAHNSQVPDVLHDQEDVPIPPGFNCSE
EAKHVAEITEKLANFDIETDTETFMPTDTEQSGTVTETDEADNEQSTSSNTEKQEQEQQRLQQTLEVVHSQGLLSTVKVMCDWMRTQTAIISVCAQSSYSLWQRL
SVLLNFLPHENEIVQHGMCWVQELQVIATQTRFADWAQVFPLTEDIEMCNLPPLVDIHSKIDFTTRHRAQLSDMQETCLRICCLRRFGYFLTTIENIEFTYQPETAM
FFGPSIDSNISSNSHDTMTKMVSNIIDRTFQYISLKTFSFRPPKLLL
**>Smg-6**
MAAEEEIVRISYANLEKIVHSSNSGTSDSKGRNERRQGLNRQQRSKRPEQQIYRPGQLRQSKVCKSGERSGDIGRDDDDSLAGADAWVGGELQVNSTDEDKGN
GSRESSGDITYDDINQRLDNLYIESPPREIDSKTKRDTNDTLSAHDDDGVCSTSTHSPDDKSWTADREKDGGKRRGKRPDIKIYVPRARLGDREKKSEEKDDVFESA
SPSRSNDCRDKGFDKMLPEQLDEIVVSDSADSLRKVQVKTKGGAMKVTVMNESLTSGGENSPTLPRKELGPVRGKNYQKPVQNTRASPGPDANLKQKNKDFDN
KAIVPPRGTEKFSRQFESSKKGKGKHEFGHNEVTVSQDIRKNFPQKVERSVKSLNKNRRSNSSDSRPKDHGKSEANSCQSDKGQFSSETFPRTKVPNNKSSKTGGK
DTFTQSKNGKDTYGTNKGPVREGEKFYWGMRRQRTGSISSEASIASNPSNGSFYSGSSDFTEDDEPEHILNWGEEVEKAHLEELARLVHDGAKKLTNSLDSYPGF
GPTEGNAGTEKKIEVPSTLDRQKSVSADEIYHGKAKEGGRTRHRRHRHRKNSSRQGSRDSSVHSSIHGEDGFSERGHRRRRRQRRNSQGSQGGYEHHHNRLQQ
KQQQQQHEDRHYAESGGDSGRNNRSGLQVTIGDHNRHVEISRSGGPRQRHRSDEKWDPNNVRDYDVEEGEEDWERNYDYGEDREFKRDDRHGQQKWEKE
HPPRFKREGVGAGRSRRGSVRDEIQRQQSGREKEERNRHEEHGNNEERGELKQEKDFDQRHSGDQRAVHPAHDASAGGRSCQRGGRGQGRNVNKAKSRRR
DERDENCNQHEQRHPIQRAGSLNHEDTAASQKARGGGLLRLPSQPEPPRPHSNPSPASHADHAFHHHADNFGGGGPSSSAQGSNTRGPAQGKRHLFDPKNPS
KPIMVPDAAPRFEDTDKGSSPGAVSGHIPSSPDSPQTLPYPQVFYPTEFMHPMYRFRPPMAPFPPFMGGYPPGSIPPHVFYRFPRMTPPDMGFYNSPAEMDSE
GLMSSSSRAQCRMMAEQILRDSVPFESQLGNILSRRPNSEDSFRMVNQIRAELQNRLEQVMLMDIEVANKHNVEQTLWKSVYYQVIESHRKRIVEDSKDFVSKQ
RLIDVLDEGTRFFENVLKKLQSTFGFDLEVFSDGNKPVPENVGRNVKFALLSAQRVMMFLGDISRYKEQASDTTNYGKARHWYLKAQRIHPRNGRPYNQLAILAV
YTRRKLDAVYYYMRSLAASNPILTARESLMSLFDEVRRKVDAAEKKRFEDRRRRMALRRKPQIRGPRVEIWVQPDGTSTQDRVEEGEEDDDLSSLSVFELNKRFVL
TYLNVHGKLFTKINFEMFAESSSLMLQEYQILIQHSPCVFSSTRLLQLMVINMFSVDNTALK
**>Eri-1**
MIIMDFKVEHAEENENKNFAEWKEEPHNSCWGASAINYFVDPSTEQIAELEGYSLKIETSGVAHRPKARLDSGTGGGANSDEILAKLSRINGAINKMTKEQMQSK
LAEYRLNTSGVKDVLKKRLKNYYKRQKMAQCNRALTDDEKMKYDYLVVIDFEATCSETNENFIHEIIEFPAVLVDAQNQAVCNVFQRFCKPRVNPTLTDFCTSLTGIS
QQQVDKAKDFTEVFKEFEEWMAIHKLGTDYQFAILTDGPWDMARFLRTQTELSGIAFPKWAKQWINLRKAYTAFYGCGRINLHQMLEDLGMKFQGRPHCGLD
DARNIAAIAIRLLQDGCIMRINEHFREGQISPQKSTGSSRASHSRGHKSPPKSHTNKKYIQRDSVQKCKEPFEVEQEGENMHELFEYLKLQRS
**>Xrn-1**
MGVPKFYRWISERYPCLSEVVKEFQLPEFDNLYLDMNGVIHVCSHPDDENPHFRITEEKIFKDICHYIDFLFRMIKPKKVFFMAVDGVAPRAKMNQQRGRRFRSA
REAELLIQKAQERGETLPTEQRFDSNCITPGTPFMVRLQDHLKYFVVDKITHDPLWQGPKVYLSGHETPGEGEHKIMDYIRYSKSQSDHDPDTRHCLYGLDADLIM
LGLTSHEPHFSLLREEVRFGGRKDKNKRPVTPEETTFHLLHLSLMRDYLSYEFSSLKGKLPFEWNLENIIDDWILMGFLVGNDFIPHLPHLHIHADALPLLWRTYMN
VLPQLDGYINEAGHLNLPRFEKYMAELSKFDEDTFESAFSDLRYLESKLGGKSINESAGASRQAGKLKKLKPFIVKQDGVPFAALEDESKNNVLDVLPDEDSGNTDD
DSDADDEQEDDDEDLSDEDEEDYNTFGDEFRLHKRDYYMNKMSYTEVTPNVLEEQAHGYVIAIQWILLYYFDGCPSWSWFYPHHYAPYISDVRGFSDMTITFDL
SKPFLPFHQLMAVLPAASKELLPLPLQLLMTDSNSPIVDFYPVNFETDLNGKQQDWEAVVLIPFIEEKRLLETIASKEHLLTKEERERNRHGPHFLYYFTPDNQGFYPS
SLPGVFPDLHLTRAKVTELDKDMFRIDVNRLRKGLLDSVRLHVYFPGFPKLRFLPHTHELAKCGCKVFQHNSRGENMLLRIQSKEGLKLEEIADELLGKETYVGWPH
LHEAKVIAVANETFKIELLEEFNKKGKKEAAAISFSRKKLTAAESAVVGREAISIKERHMDRFAIDVGNTEIVIYASPLTGKMYTYGALGTVTLEKQFASNVAPYLHQMT
VKDLEVDDKGPELLKTLDEVFPPGCDIFMIGTPHYGAQGKVKEILHEHGRIRICLDVKEEPDFRCLGNSITSEKYLNNYQAAQQIGIDGHIMSRITGSVFVEKSGPHG
GKANIGLNLKFTKREEEVPGYTKKLDEGWVYSFKCVKAVEEYIDRFPDLWLSIQQQKSNSDIYTEAKLFPEGSSSKLSDVIKYLEDLPCTKAPRMKFGAEILSEDRVKE
IEEMTAALNPVPDIVKMKVKPRLLFRPMANQGSLVPDPDADFFLYDRIVVVKAGYSVPFGKRGTIIGIPKEDDGGITPHSLYDVVFDEPFLGGINLRCSQNKGYRVP
GSAMLNLTFGEFRKSVSLDQRAALSRNSARMQQTHSSERSSYNGHGSGFSSSQNNNNAMGQRQPYGDRGGGAYSDRTALWKQQQQQYSSRSFQNSQFHQQ
PVRWENGQPTFGQVFGFQSQGAGKSQGVTPPKFVTPKVQGPQTVMHNTTTTQARPCHLLKS
**>Xrn-2**
MGVPAFFRWLSRKYTSLIVNCVEEKAREVDGVKIPVDTSKPNPNEVEFDNLYLDMNGIIHPCCHPEDRPAPKNEDEMMILIFDFIDRIFSIVRPRRVLYMAIDGVAP
RAKMNQQRSRRFRASKETGEKIEDLARVREEVLSKGMYLPPEKPKEEHFDSNCITPGTPFMFKLAECLRYYVHDRINNDPGWENIKVILSDANVPGEGEHKIMDYI
RRQRAQPDHDPNTHHCLCGADADLIMLGLATHEPNFTIIREEFKPNQPRPCDLCGQMGHELKNCEGMSREQSDVPMAVPGVEQQFIFVRLSVLREYLQTELAM

PGLPFTYDFERALDDWVFMCFFVGNDFLPHLPSLEIREGAIDRLVKLYKNVVMKTGGYLTDSGVVNLSRVQLVMSDLGKVEDEIFKKRRETELDFRRRDKQKKQR
MKQQQNAQRPKWMMGGQFAPQALGKPGSLNAVANPRQAAFESRKQNMSTSGGGGAAAELKSLMQKECFQVGDKRKAEEQGGRGGQDSDEEEEAPDDV
RLWEDGWKDRYYKSKFDVDADDQEFRYTVAKHYVRGLCWVLRYYYQGCASWKWYFPYHYAPFASDFVNIGDLPNDFEKNTQPFKPLEQLMGVFPAASKKHLPI
TWRTLMEDPESPIIDFYPTDFKIDLNGKKYAWQGVALLPFVDETRLHKALETVYPDLTAQERKRNTKGSDRLFIGVKHPSYEFLEGLYEGDLSSNPVPIDPKLTNGMA
GHIWCDEYNIVKGGVVPTPLPDVPDILDNKCITVCFRDLQFDADFLFKAVVLPGAVMPPPTLKPDDWNNRNPGQRYRPQLGFQQNSGYRNNKDMSSAVRMIR
NTAGMGGQGYVMSSQSMMPSVRPSLMGAVPFDYNNLRTLQRPRYNGSQGAGYNSSQSTSHRSGGYGSTQNSTQRSGYGSGHQGSTHQRGLFQLFRFFPRTI
HPSRHMKRDGIQVVLWL

**>Adr-1/Adar-1**

MSQVRSPYRERSTRSRGRGQGTKLKRAPIQDKEVTKGTSKKNLEQQITNFLEGKKEPQRTIDIARAVGLVRAKDVNPTLYDMLRRGAVNQNAHQPPLWSCQHSY
RSAGFGQPYNDFQQQDMVPYWVGPRMDMSVPQGMQGWGNIDYQQQDMGTHWAGPRMDMNVRQGKQRWGNDYLRGNGRPNGLYPTPMESGNHT
RRPHSRAGSRNFTPQQRNTTIESRAGSYHQKNPTQHQDLFKKTGYDHNAPGAYRRNSKSQDNRYNDRMRKNSPKYPSKSNNTEGQTAPAASVESSRQEGDNE
SHSTPDHTDPPVAGPSAREELTLASGESLTRSPTVEETILKESGATEEKNSGGNSPEEYYESDSDEDIDYNRELEDRLTDVTNNDVRTQWRELGYELSDGEHWEDED
DFENDEEMWDEDYQGHLEHPSRRQGLGDSGWSSGKQDQLVPRLLEVLRSVPMNMTTKMDLLSRLNVGQENLEDALKECENRKLITCKNGFTYQLTEGGMQQ
LGDLGMPVDEARDNTEQLDRPSRDNTEQFDRPLYPAELIKSETFFAGKPEGMVTPGYGRGASINRAGLQSSSQETNEAASASYRDRSPLVSPVEGLHSKMPWLSV
KTSPPAPESLYPGTHLPQYMSTPQLHLGPASSSTIAIGQQLIKDKKTDIPGMEFLILCDNKQVGESSSTQDSQKQQRRPPLLTTPLVRGSNVSPSMTGAGSNRYPYM
TYINQPPTSQPYFNQKRLSPTFQQPMSLQSLPSKLQITAESFAALNKNPISAFMEYSQSRHLTATIEVVDQHGPSHKPVFTFSAKIGNRAFPYVSCSNKKDGKKEAAE
QAIRILIAEGQFHLPQQIGVMTMPEANMTDHDKFAAKVHQTYNQLIATVPESFPGRKVIAGIVMQTDTVDDSQVVAIGSGNRCITGDKLSQEGNTVNDCHAEIIT
RRGLIGFLWDHVIDYVKAKDDSILEPSPKTGKLRVKKNIKFHLYISTAPCGDGALFSPRDVISNNAPIPDGNKHTPIYTNNAHGVLRTKMEGGEGTIPIEADYKEQAW
DGLITGERLRTMSCSDKICRWNVLGLQGALLSHFLDPIYLASITLGYLFDQSHLSRAVCCRLDRGEPSLKVQVSKPYTLNHPTLGRVTACDPPRETQKTKSYSINWRL
GEDRPEVLDGSLGICYSALEQKMFSRLSKKNMFDKYKKACALYNWKDLPRGKTYHETKQMSKKFLQAKEAMFQKFKATGCGKWISKPREEEMFS

**>Adr-2/Adar-2**

MVLNEIHRGVKFELVSEVGERNFRLFNMKVTVGSRDFMGSGRSKKLAKYNAALNALKVLHGINHFERRDMTAVNQESIQGDPQVLADHVAQLVFNKFGELTDN
FTTPYARRKVLAGIVMTTGASGDPGQVICVGTGTKCINGEYLSAEGKGVNDCHAEVICRRLLLRYLYDQLALHLSPATAADQSILVEMPEGGYQLKEGIKFHLYISTAP
CGDTRIFSPHEKSNSGKKVKEPSESATVAMDTAGDNSAAELEGCSEKPPVKKEQDNGAANSGIDLEGVADKHPNRRARGLLRTKIESGEGTIPVKVSGTIQTWDG
VLQGERLLTMSCSDKITKWNVVGLQGSLLSHFIKPVYLSSIIVGSWYHSDHMVRGVYGRVSHMKLPSPTYRLNKPFLSGISSPECRQPGKTPNFSVNWFIGEENME
VINPTTGKTDTGGISRLCKQAMFRNFLNLYGKIGSLSSQGVVDAPRLYSEAKAAVMEYQLTKQSLISAFQQANLGVWVKKPHEHDEFELVSN

**>Lin-15b**

MNEFSTSVLQFTHLDQSKGSKNIGVFLPAEEVKLKRTRVPKQPPMMSDGVSPTLPKCNKGPKRPRSVVWYYFDKILHEPQRARCKICGATCHHANNTSNLFKHLR
VKHPSTYKEAESQREAEMELYLELKAKSGKPFIRPGIRGRRPKSLTAITAPIVKAEESVMMNTSARETVVVRKSYGAEKVKQTMTRALLKMITVDLHTASIVNGKGFR
EFVRSLDARYEIPSKKNIVKNLLPELAEETKHKIKLDVVIASYFALTVETWTYRESQIFMTLSAHFIKDSWEMSSIVLETFDCTEDKTETNVATSFKRITDEWGITDRVIA
LVADNSEDVIHSASLVNGWEELICFGHTINQVISHALTSVAELVRIQKKTSDAVAYFEKSIKASDTLAAVQQQHSLPVHKLKQENPRFWTSTYAMFERMVEQYEAVN
TVLCFLGKDHMCLCDDEVELIRSVVEVLKPFEAATKELCTEQYTYMSKVIPIATLLQQVTSVGMATLPGPQSALKNALISQMQQHFTSVENNYKLAVSTLLDPRFKR
HAFTDATALELSHQRLLSELATVVQISDTPSLPSDPSYANDTAGSFWNMFDRKVSEAGAMKTSASEAENETRRYFKEANIPRNANPLNWWKLNEVQFPHLKVLA
QKYLCIPATSAPSNRLFSKEGDLTAFKREQIKPGNLHTVLFLNKNLA

**>Eri-7**

MNEFSTSVLQFTHLDQSKGSKNIGVFLPAEEVKLKRTRVPKQPPMMSDGVSPTLPKCNKGPKRPRSVVWYYFDKILHEPQRARCKICGATCHHANNTSNLFKHLR
VKHPSTYKEAESQREAEMELYLELKAKSGKPFIRPGIRGRRPKSLTAITAPIVKAEESVMMNTSARETVVVRKSYGAEKVKQTMTRALLKMITVDLHTASIVNGKGFR
EFVRSLDARYEIPSKKNIVKNLLPELAEETKHKIKLDVVIASYFALTVETWTYRESQIFMTLSAHFIKDSWEMSSIVLETFDCTEDKTETNVATSFKRITDEWGITDRVIA
LVADNSEDVIHSASLVNGWEELICFGHTINQVISHALTSVAELVRIQKKTSDAVAYFEKSIKASDTLAAVQQQHSLPVHKLKQENPRFWTSTYAMFERMVEQYEAVN
TVLCFLGKDHMCLCDDEVELIRSVVEVLKPFEAATKELCTEQYTYMSKVIPIATLLQQVTSVGMATLPGPQSALKNALISQMQQHFTSVENNYKLAVSTLLDPRFKR
HAFTDATALELSHQRLLSELATVVQISDTPSLPSDPSYANDTAGSFWNMFDRKVSEAGAMKTSASEAENETRRYFKEANIPRNANPLNWWKLNEVQFPHLKVLA
QKYLCIPATSAPSNRLFSKEGDLTAFKREQIKPGNLHTVLFLNKNLA

**>Drh-1**

MAHSNSTGPRKGSRYHRLDDIPTDTFTPRTYQIQLLDSALKRNTLLCLGCSESKLFLALMLAKELASPTRLQRSQGGKRTFYLTDTDDDVEMLYQTLPHHTDLQVEF
CYPHSLLEDDTQDEISGTQMSLQKWCLLVDSSSILVCSGHLFLTALCKNYIDLKEVNLLVFDNCHRAVEVENHPYKEIVNGIKSLPQKSDQPHILGVTASIAGTDCSDP
LKLQQTISEMEEAMFSYAETSMLILSERFGCRPKESVVQCLDLELDDEQEVLKDTLEEMLEQNYYFFSDCRIAVDETVDNRDPRETPMQVLSQCLNILNVLGSWCT
ASISEYFVMQLEKIVKCEKNEIHKKFLRCVITTLRVLIKEFEKQFHPDYHVNELLEYTTPKVRELVKVLRKYKPEVDFIIVSNGDSFNDDEEFSDLSDEDDDSIMSDSDG
EDNSGEPFNGSSKPMHIAVKRTADGGHERIMTFSEEEEKNLCGIVFVENRYIAFGLNKVIEEVCSWDENLCFVKSCHVTGQGLRSANGKPRVNRTYKRQEEALRKF
RTQECNLVIATLELEEGIDIPKCNLVIRFDPPKDFRAYALSKGRARARDAAYVILLDDENFEGFNTSLKVFKGIEQVLLGENRSLQAIRSNIPGEDADNEYDSKDDDDSD
ASSDERDNLPPYCTMPHSLSSPHVTLQGAIALINRYCAKLPSDAFTHLTPQCRIEKLDGYAPSYIANLRLPINSPVKEELKGPAMRTKKLAKMAVALKMCEILHKKGEL
DENLVPVGKEMFLYEEEEEWVNEEDLAGQARPGTTKRKQYYIKKTTDALLQCRPKPGQPCFLYLINLKLTGPITDEQNTRGRVIYAPEDTVQTLGILLSKKIPLMPYFP
VYTRSGEVTASIDLQQTDLICSEDEINRLLNFHHFVFSGVLHLEKDPMEFDCVNADCGYIIVPLNGKGCEELTIEWSFVDKVIDFIKVEKRAGFSKKSGDESFQFCLDD
FQDAVVMPSYRNIDQPQHFYVAEIRHDLNPLSPFPSPELYKTFQDYYTTKYGLMITNTEQPLLDVDHTSARLNLLTPRYMNQKGVALPTSSAETKRARRENLQQKQ
ILVPELCDVHVFPASLWRKTVCLPAILYRANYLLLADELRRKISRETGIGVEELPPGFRFSKLDFGFDTSPEKLMDAEVNCNKKISSNIKEESAERSPDPVLCCMEKDSS
NNRMIKESLVNDTDKAKDESSTSSDSDSGIASSSSDNNIDITPSHKILIKSSEHLEHGSVNPSKKFQELCGHDNNVVLNTQSWCLDHNNILYNVTPLGSSFPGTNQCNI
ISNTEAVELPSTSSAVPQQDSNGSTTTSTVINFSCTSAPAKVSPDNATVGATISIMAPPSIGVVSSCEPDKPFDSTDSEPLAAETLPRQAIHGDKTSVALSLPCSTTSSS
TLLTSSAPLESPSAATSTWSMPITSTTFGEKGHSSDYPPLFVSASEVTNASTINKACYLSPRVLTNFNSDAASNRANCQADLFITTDINGCELNKGLNDASPALLATET
SPAIDHSHPNHCSCINHPCVHHSDAQIISADQKHIHAIYNNVSHNNAESSSNSAKITSGKAKSHINGHVKQDIKYSVAQRTINEFCNQDKTESKGHAKAKCQLSSD
GPAIIMNSLSQELSDLDIDLIVPQADTPDTFHSFEDSTRSGANTRKDNQHKKTTVRLSAPANSEWPKGQQTTTVDWPDVQQTGNSERLNCQQFTLHEQRKPVH
KGFDKISVDKGFDKISVDKDETGSANVTTDNYNATAAAEDIHVIKFSFNTDEPGVKCKEAKEPSPPDNKKESSPTDSKKECNPPDSKKELPNENTENRLTFDLTRVKS
PLETQQRNTELEIGVWKSSEKRIYDKHQAADSHREKITSLDHSSYNEMTETNHSLCQPTFSCVTTGSLTSISLDEDRDPQTFVGPSPCLIMQALTMSNANDFFSLER
LETIGDSFLKYAITVYLYCSYPGIHEGKLSYLRSKQVSNYNLYRLGRRKGLAECMVSTKFEPYENWLPPGYVINDERRRGPVPKVVIVTPGSKINNSLRNFYLDETADA

AKRDEAILFNKELEWIQQSQEADQQEEVQEPNNDMTPYSLQCHHGLPDKSVADCVEALIGCYLTTCGRKAALIFMSWLGLRVLPKKKKLILDVSKIEEHEKSSVGK
LSEQEFDELRCPPSPMFSNTPANVAKLNHLLQGFDSLEEKICYKFRDRSYLLQAFTHASYHYNTITDCYQRVEFLGDAILDYVITRHLYEDSQKYSPGILTDLRSALVNN
NIFAALAVKWGFHKYFKAISPSLFQVIDKFVKRQKEKKEDDIDIDEEFRELSLEENEPEENTEEDDEEEEEVEMEIPKALGDIFESVAGAIYLDSGMSLDTVWRVYYR
MMKPHIDKYLKSIPKSPVRELLETEPETAKFERPERTMNGKVRVTVNVVGKGVFSGVGRNYRIAKSAAAKKALRSIRTLNQNLMN

**>Drh-3**

MASERDESISELFATIILSEDECNMPIWEHILHHNVVDSSLVNVIDSQQEQQIVKPDDENQEEYFSDQDFPSKDVPRATVVSGIPSLELRNYQRELAEKALQGLNTVI
CAPTGSGKTRVATHIILEHLKSRQDKKKVAFLARTVPLTRQQYKTLRKYLPNEFKVTYITGQCEDSMSLKMLIKHHDVFVMTPMVLMNNLQRTGLRLEKFTLIVFDE
CHHTRKDEPYNLLMFEYQKLKHGGRDKPRSNLPQIVGLTASIGFEKSRGAQGSVLELLGNLDAPYLSTVKKEKSELQAIVPVPNEYFKELTAVQPDECFQMIITNMD
KLEKAISKHAEGMKNNKLQQLMAFVPKNKRTQQYEQWAVKLKMAINNFTADSQLKENKLSIMVSTTISDFLTAYNFALQTYDLVESRDMLTYLTKFFEKYLQKED
MTEEERIFLTDFEALRLNILRGQDKRNPNLRILADTLKTYLVNRGSGSRCIIFVKTRALALALTSWLNRCDIPEIRDLKASMFTGVNASGDKGGMTPANQEQILHNFS
SGQTKALVATSVAEEGLDIPECNLVIKYNHIGNEITSVQTRGRSRKLGGVSILLAMADVVKKEMLNHENAKIIDKATDEVATIEISVIKQHIENQQLEILRASEEAEVRA
AVRRRQSQSVQFKMRCSRCRSLEIIGADLRTIYGSYRVALDRNLLDDKHIKCRPTDVKLLDGLEIFGDVICCGEGKPGMTCGQKLGKMIKYSSIPYFATSCDKFVFCS
GDKTENFKQWKKASEVYFIDELTKEDIRRYIQMGNEEVKEFGDSDEGSDSE

**>Drsh-1**

MFPWLEEGGLTTYRMALVQNQHLAVLAKKLKLQDFMLYVHGPDLCHESDLQHAMANCCEALMGALFLDGG

**>Xpo-1**

MRMKMATLTEQAEKLLDFTQKLDIQLLDNIVSCMYTADGQQQRMAQEVLTTLKEHPDAWTRVDTILEFSNNQQTKYYALQILENVIKTRWKVLPRAQCEGIKKYI
VGLIIKTSSDALLLEREKVYVGKLNMILVQILKYEWPRNWPTFISDIIGASKTNESLCQNNMAILKLLSEEVFDFSSGQMTQAKAKHLKDTMCSEFSQIFQLCQFVM
DNSQNAPLVGATLETLLRFLNWIPLGYIFETKLITTLIYKFLNVPMFRNITMKCLTEIAGVQASQSYDEQFLQLFTLGMNQLKQMLPLETNIKEAYQGGTDDEQNFIQ
NLSLFLCTFLKEHAQLIEKKQELNILLMEALHYLILISHVEEVEIFKICLEYWSSLASELYRENPFSDSSSTPLIFSQMRSQTQEMPVRRQLYNPVLSKVRLVMISRMAKP
EEVLVVENDQGEVVREFMKDTDSINLYKNMRETLVYLTHLDYTDTENIMTEKLHNQVNGTEWSWKNLNTICWAIGSISGAMHEDDEKRFLVTVIKDLLGLCEQK
RGKDNKAIIASNIMYVVGQYPRFLRAHWRFLKTVVNKLFEFMHETHDGVQDMACDTFIKIAQKCRRHFVQVQVGEVMPFIEEILNGINTIICDLQPQQVHTFYEA
VGLMIGAQVDQVAQEHLIERYMLLPNQVWDGIINQATHNVQVLKDPEAVKQLANILKTNVRACKALGHPYVVQLGRIYLDMLNVYKVMSENISSAIATNGESVT
KQPLIRSMRTVKKETLKLISGWVNRSNDPQMVADNFILPLLEAVLLDYQRNVPAAREPEVLSTMATIVNSLKSHITKDIPQIFDAVFECTLTMINKDFEEFPEHRTNF
FLLLQAVTQHSFQALLNIPPAQFKLVLDSIIWAFKHTMRNVADTGLDILYVLLQNISREEGTAQSFYQTYFTDILQHLFSVITDSSHTAGLTMQATILAYMFNLLEHGKI
TVQLNPNMPAAQNIVYVQEFLLSLLKTAFPHLNEQQIKIFIEGLFSFDQECNALKEHLRDFLVQIREFAGEDMDDLFLEERENAIRNAQDEKRKQQLAVPGIVGPHE
VPEEMQD

**>Xpo-2**

MLKSPEQVQKQLSDAISIIGREDFPKKWPGLLSEMISKFETGDFNIINGVLRTAHSIFKRYRHEFKSQELWEEIKFVLDNFATALTELFKATMDLATKHGNDPKALKVIF
GSILLICKIFYSLNFQDLPEHFEDNMKVWMDYFLALLSADNKLLQTNDEAEAGLLEQIKSQVCDNVALYAQKYDEEFSPHLPGFVTAIWGLLISTGQGVKYDLLVSN
AIQFLASVAERPAYKGLFDSVDTLASICEKVVVPNMQLRNADEELFEDNPEEYMRRDIEGSDVDTRRRAACDLVQALCKSFEGPVIQNFSRYVQALLEEYVKNPNA
NWKSKDAAIFLVTSLAAKGQTQKQGVTQTSELVNITEFYQAHILSDLKSSNGNFCSWYFDAIY

**>Dcr-1**

MEEFEDLQTITESLADTFKKYLNPQAVVEKWKFKEKESELVFLSDLENSREESVVKLLNWVHENKINGEPALNLFVQTLIELSEDSAGANSTAYVNSHHSTNAINGQ
KVELKKCVGYILQLVNRKWTQKMSDTEWQEIIDVMKSHKFIMEKDARTCKEKTEPYEMCLELIHSIKRNKPEWPCDFINAIVERNPELVEMQQDVKANQFGNVR
FKTVRLELPMHTDVNFPDDDSTVSALSSNTLPSMSDDWENEINASSHLDSRRREKSKFHLSMEMKNSPGVQMVISMGALNDDQITTEPQDKKAKFISHSQSFS
DEEFESEEADQNEKEQENAHSNEQPSLNLRQYQLELAANALKGRNTIICAPTGSGKTRVAMHIILEHLKGQKENEPKRKVAFLARTVPLVMQQYKSIGKYLPKPYE
VTNLTGESEDSMHLHMILPDNDVIVMTPRILENHFHRNCLPNLGVFSMLIFDECHHTRKGEPYNTLMYSYLKTKKTNPEIKLPQIVGLTASISVEKAVQDEEAVKCIL
KVCGNLDVDSISMVRQYEVELKETVPVPEEKMIKLHERDNDQPVEKILSIMEKLEAIIMYHAKEIRNQELNSFVQKIPTDKKSQQYGQWAVKIKNLAKSLPRNEEKE
TNLSVRYIIIVSDYLMAYNVALETHDLVQLRDVLFYLEKYFVKYRENEKRTSGESTCYTLFEGLKQVLKKRGDDENPNLTLLKHTLLEHLVGKGEKSCGIIFVRTRALTDA
LTSWLQRCGNKDLQRLKATVFTGTAASVDQGGMTQTEQEEIIRRFRTGDVRLLVATSVGEEGLDIPECNLVIKYNHVGNEVTTVQTRGRSRKEGGMSILLAMDKIL
RKERINQEKAKMMTRALQKIASMNRKFIRKSNQAHQEQIMKEEEILEYVKEQEESRLQRKPFKMVCHLCRKLVISSKDIKTILGTHRVVVDRNILKLVKIDPYKDGK
YFDEVQMIGSVRCMAEPEPGKHCTNKLGSMMLYARVPFLVLGIKYFGFDVNSKIGLEFYNQWKKVPYFIDEIEPEDIQRYLPEKTIETNPGNDDVEDDDDDDDDD
DGDGDSDDDEGNKPPELESPVKNIQTHHFNVQDMSSTKFKHSVETQKSVRSDDGQSDTDSGRYSDKASISSATSTTELKQVIPDVRHQLLLGPEEPNNVRRAFVT
GYSNPDKKPAVQITQQSSELLSSFKEIQPYFSEFSDISALVPPTNFSTESHIGAPLVSDCSDISVQKLEDRNSGENSSGGPGQSTPANEMPSLDSIQVSLESDTRDSSPT
APDSSAGPTSYIGPGVGAVADPVNHIGPVLRSLESNDGDRSHSSSVDDNYQNQKPVNQIPEVKGDGEDSVGNSPHSQADDVLQISQDETDDDETDVKKPLPGY
GAGQQRRDGDNDDPPSQKGSSMMSTLL

**>Pasha-1/DGCR8**

MFLQKKKICFPFYHSILRSWRERRKHLGERNRRSRPELPPSTKLITCPIPGNPKADGNGDNVKKREFILNPTGKSYLCILHEYMQRTLKIQPLYVFKELENSKTPYGAT
VMINNIEYGTGYASSKKVAKQEAAKETLKVLIPDLFKKITDQEIKRNISDLSFFDDVKVTDPRVNELGNKVGQPSPFQLLLECLRRNFGMGNTQCEVSTKPLKNQKC
EFTINVGKHSATVVAKNKREGKQLAAQAILAKLHPHVPSWGSLLRLYGTSVEKPIQKPEEMNDVKSHVPNHSVLESLRQEMRKLHRQKEAIQSKGKIIISSKDLPCK
MSGVDL-MFLTLRPCQTLAADVE

**>Xpo-3**

VLDSAGIFKSEGEDSEFLCKLSKLINGIGLGLISSWQKLSKNGVCKNSETTLRALEAKVAYMLRFLGDDDDDVSSAVIEFATEYIAMMKTLPSMSVVQKGHIEGLLFTII
KKLKYDESYQFEKEGEEEAEFQEYRKQVRVIFNNIASLDTQMILLRIHELVSQTLPRWESLDFRDVEIAITLLYSLGEAIPASHGQHFSGNPEKASALQDMMRTLITSQ
VSRQGHFAVLLQFFETVVRYDRFFVCEPQHIPEVLMAFLDERGLRNSSPKVRSRVAYLFLRFVKSTKTHMTPYLEDVLKQMQDLLTVNSPDNGLSHSLSDSDQLFIY
ETASTLVVSSSVEPEKKQVLMKQLLMPIATKFNVMLTKLTEEQDEKRQQAYADCIHNAIALASRASKGFSGQQTMKQCGCVSAFTELLSIFLAALEVTNQRASLQQG
VRQYLHRMVVCLEEEIMPFLPEAMERLLKQPDARELYDFLPLVNQIIMKFKANISPFLTQVFMPLVMTIYQVLSAPSDHNDHCSAEEKKLLQRGYFLFLSTIVCNCLD
VLKAQSMENLNQVLMSLIQGAVELPDPQSQKCCFSTLKKLIEVWGCKEGLPGFPAFIYKEIVPACFMAPMKQTFDLADGQTTLVLGECAACMQEVLQQRGQEFL
DFLKGDFFPSRGINQESGQCYITNLQTDLKSFRSWLKKFFLQAKS

**>Mut-7**
MLQGLGSQLRSCGVDTFIMGSSQRHSDAIQISLHDNRIVLSTGKAYEQLRARLGDAMIYCVRHDVSAMDQCFDVLRHFNVKVTKDDIFSRCSMCNNADFVTLPS
KEMEKLYMYKSQQSRFAAGGQRDPNVSLYDVDQGTSSTFSQYGIDPASIAFLHNGVKIQVETVPKVESFSTIQEFFVCTNCGKVFWVGSHYSNICTQFDKVLRLGL
DLDDPRPEELFDEDGQEEDYGYNCDGAEAVDQWGDDYN
**>Cid-1**
MDSILKSIPPPSISQLDALSASLDQVYEQQGIDEWEQKSRQEIVNRLESSIQERIPEVLMFMYGSSLTGFGLKSADINVDLNSSDKKMKMTYLLKEIYLILKDRTDTGF
SSVRSDFSAKVPALLMLDDATGLLVNIAIHCYSAHCSGELLSIYSDFDIRVRKLAVAFRYWAHICGLDRQSDGFFPPQALNLMVIYYLQVVSPQLVPIINPPKVSGEDI
GDFRKDRPAFEKMRAKVMKYSMEKRNSMGLGELWLGLLRFYSLDFDVSNTVVSIRTNVDLSRNTKPFNSRKLAVEDPYMPKKNITRMVNNSRIYEYWQDSIRK
AFYYFGLPRNSQGHSLISEEALRERSLKEKSEASTSGAKPEPKTSYTKGTAQQASSGGTEKPKAQSETEAVISLSGSDALANSQSVKIPPNPLIASSSKVPEANKQPSP
QQRTAGDLAVTTLTPDKQSSSKLKNKDLDLPSADQPDNTAVKNKHGDNSNVLPVNHPVQNNGKEVSHFAQGYAVDIIGKAFDQLTVQTSDSDKDSSKDSANASL
DEGNASSESDEVSSHSEASCDKSEKYSPSSAEEQCKYEFSEENFMTDGKGPTLVCTYCEQEGHLKNACPEDELPEVLPLPPLTPMHIKVLSETLNKVPSEVGLNDESF
GDRVIFLQNLEGYIQQTFDDAQLALFGSSCNGFGFDKSDMDICMTFSRRSGKFDKIFVIETLARRLKQNMDLTAVQAITTAKVPIVKFTVKKNNLEGDISLYNTLAQ
QNTKLLYCYSKIDPRVRILGYSIKTFAKVCDIGDASRGSLSSYAYILMMLYYLQQVRPAVIPVLQELYEGKTRPSLEIEGCEAWFMDDLSKLDKFWPEKGQNKMTVAEL
WVGFLRFYVEEFNYKELVVCIRQKEPLTRFEKLWNGTSIAIEDPFDLTHNLGSGLTRKMNNFIFKTFINGRMLYGSPIDDCMEIFKSYQCPSDYFFDTDLLSECRPPNV
RGCRKCGKMGHLIRHCPSNRKEDDQKKQQSQANHGKQQNPNSQYKQTPPPQRQQRQLTPQKQQGQRNQPQLQERDIRDKEKRERERSQSESGNQGNSSAN
INRNQQAPKSQGPPFTHSQYNLASNPSLQPVTPAQLKNFQQTQMYHNSNYQYKQQYSMSYQQQQPRRNMGPSGQSSHGYNAPPPLLSGPMRPPMGNMPP
FVYNHPVPGQPNYHGPQMQPNAAFRGSPPQQGISYNSQGGVGPMPGGYGILAGGQGGVGQVGAGQMGQFSMNPVVQSIFASAHGQIMDGGSNQIQRR
KQ
**>Gfl-1**
MSLQNQDAAGRLKGVTVVKPVIYGNIARYFGKKREEDGHTHQWTVYVKPYKNEDMSAYVKKVNFKLHDSYPNPNRVVTKPPYEVTETGWGEFEIIIKIYFNDPM
ERPVPXYHLLKLFQSETDVMLGKKSLVVEYYDEMIFQDPTTAMNSMLANTTSLGIGPYKHETDFEEKRERTKNKISNAHSKIRFEMDQMNEKLKRAKDSIAKMKK
EISKLEEGATVDAEPSL
**>Mes-2**
MTTPEGACHTRSGAQRLLQASAATAPSSPSNNSLTESFPLGVVDWKKRVKAEYTRLCFLRKDKRADEVKNAFAANRIEIDRELAEEEEWLTRQPIQKLIVPETLPGIS
PKKCEVISGMGFPSQAGPLKLMMAVKSVPTMYSWAPVQQNFMVEDETVLHNIPYMGDELLDQDTSFIEELLKNYEGRVHGERISGIGDDLFLELVNSVAQQYR
MENGEEKIKREVKDGKEEFMDVSDIPLEVFEAICAAFPDQGGTVEELKDKYKELVESQKASLQDAPPECTPNIDGPKAQSVPREQTMHSFHTLFCRRCFKYDCFLH
PYHPTPSMLSRKMPEKKTKLEHCGADCYLHAIGSKDSRNGEEKDIKDNPKEVGRSSPVRPEVNGRKRKTNNGGSVASSENSEESNNADDVDIKFPPRLSPTVED
WTGAEESLLRVLADVFRSNYCSIAKLIGTKSCQQVYRFAVKEEAHIPDINEETAQTPPRKKKKKQRLWSMHCRKIQLKKDGSSKTVYNYQPCDHPGQRCDETCPCI
MAQNFCEKFCQCSSDCK
**>Ekl-4**
MAANMDVRDILELDTGQTQNDFVTKESLMTEGKKKKVRKLDAGIKRPEGMHRELWGLLHTDSRYQTDAAPIIPSDSKQGYKQMKARIGSSRVRPWKWMPFTN
AARKDGAVFYHWGRVADEGKDYPFARFNKTVDMPVYSDLEYQQHLHDDNWSRQETDHLFDLCKRFDVRFIVVHDRWDREKYKEHSVEDLKERYYNICNTLAK
VRAPGGSEPKVKAFDAEHERRRKFQLIKLFDRTQEQVEEEEYLIGELKKIELRKKEREKKQQDLQKLITAADSNADNRKAERKNTKKKLQAREPKEGGGLMTAEPG
GIKFSDFKASGTSLRSQRMKLPATIGQKKTKAIEQVLEELGLEFNPMPTEDIVNSFNTLRQDIVLLYELKLAMSTCEYELQTLKHRHDQLIANKNGGGSLSSYLMTTP
TKVPDVQSPTIEDAAGSNSQDGIPGSPSKRKLLVDTIDVVGTGSTPS
**>Mes-6**
MDGERIFSLSKRQLQMMHADISGDEMDDVSSTASTKEEESSRSETPTLRSRNGIRNKRQMKKCRLQFKCTNYLKEDHGQPLFGIQINLVMKETDPILFATVGHNR
VTIYECHDGGKIKLLQSYVDPSTDENFYCCAWSFDDATGQPILAVAGVRGIIRIISPTMGQCIKQFVGHGNAVNELKFKPKDNNLLLSVSKDHDLRVWNVKTGVCA
VIFGGVDGHRDEVLSADFNLSGTKIVSCGMDHSLKIWNLEKPDIRKALQESHNYNISKHDEPFPTEYCHFPDFSTRDIHRNYVDCVRWLGNLVLSKV
**>Rha-1**
MSGGYHGYYSQTNTHNNMHKGRQRYFDSDKEDLGPTSNREDRFFNDSNGPLGKSHGFTASERYGINDGKKFESDRGRKQNIFKRRETHRVPTFENAEDTTEQS
SSRPKRHPPGLRGREIGLWYAQRHIGQKEKERKNRPTVSMDKTRENSIRELLSNLKQDVSHQKNVTGSKNETASNILASSSSSKSWTSDLASFPSQFSQSSQHIVKD
EEYKEEVKEEPPDSWEDGDFDEMETEVEKPGSRSQEEPVDLDGYDDEDNQLPSVTGVVEYMKSHGDYPYAGEEPESDFMAINTDSLNPEYDMRMKLSMERLC
ENPQYVKMVEFRKKLPSYQMRETIVGTINSNQVLVVSGETGCGKTTQVPQFILDDFIQRGIGSQCRIICTQPRRISAISVSERVAAERCEKVDKSYTSSVGYQIRLETK
LPRSHGSILFCTTGIVLKFLEKDPCLKRATHVIIDEIHERDLQSDFLMIILKDLLPLRPDLKVILMSATLNAEMFSEYFNKCPMLSIPGFTFPVKEYLLEDVIEMTGYTSED
ISPRRRQWRKLKPGEREEQDNYAVWCRNLQGRYSQKTISVLQNFDFSKVNIDLMAHIVNYICTRLENGAILVFVPGWEDIKKLNEAIQNTPRCKSGSLKVIPLHSL
MPTVNQREVFERPPPGVRKIVIATNIAETSITIDDVVFVIDCGKIKVKGFQPEINLSSLETQWVSKANAKQRRGRAGRVQPGYCFHLYTEFQYSLAPDYLPPEMLRTR
LEELCLQIKLLKLGYIVPFISKAMQHPSKEAITHAVETLMELNALDANENLLPLGYHLARMPVEPHTGKMILFAAMFCCLDPILTVAASLSFKDAFTIPLGKEHLADQT
RIRLSGDSKSDHIMLINAFKGWEDSLRRGTSQRYCWENFLSDNTLKMLRDMKKQLAELLYDIGFIDSKDPKNLSANKNSDNQGLVKAVLCAGLYPNVAEITKVPTA
KSSKYRGAGGLGMLVKDNTKIQAHPKSVNMKQTFFESKWMVFYHKLKTTKVYIHDCTMVSPYPLLFFGGEIKILKEGGMDLVSVDDWVKFKASASTAQLVKDLR
QQLDKLLTKKITDPGPTCWDQTTSEGALMAAIADLITMEEKGYTVRGQSMGGSIGKKETGSSSKV
**>Ekl-6**
MAASINPLKSERIDNQLIFQSLTLLKAPAESKSTLSDGKEELLDNVLQENLVSLSKFLSHSPKLLLMLGINLDRNEKFINNDTELTTRQKFTQVCVELLQCLRKNLAEEIS
VYNDTLGTSQPAEKEIPNLPPGALSLQQESLVSTTLQFVLVLGMSPYLSPGIGIPVSHRLGPGQILLATSHDYTSELGHLQQIQFLLPVAKLLTDIFGVSSLQLLLVNSYLS
DLITVLIQIRYCAKMVQKNTESQKSLNLSSVANNNTLPKESAGPPGKTPAVVCETFEAHPTEEIYVYDSEIYRQKYLDPNQRLWTLQSVINYANAAIDTTVRSSSTPIV
IKAVLMLSGISKAGSTVKTPPWFRRTIACLLRDVLMLSSGVQNFICSFIGVQSSSTINSQDWQRCQAVAQIISKSPLTPNITQQFYTSVTKQLTELLQSSQVKEQPLVT
RAVGATVLELYTQHPDLVQSLCLSHLHAPLLSLTQHNAQYLQADILVSEDVMTSCLECLHRLYVLGQEPQSLILSTLVHIVGVLFEVFIFAMNSASNIKTLCCDLIKMYL
SHCDVKEAVTCLLTLTSNAGDPQVSPFPSLDSRAKLVWGGNGGVLAVADNTETILDTDSWTKPVDGAMAILTDVNSDVLLSQLFLNLLEKLTNMVTTETDHLDIQL
PGYLLSPKQRMQNISELRKKISYVSFLASLGEKFGEKVIQSGHHVLAFTKATLERCIKICLQTDDEPTKLFEWETVSMAMGLLTAVLGGAVQLTEQDKKLLDDLLPLLT
AVSESDATDPTVKEMAEDLKIAVATRGLVWSELKPTDKKKNFKPPAKSEKKEKPKESKCLIQVLAETTFDESTSNVDSDEISAPNDSNSSKIQQALQDLCDPLLPVRS
HGLISLARLVDDREPDIVGKSEMLLKVFLENMTHTDSYVYLAAVNGLAALSDLNPNLVVPRLAAEFKAQCLKDPGSKSPETTLKVGEALVKATKRLGELTPHYRHLLL
PAILIGCRHSDSLVRASSLSGLAEVCKLLRYAIGPVLFEIFTCCRDVISSDVDPEPRKAAVMTLTQLLQGLGKEALL

**>Zfp-1**

MSKKVKEMIGGCCVCSDERGWNENPLVYCDGAGCSVAVHQACYGIVLVPTGPWFCKKCESQERVARVKCELCPQRAGALKRTDTGGWCHVVCALYIPEAGFG
NVQTMEPILLSGVPNERFNKTCYICEEQKRDTKATAGACMQCNRNTCKQFFHVTCAQAQGLLCEVTGSYENVNYCGYCSHHYKKLKNQSNVKTIPAFKPIPSENG
TPDSSPEKIVQNKTEGEIKPPSQPTIEQVKSPQVDQSVSSTSDSVPSQLLAYSKPDLVSPGCSQVSDITSAVSPSGEDLKSPQSGSDLSQEPSSSGVITQQKSIITKLLQ
DGGATSSTENIEVDVGGVTEEVKLNVDVLPVTSEETSQAMSKRQRQVNSQKPEKSAVKKTKSTTISSTVPHRKDGSKKAMKKRRDSGGSSTKGATTSSNTATTLPG
HHNYCGSIFGGSPYFQSLSAPGRVSLSNFASHDSSALNGTPPLVQPPKQFPSIHTSKQGQDHSQDFPLSMEQLLEQQWDHGAQFLMDQGQHFDIASLLNCLHRL
KGENQSLEDQVRDLTSRRNHLMAVNARLSLPLSVYDGCLSNSQSSSMGSLEDISSPDDVQPKSGLTTVQIPCNSDLIVEDILSPVDTEHPSLSNHKLTTSCSPGYYPD
QICPPQPLMYSVVNPPPPTLSVISTTLPSRHTSTPAHASMSSHHTSALPHSAMPALEPSHIPNLSGLSLNNSNVTTLQDTT

**>Mut-2**

MYPRGSGFPFSNLSGFASSVRFFQQPVHVNLPQMSTNSNSRMQQRWPDFIPLENLTRHNFQQSSSGPNFQQNLTQPQCFPVNSVQHTRLLNTSAALNGSSSA
QKRRMNGTESSPKKKQKTSNSISPLTLPVEPGNFLIPVARGTEKITHDIWEFFLASQMNEEDLKKKLKLRKCILSVMTGAFPNCRLFIVGSSMTGFATKTSDVDMCL
MISENDIDQRRDATVILSSIAKALRACSFVRTPQVIKAKVPIFKFFDAVSGVECDLNINNTVGVRNTHLLRYYAYMDWRVRPLMLFIKKWARFHDINDASKKTISSYS
LTLMLIHYLQAGVIPAVLPCLQTLKPHLFSIQADVCRLSLNFEPDNVKFASKNVATLGELFLGFLNYYSNIFCYESQVISVRLGRTIGRHEIYDRDNTMQWKHLNIEEPF
DRTNTARSVYDLYVFQRILRVFKVSFNTLVNTRDISQILSRPF

**>Ekl-5**

MYPRGSGFPFSNLSGFASSVRFFQQPVHVNLPQMSTNSNSRMQQRWPDFIPLENLTRHNFQQSSSGPNFQQNLTQPQCFPVNSVQHTRLLNTSAALNGSSSA
QKRRMNGTESSPKKKQKTSNSISPLTLPVEPGNFLIPVARGTEKITHDIWEFFLASQMNEEDLKKKLKLRKCILSVMTGAFPNCRLFIVGSSMTGFATKTSDVDMCL
MISENDIDQRRDATVILSSIAKALRACSFVRTPQVIKAKVPIFKFFDAVSGVECDLNINNTVGVRNTHLLRYYAYMDWRVRPLMLFIKKWARFHDINDASKKTISSYS
LTLMLIHYLQAGVIPAVLPCLQTLKPHLFSIQADVCRLSLNFEPDNVKFASKNVATLGELFLGFLNYYSNIFCYESQVISVRLGRTIGRHEIYDRDNTMQWKHLNIEEPF
DRTNTARSVYDLYVFQRILRVFKVSFNTLVNTRDISQILSRPF

**>Ago-2**

MTELKGRVLPAPKLQYGGRTKAQAVPNQGVWDMRGKQFYQGIEIRVWAIACFAPQRTVREDALRNFTQQLQRISNDAGMPIMGQPCFCKYATGPDQVEPMF
RYLKNTYQGLQLIVVVLPGKTPVYAEVKRVGDICFGLATQCVQAKNVNKTTPQTLSNLCLKINVKLGGVNSILLPSIRPHVFREPIIFLGADVTHPPAGDTLKPSIAAV
VGSMDAHPSRYSATVRVQEHRQEIIRDLATMVKELLIQFYRSTRFKPTRIIFYRDGVSEGQFSTVLGHELRAVREACMKLEVDYQPGITFIVVQKRHHTRLFCADRR
DQTGRSGNIPAGTTVDQGITHPTEFDFYLCSHAGIQGTSRPSHYHVLWDDNRFNADELQILTYQLCHTYVRCTRSVSIPAPAYYAHLVAFRARYHLVEKEHDSGEGS
RHSDNSEDRNPLYLARAVTVHPDTCRVMYFA

**>Dcr-2**

MAHSNSTGPRKGSRYHRLDDIPTDTFTPRTYQIQLLDSALKRNTLLCLGCSESKLFLALMLAKELASPTRLQRSQGGKRTFYLTDTDDDVEMLYQTLPHHTDLQVEF
CYPHSLLEDDTQDEISGTQMSLQKWCLLVDSSSILVCSGHLFLTALCKNYIDLKEVNLLVFDNCHRAVEVENHPYKEIVNGIKSLPQKSDQPHILGVTASIAGTDCSDP
LKLQQTISEMEEAMFSYAETSMLILSERFGCRPKESVVQCLDLELDDEQEVLKDTLEEMLEQNYYFFSDCRIAVDETVDNRDPRETPMQVLSQCLNILNVLGSWCT
ASISEYFVMQLEKIVKCEKNEIHKKFLRCVITTLRVLIKEFEKQFHPDYHVNELLEYTTPKVRELVKVLRKYKPEVDFIIVSNGDSFNDDEEFSDLSDEDDDSIMSDSDG
EDNSGEPFNGSSKPMHIAVKRTADGGHERIMTFSEEEEKNLCGIVFVENRYIAFGLNKVIEEVCSWDENLCFVKSCHVTGQGLRSANGKPRVNRTYKRQEEALRKF
RTQECNLVIATLELEEGIDIPKCNLVIRFDPPKDFRAYALSKGRARARDAAYVILLDDENFEGFNTSLKVFKGIEQVLLGENRSLQAIRSNIPGEDADNEYDSKDDDDSD
ASSDERDNLPPYCTMPHSLSSPHVTLQGAIALINRYCAKLPSDAFTHLTPQCRIEKLDGYAPSYIANLRLPINSPVKEELKGPAMRTKKLAKMAVALKMCEILHKKGEL
DENLVPVGKEMFLYEEEEEWVNEEDLAGQARPGTTKRKQYYIKKTTDALLQCRPKPGQPCFLYLINLKLTGPITDEQNTRGRVIYAPEDTVQTLGILLSKKIPLMPYFP
VYTRSGEVTASIDLQQTDLICSEDEINRLLNFHHFVFSGVLHLEKDPMEFDCVNADCGYIIVPLNGKGCEELTIEWSFVDKVIDFIKVEKRAGFSKKSGDESFQFCLDD
FQDAVVMPSYRNIDQPQHFYVAEIRHDLNPLSPFPSPELYKTFQDYYTTKYGLMITNTEQPLLDVDHTSARLNLLTPRYMNQKGVALPTSSAETKRARRENLQQKQ
ILVPELCDVHVFPASLWRKTVCLPAILYRANYLLLADELRRKISRETGIGVEELPPGFRFSKLDFGFDTSPEKLMDAEVNCNKKISSNIKEESAERSPDPVLCCMEKDSS
NNRMIKESLVNDTDKAKDESSTSSDSDSGIASSSSDNNIDITPSHKILIKSSEHLEHGSVNPSKKFQELCGHDNNVNLTQSWCLDHNNILYNVTPLGSSFPGTNQCNI
ISNTEAVELPSTSSAVPQQDSNGSTTTSTVINFSCTSAPAKVSPDNATVGATISIMAPPSIGVVSSCEPDKPFDSTDSEPLAAETLPRQAIHGDKTSVALSLPCSTTSSS
TLLTSSAPLESPSAATSTWSMPITSTTFGEKGHSSDYPPLFVSASEVTNASTINKACYLSPRVLTNFNSDAASNRANCQADLFITTDINGCELNKGLNDASPALLATET
SPAIDHSHPNHCSCINHPCVHHSDAQIISADQKHIHAIYNNVSHNNAESSSNSAKITSGKAKSHINGHVKQDIKYSVAQRTINEFCNQDKTESKGHAKAKCQLSSD
GPAIIMNSLSQELSDLDIDLIVPQADTPDTFHSFEDSTRSGANTRKDNQHKKTTVRLSAPANSEWPKGQQTTTVDWPDVQQTGNSERLNCQQFTLHEQRKPVH
KGFDKISVDKGFDKISVDKDETGSANVTTDNYNATAAAEDIHVIKFSFNTDEPGVKCKEAKEPSPPDNKKESSPTDSKKECNPPDSKKELPNENTENRLTFDLTRVKS
PLETQQRNTELEIGVWKSSEKRIYDKHQAADSHREKITSLDHSSYNEMTETNHSLCQPTFSCVTTGSLTSISLDEDRDPQTFVGPSPCLIMQALTMSNANDFFSLER
LETIGDSFLKYAITVYLYCSYPGIHEGKLSYLRSKQVSNYNLYRLGRRKGLAECMVSTKFEPYENWLPPGYVINDERRRGPVPKVVIVTPGSKINNSLRNFYLDETADA
AKRDEAILFNKELEWIQQSQEADQQEEVQEPNNDMTPYSLQCHHGLPDKSVADCVEALIGCYLTTCGRKAALIFMSWLGLRVLPKKKKLILDVSKIEEHEKSSVGK
LSEQEFDELRCPPSPMFSNTPANVAKLNHLLQGFDSLEEKICYKFRDRSYLLQAFTHASYHYNTITDCYQRVEFLGDAILDYVITRHLYEDSQKYSPGILTDLRSALVNN
NIFAALAVKWGFHKYFKAISPSLFQVIDKFVKRQKEKKEDDIDIDEEFRELSLEENEPEENTEEDDEEEEEVEMEIPKALGDIFESVAGAIYLDSGMSLDTVWRVYYR
MMKPHIDKYLKSIPKSPVRELLETEPETAKFERPERTMNGKVRVTVNVVGKGVFSGVGRNYRIAKSAAAKKALRSIRTLNQNLMN

**>CG5382**

MGLCKCPKRKVTNLFCFEHRVNVCEHCLVANHAKCIVKSYLQWLQDSDYSPLCTLCNHNLSDEEYGECVRLTCYDVFHWGCLNQFAQRLPAQTAPAGYTCPTCSA
CIFPQSNLVSPVADAVRILLQQVNWARAGLGLPLVS

**>CG5434**

MAQPGAAAPNLYAELNRCLQKQEHEKAIRIANKILQQNPEEFKAFNCKIVSLLLLDRFDEALTALNKDKTHTGTLKFEKAYCEYRLNRTNEALNTLRSITSHNTRSKEL
LCQVLYRIEEYEECYSLYRDVIKNSQDDFDSERETNLAAVVASLQMWNQKDMGCPDIDESSYEICYNNSCYYIGKGDLKTALEKLDKAEELLKSDGDLTEEELDEELAI
IRVQRGYIFQSLGKNDQASQLYNLVLKTKPSDAGLVAVVSNNVVTINKDQNIFDSKRKIKAATGDNLKHKTVSAQRQHIDINQCLVHMYSNQSEQCHQLAKKLAD
QYPDLDAPLMIRAAQYIRDKKNDEAIALLKEYIKTRPEKAFKAHMIIAQLYLGLGSVYQACDALKALGQESFKPGIVSTLVTLYLSQEDKDAASKALIDSVNWYKKND
PKSPALMTLTRANADFQLKNGKADEAARMLEDLRKANPGDALVLAQLISSYAQFDPAKAQQISKDLPPVVKIISGLDIEALEASFSTLGPKYMKKQQPLKGEASPGA
SGDTKSGDTLMQKKKQRKKKKGKLPKKCKPGSDLDPERWLPRKERSYYRGKRKDKKKEIGKGTQGTSAATQAITDSLDASKQPASGEPASPRPSASASKASPHSTP
TPMPLPQGPRQQKPVQANKKKKKAGKGGKW

**>CG4572**
MEFSFAISIGLLGLLGFLEIVSASVPLFLTPYLEKGQIKEARGHSRVTEENTTGVTIPESYSGFITVDKHLKNHLFFWFFPSALNASAPLLIWLNGGPGVSSMLGLFWEN
GPLELRSDRKDFQKREHSWADPFAMLYIDNPVDVGYSFTESGDAGHRSTQKGITKDLYSFIEQFYKMFPEYKKRELYIGGQSYAGKYVPSIAYYIDEQLQKNRTDIPLT
GVYIGGPFFDPPVQTLSYYEYIYSMGVISHAQKVEYRQDTLKLIQQFHAGELKNVELKEVLDKILRNLGLATLDNYVTGNPADYMLVNECMNTPRIRHAVHAGNRT
FHIINRAVNAKFGGDLFVSLKSEMAELMNKYKVLIYTGDYDVIVSSVMVESALLTTPWRHQAEYNATRRSMWWSAEGQLRGFYSKTDQFCRVVVKGAGHQTPH
DQPQASLEMMREFIQHGCVAIKATGE
**>Hsc70-3**
MERLTPLFLFVLICTTCFVHAKEKDDDDETDKKKSDVGTVIGIDLGTTYSCVGVFKNGRVDIIANDQGNRITPSYVAFTPDGERLIGDAAKNQLTANPENTIFDVKRF
IGRSWDDKSVQHDIKFYPFKVINRNNKPYVVVDTVDGEKTFAPEEISAMVLGKMKDIAEEYLGKKITNAVVTVPAYFNDAQRQATKDAGVIAGLNVMRIINEPTAA
AIAYGLDKRDGEKNILVFDLGGGTFDVSLLTIDNGVFEVVSTNGDTHLGGEDFDQRVMDHFIKLYKKKKGKDIRKDNRAVQKLRREVEKAKRALSSAHQVRLEIESF
FEGEDFSESLTRAKFEELNMDLFRSTMKPVKNVLEDADLKTDDIDEIVLVGGSTRIPKVQQLVKEFFNGKEPSRGINPDEAVAYGAAVQAGVLSGEEDTGDLVLLDV
NPLTMGIETVGGVMTKLIPRNTVIPTKKSQIFSTAADNQPTVTIQVYEGERSMTKDNHLLGKFDLTGIPPAPRGVPQIEVTFEIDVNGILKVTAEDKGTGSKNQIVIQ
NDQNRLSPEDIERMINDAEKYADDDKKIKERVDAKNELESYAYSLKNQIGDKEKLGAKLSEGDKEKITEAVDEAIKWLESNADSDAEEFKEKKKELEGIVQPIMTKLY
EQTGGSPPPTEEDSDEKDEL
**>Hsc70-4**
MSKAPAVGIDLGTTYSCVGVFQHGKVEIIANDQGNRTTPSYVAFTDNERLIGDAAKNQVAMNPENTVFDAKRLIGRRFDDPTVISDMKHWPFTVINEGGKPKIR
VEYKGEKKTFFPEEISSMVLTKMKETAEAYLGKTVTDAVVTVPAYFNDSQRQATKDAGTISGLNVLRIINEPTAAAIAYGLDKKVGGERNVLIFDLGGGTFDVSILTIED
GIFEVKSTAGDTHLGGEDFDNRMVNHFIQEFKRKHKKDISENKRAVRRLRTACERAKRTLSSSTQANIEIDSLFEGIDFYTSITRARFEELNADLFRGTLEPVEKSLRD
AKLDKAQVHEIVLVGGSTRIPKIQKLLQDFFNGKELNKSINPDEAVAYGAAVQAAILHGDKSEEVQDLLLLDVAPLSLGIETAGGVMTALIKRNTTIPTKQTQTFTTYS
DNQPGVLIQVYEGERAMTKDNNLLGKFELTGIPPAPRGVPQIEVTFDIDANGILNVSAVDKSTGKENKITITNDKGRLSKEEIERMVNDAEKYKNEDEKQKTRISAK
NALESYSFHMKSTVEDEKLKDKISADDKKVIIDKCNEIIHWLDANQLADEEEFQHKQKEIEGVCNPIITKLYQGMGGAGGMPDFTGGAPGAGSGASAGTGGGSG
PTIEEVD
**>Cog-2**
MTVDVANKGFTLPSGPSSLCFDKEEFMKKDFEVDKFIMECRKKVPLETLRDDLDLYLKVIRSAMIELINKDYADFVNLSTNLVGMDKAITNLTTPLGQLKEEVLSVQT
AMDEAIKAVQDKIRQQQEIQRKKATLQRLMNITHSVEKIEQLLGIKHGLDASETASELTGPLIERVATEFNKLQFYVTRSKGLPLVDQVKPRISAITTTLQCNLEGSFLS
GLETGNTAMLRQCLRTYALIDKTQDAEKLFREHIVQPYMERIINDDFLRHHNKDISVMFDHVLQFFSDKCSMLTGITTGTSGGEIVRGYDFVVHAVFPEVVENLEQ
QIPTIFAPGNPDVFHKKYTATMKFLDQFEQYCGTQASVRRLRQHTSYTTLMNKWNSQLQVYFQIRFQDIAGSFEASIFSGFIKSSEGTGFHLSSSAALWTSLVRCW
QDDIFMPAYTHRFLKLDLQLLSRYKTWLDDVYQEEMSRKNESNPESKPRSATPDMQSMPSLSDDVMTRSMSPKPTYGQTDQSRPVLTNGQIVALMADMDLLIA
KIPILYEEIIKPKLSTLGLDDSSAVKDALIGGLQASCECRPRFWSIVTEDVVKRCSTFLAQVNDIPRLYRRTNKEVPSKPSTYLSSVIKPLSQFCAEHSQALNQQQKSEFL
THVFGTLSQQFSEVTAEVLTSTRKMEESLRRLKKARGGEKEKEKAGVGDSDKIRTQIIIDIQSFHSQMQEFGLVPSSIEGFTKLSTLAQEAQADITSSN
**>Cog-3**
MTRAQIITSMKKEDIGSKVTLKQQPFATPEKVHDVVAETYRNIKTKLPIVQRSMSLYLANKDTEAILFRPIKVNVQQSFQKFQELLTQEYSEEDILIIACPSAEQ
**>Trs-20**
MSGNYYFVIVGHHDNPVYEMEFSPPNKPSEKKDDHRHLNQFIAHAALDLVDEQMWSTNNMYLKIVDKFNEWFVSAFVTAGRMRFLMLHDVKNEDGIKNFFT
EIYETYVKFAMNPFYEINTTIKSASFDKKAQFFGKKYLSG
**> Pi3K5PF**
MIPIGGTTISVFGKRGTLRKGMHDLKVWPDRKPDPSIESATPGKSKDSSDRMSQLSKLAKKHRDGHMTKIDWLDRLTFREIELVNEQEKRDSECMYLMVEFPRV
HFENLNFSVVYFEKEGERPYEFRTQAEVVTIHDPELLMENIVESKHHKLARSLRSGPSDKDLKPDASTRDLLNTIVGYPPTKTLSSEEQDLVWKFRFYLSSQKEALAKF
LKSVNWKLAQESTAALEMLLLWSPMNVEDALELLSPAFSHPRVRKYAIMRLRQANDEDLLLYLLQLVQALKYENFEEIKNRSDRDFVPLRPEFDTGRRDSDLTSLM
ALEPVTGLDQHPLSQSNITSSTSSLEEETKVRSDSATTEEEILDAMNSGPSSGKNGMSLSALDDEMDLATFLISRACKNPVIANYFYWYLVIECEDNKDIQKEEVLN
MYKTVLKRFSLALSKGPPQSKKQRAILSRQQTLLKKLCTLVESLKNVGSQRKKIEHLQAMLHKPEFSEMLSSEPIPLPLDPSVLVKSIVPEKANVFASKLSPCLLTFETVS
GKEYKVIFKYGDDLRQDQLILQMFQLMDKLLRHENLDLQMTPYKVLATSKEHGFVQYIESKALSELSKEKAIQNYLRQHNPSETGPFGIAPEVMDNYVKSCAGYCV
VTYLLQVGDRHLDNLLLTKNGKMFHIDFGFILGRDPKIMAPPMKLNRQMVEAMGGYDSEHFQRFKVFTYTAFLALRRSANLFLNLFSLMVDTNIPDIALEPDKTV
KKVQEKFMLHLNDEQAVQHIQGLIDDSVNAFMPTLMEYGHKVAQALR
**>Rab-7**
MASRKKVLLKVIILGDSGVGKTSLMNQYVAKKFSNQYKATIGADFLTKEVMVDDRLVTMQIWDTAGQERFQSLGVAFYRGADGCVLVFDVTMPNTFRSLDSWR
DEFLIQASPRDPENFPFVVIGNKIDLENRAVSARRAQGWCHSKGDIPYFETSAKEAINVEQAFQAVAKNAMAQESDVELCSDFPDPIKLKDDQSKPKEGCAC
**>Rab-10**
MAKKSHRKTSWKKKSHWKTSWQKKVTGKLHGKKKSQENFMAKKSHRKTSWQKKSHRKTSWKKKVTGKLHGKKKSQENFMAKKSHRKTSWQKKSHRKTSW
QKKVTGKLHGKKKSQENFMAKKSHRKTSWQKKITGKLNGKKKSQENFMAKNSHRKTSWKKIVTGKLHGK
**>Sid-3/Ack**
MHTLRHTNLIHLYGVVLSTPLMMVTELAPLGSLIERLRQVDQAPLISTLCEYAIQIAVGMTFLESKRFIHRDLACRNVLLKTNELIKIGDFGLMRALPSQTDHYVMSE
QKKVPFAWCAPESLKSRQFSHASDAWMFGVTLWEMFTYGQEPWLGLNGSQILHKIDVENERLPKPSDCPSDIYQLMMHCWAHKPQDRPSFAALKDFLSEVLP
ENVKATRSFSEEKRLKIEEGDLITVIDGV
**>Stau-1/2**
MSHNHMLSNQQQQHSLMNSGATASGGQQRTMIAQRGSQVLAQPKSAYQLQPASVSQLSHIQQQQLVNNFHSQKQMSPDISSTHQYNSAIGGLPSQNGNYQ
SLTSTSLHSNITLEGNALSVQHQHHQYPQTQQPQPPIQPPVPQHHYQSASVSQPQQPSQHHLHQSMQQQHHHLLQHQQQIQIQQQPIPQHQSSSYQQHQ
QAQLQIQQHLQQRSLHQQQRFQKQQQVQPKQILQHPTSASKNVSSQIQQPSQPSTLQLQPQQQSKPLIINGQSGITDSKKESSDKVEVKLDDNEESEESKNNAT
EMLGLANTKEKTPMCLINELARFNKMSHQYTLVDEQGPAHKKTFYVKLKLGDEEYSASGESIKKAQHAAAAIALVETKHPHPPPKPARYSGSNSVDSGDENITPTV
ELNALAMKRGEAAIYKAIEPQQPPYYHQPGMDYRGIYSQRYHQYMRASRDPRYRGSGVLWPLRYHYPRMNRPFYVSLRVGHREFIGDGPTRQAARHNAAQKA
LRILKNLPVQHEAKKDQEQAEEESNDSLKSEISLVHEIALRRNMPVNFEVIREAGPPHMKNFITRCLVGEKVTEGEGNSKKTSKKKAAELMLEELRKLPPVPTPLYPR
PKSKIQLNKKKNRNLIKSELQQQKADPNYGVGINPISRLIQIMQAQKKKEPVYTLVAERGLPRRREFIMQVEVEDKNCTGTGPNKKLAKRSAAEAMLQILGYTKPSP

QPAKSAFKHPPSGEGNQSNGDKKVTFVEGDPPAESTEKPSGPTQSIQQRVPGLLHLPTSNSKLVMNSAASSSSSLTQKESLVNLASILKPNLRPEVQLRELCKALGQ
ELEIDDFTKKRQTGTEHITRITVGKEPSQNFHGSSNTLESSRDMAALDALKVLLAKVKDIHPGGDGPQVKKELLTRSSSGMKKDISK

**>Gmer**

MVGGLFRNLKYNLDFFRVNSQINDNVLSISHQYNVQKVVSCLSTCIFPDKTTYPIDETMVHNGAPHDSNFGYSFAKRMIDVQNKGYHLQHGRKFTSVIPTNVYGP
HDNFNLEDGHVLPGLIRKVYEAKASGTPFVIWGTGSPRRQFIYSLDLGRLFLWVLREYDEVSPIILSVGEEDEVSIKEAADLVVEAMGFTGEVIQDTTKSDGQFKKTA
SNAKLRRYLPDFKFTPIKQAIKETCDWYEANINTARH

**>Clint-1**

MLSTSIWKLRELTDKVTNVVMNYSEVETKVREATNDDAWGPHGSLMKEVAQYTFTYEHFPEVMGMLWKRMLHDNKKNWRRVYKALILLNYLIKNGSERVVTS
TREHIYDLRGLENYSFTDEQGKDQGLNVRHKVKELLDFIQDDDRLREERKKAKKTKDKYVGVSSEVMGLGGGAAYSDRYDEEPRSYRERGPNQMEEIEDWDSGK
KSVVSGAIDKAKDLWNRAQGRQAPDEIDPERYEEEGWDKSKEKEKDSRERKAERDRYDFKDDDEEYTSVERTHTTKTEKITTNRRTRSIGKKLDLGAASGLGKDG
DSQSQTSASHDSAPSLFDLSDPAGGQEGFADFSTFQSAAVTNDEFNPRASGSNDFGDFSKVGTNTSSTQGGFADFAQFNSATSPPSISPLVASVPAGPATTQSGSS
AVNDLFDVFSAPSVAPGLTPMSAGGMMMPGMVMTNMNVPQGMNMVPGMMPMSGMTPVMQQPMMMQQSSMTQPMLMQQSGVNYPGMMVLG
SGVNSNMQVRPSGLASQNTSVKQGTSGEKKGNTWNDTPVNISLDGLTPGSKLHKQSSQGPSLNQLTGGQPMPQGMGLITPGMGGINQGMANMSLQSPTN
PLMPSSMMVAGNPRPVMSNMGMMGMGQTPMMTGMPQGTMMAGIPQGTMMNGMPQGTMMGGIPQNPMGIQMMSTTNMAGQASFQKRTDTAF
SAFGNVK

**>Haf-6**

MALLLSCQRLSHIYKIPSLSPKLCRGFLKSSRGCENIKSPFTDSSLTHLIESHVKQNGTLSRTYHTLSKLKNRKFIDEVLQGPSWRSRQIFRSCRRNETTTPKKEANKLTE
AGKAPRRSLPKPVDVRRLLGLAKPERWKLAGAICCLVISSAVTLSVPFFIGKIIDLIYTSEKENMLSTLRTFCASLIVIFIIGGLANFGRVYILQSASQRIVKQLREKLFTRI
MHQEVGFFDKTKTGELINRLAVDTQVVAQSMSQNISDGFRAILQASTGIGMMSFISAKLTLISMGIVPPVAIGSIIYGRYLRKITKQVQNSLADATQVAEEKIATIRTV
RAFAHEPKECKAYSDGVEKVLQLSYKESLARAIFWGFTGFSGNVIIISVFYFGGHMMAESNISMGDLTAFLLYAAYVGASLNGITSFYTELNRGVAASERIWNLIDRK
PLIELDSMSKPAITLNAIEGEIGMKNMAFSYPSRPDAKIFSDLSLSIPAGSITAIVGPSGSGKSTLGALLLRFYDPDAGIITLDGHDIRDFEPRWLRSKIGIVSQEPILFSTT
IKENISYGSEIPETVQMDEIIYAAKKANAYNFIQEFPDGFNTLVGERGIMLSGGQRQRVAIARAILKNPKILILDEATSALDAESEYQVQDALEGLMVGRTVLVIAHRLS
TIKSADNIVVLDSGKVVESGPYDHLIGLKDGLFKKLVERQTLIK

**>SR-CI**

MGCDFEGGLCGWSHDTSDDFDWTRNRGETPTADTGPLVDHTTNSVHGHYLFAESSSPQEAGKKTRLISPIYNPDSVVGLCLEFYYHMFGTDGDGNVGQLDVYI
KPKDISPDKLTFKHQRFHKEGNQGDQWMKADFEIPEEKQPFQIVFQVTILNSWTSDIAIDDIRLHNCKEGDQTS

**>Pgp-11**

MMTVFYTSFVAAIYMGLLIPTAKDIFTAKAALVKLSSIIHKKSTIDSSSTEGLTPQTMSGTIEFKNVTFAYPSRLEVKVLNDLNLTLSKGQTVALVGASGCGKSTVVQLIL
RFYDPIQGQLCIDGHNIKDLNIKWLRRHIGFVSQEPVLFATTITENIRFGREDATMEEIIEASKKANAYDFIMALPLQFETYVGEQGAQLSGGQKQRLAIARALVKDP
KILLLDEATSALDTESESIVQDALEKASQGRTTIIIAHRLSTIRNADVILAFSGGCVAEQGTHDELMARGGIYTQLVALQSKKNEHFSDSAIRSPNEHSKKAENFKWD
MANGSHTPNLSNLKQQNNQKILKYKKAQHAVLRLLKLNAPEWFFITCGLIGGSLSGFMQCIWAFISGTILQALSSVDLQFQERAMNNSSLLSVGLAFFVIITNLAQ
DAMLAVSGESLTTRLRDRLFRATVRQKMSWYDDPQNSVGILTTRLATEASQVQEAFTTTIGLLINSATILVLGLAFTFYFSWKLAFIFVCAAPIVIVSARVQQYLTLKIK
GTSRKSIEIATAGALESISNIRSVASLTIEDKFYEQYASSLKKASHSMNGYLLSAGICYGFYSSIPYFVNCASYIIGKILIESQELEFYNVIRTITCMSMSAISLFRVLILLPDTT
AANSSAEKIFTLIDEAPTIEECRGGTLTIDEDEFKSSVRFSGVHFHYPMRPEQSVLNGLDLQVSPGKTLALVGESGCGKSTTMQLLERFYDCQQGNILLDDKYDIKDL
NIPWLRLQIGLVSQEPVLFDRSVSENIAYGDNSRKVNMEEIIQAARDANIHNFIEALPCGYETNVGSKGTQLSGGQKQRIAIARALVRNPKILLLDEATSALDTESEKV
VQEALDKARKGRTCIVIAHRLSTITSADTIAVLKKGVIHEIGTHRKLMAKRGLYYQLQMTQTG

**>Haf-2**

MALLLSCQRLSHIYKIPSLSPKLCRGFLKSSRGCENIKSPFTDSSLTHLIESHVKQNGTLSRTYHTLSKLKNRKFIDEVLQGPSWRSRQIFRSCRRNETTTPKKEANKLTE
AGKAPRRSLPKPVDVRRLLGLAKPERWKLAGAICCLVISSAVTLSVPFFIGKIIDLIYTSEKENMLSTLRTFCASLIVIFIIGGLANFGRVYILQSASQRIVKQLREKLFTRI
MHQEVGFFDKTKTGELINRLAVDTQVVAQSMSQNISDGFRAILQASTGIGMMSFISAKLTLISMGIVPPVAIGSIIYGRYLRKITKQVQNSLADATQVAEEKIATIRTV
RAFAHEPKECKAYSDGVEKVLQLSYKESLARAIFWGFTGFSGNVIIISVFYFGGHMMAESNISMGDLTAFLLYAAYVGASLNGITSFYTELNRGVAASERIWNLIDRK
PLIELDSMSKPAITLNAIEGEIGMKNMAFSYPSRPDAKIFSDLSLSIPAGSITAIVGPSGSGKSTLGALLLRFYDPDAGIITLDGHDIRDFEPRWLRSKIGIVSQEPILFSTT
IKENISYGSEIPETVQMDEIIYAAKKANAYNFIQEFPDGFNTLVGERGIMLSGGQRQRVAIARAILKNPKILILDEATSALDAESEYQVQDALEGLMVGRTVLVIAHRLS
TIKSADNIVVLDSGKVVESGPYDHLIGLKDGLFKKLVERQTLIK

**>Pmp-1**

MVSPNNMAAHSKLLKFTPRNSALITAAGIAAYFIYKRANKKATKRKSNAPEEEVKVVIAQHGQKKEKAVVNSVFFARLGKILKVLIPGVFTKETFYLAIVAVSLVARTY
ADVWMIQNGTSIESAIIGRDAQQFKVLLFKFIYAMPVISVVNNLLKYGLYMLKLRFRIRLSKYLYDKYLKGFTFYKMSNLDNRISNADQLLTQDTEKFCDCVAELYSNL
SKPILDVILYSVKLAGAIGVMGPTYMLMYLAFSGAVLTRLRRPVGRMTVAEQKLEGEYRYVNSRLITNSEEIAFYQGNSREKFIITGTFHKLVDHLRNFIHFRVQMGFI
DNIIAKYIATVVGYLVVSRPFLNLAHPRHKDSSHSELLEDYFKSGRMLVKMAEAIGRIVLAGRELTRLAGFTARVTDLMKVLDDLNTGRYERTMVSTDNGRGSSNNL
KALTLKPGNGKIIIKDHIIKFDKVPLVTPNGDVLIEELSFEVSSGKNVLVCGPNGCGKSSLFRILGELWPLFGGTLTKPEKGKLFYVPQRPYMAVGTLRDQVIYPDNHA
EQLKKGVRDEELAEILDKVQLNHILEREGGWEAVQDWMDVLSGGEKQRIAMARLFYHKPQFAILDECTSAVSVDVEGYMYQHSREVGITLFTVSHRKSLWKHHE
YYLKMDGRGYYEFKPITDDTTEFGS

**>Ran**

MAQMDDIPTFKLVLVGDGGVGKTTFVKRHKTGEFEKKYVATLGVEVHPLVFHTSRGPIKFNVWDTAGQEKFGGLRDGYYIQGQCAIIMFDVTSRVTYKNVPNW
HRDLVRVCENIPIVLCGNKVDIKDRKVKAKTIVFHRKKNLQYYDISAKSNYNFEKPFLWLARKLVGDPNLEFVEMPALIPPEVQMDAALARKYEDELRVAQETALPD
DEEDL

**>Tbc-3**

MTTLAHLFQEKEVQEIFERILYIWAIRHPASGYVQGINDLVTPFFVVFLSEYIENDIEVENCDLTELSQETRDLIEADSFWCMSKLLDGIQDNYTFAQPGIQMKVNAL
RELIKRIDVPLFNHLERQGVEFLQFSFRWMNNLLMREIPLRCTIRLWDTYQSEINGFADFHLYVCAAFLKRFSEESLREHDFQGILMLLQNLPTLHWRDKEISELLAE
AYKLKYMFADAPRHLLLKS

**>Mtr-4**
MSVLRDAGDAAKGDQRGRRGGSKANESDCYKIVKLIMERNFAPVIVFSFSRKDCEGYGMQMSKLDFNTDEEKTLVEEVFNNAVDALSDEDKKLPQVEHVLPLLR
KGIGIHHSGLLPLLKETIEILFSEGLIKALFATETFAMGLNMPARTVLFTGCRKFDGKDFRTVTSGEYIQMSGRAGRRGLDERGIVIMMVDEKITPDVGKAILKGDPD
PLNSAFHLTYNMVLNLLRVEEINPEYMLERSFFQFQNYAAIPDLLERLQVKEQAYKALAVDDEDSVSAYYRIRQQLDTLGRELRHFILKPTYLLPFLQPGRVVKVKNE
DLDFGWGIVLNFHKKMDQSAATAETGTLYVVEILLNLTKDSLKTRSSAEIRPCPKGEKGEMQVVPILTRLIDSVSAIRLYIPNDLRSTDSRFLVLKSLQEVEKRFPDGLPL
LDPIEDMGIKEKSLLDTIKKIEAFEHRLYSHPLHKDSRLAELYEQFEKKSKLANDIKALKMELKKKKSLLQMDELKCRKRVLRRMGYCTASDVIELKGRVACEISSGDEL
LLTELLFNGVFNELTPQQACALISCFVFQEKASETPKLGEELAGPLRIMQDTARRIARISKEAKLELDEEVYVDSFRPHLMDVVNAWCNGSTFANICKMTDVFEGSII
RAMRRLEETLRQMVQAAKAIGNTELENKFSEGIRQIKRDIVFAASLYL
**>Tsn-1**
MTAPAAPAVIQRAVVKQVLSGDTVVLRGQPKGGPPPERTVCLSNITAPKLARRANPNVDTSVETKDEPWAWEAREYLRKKLVGKEVVFTVEYKVPGTGREYGIIYL
GKDTSGENITESLIAEGLVDVRKTGLKSDDPNQQRLAQLEDAARAAGKGKHAEGEASKHVRDIKWTIDNPRHFVDSHHNKPIDAVIEHVRDGCTVRAFLLPSFEYV
TVMLAGIKCPMNKQDTDGKQVPEPFMEEAKYFTESRLLQKDVKIILEGVSNQNLLGTVLHPNGNITEFLLKEGFARCVDWSMGVVSQGAEKLRAAEKSAKDKKL
RIWKDYKPSESSINIKDKSFSGKVVEVVNGDGLVIKLENSFKKVFLASIRPPRPADATGDGPVKETTSKRSRPLYDIPYMFEAREFLRKKLIGKKVNVEVDYIQPPNQG
YPEKTCCTVTIGGINVAEALISKGLATVIRYRQDDDQRSSHYDELLAAEARAVKKGAGLHSKKEAPIHRVADVSGDVAKAKQFLPFLQRAGRSEAVVEFIASGSRLRLY
LPKETCLITLLIS
**>Tnrc-6**
MQQQDQENTVSVSTWQSMGKSDSASDAGDDSSSVKSGSTITNMSNMTPQSQENTSSNNSTALQSNSSSLWSANNSTLSGIDSSPWGNSGSPAPSSINSMG
WTQSNLSLSTNQGHNSMQQGSSMISSNSAFNSTGSSSSSSWPSSMQTNTSSLPPISSHWPGSSMLGIGDFTKTDWNNPTTPIDTLSKDIGNSDPRMWGLSSD
KSGDSQWDISKSQWGNPSLSATGDQGASSTELSFAQATLKGLKVPPVNSAPQTVINSKQEEILRAIENHEGWGSRPIRQDTSWDVDNSPKSHRKFSTDSNAGAS
NVWNNSNGTAIWEAVRENQSGNWEAAPAGNTWNAEKEQAKWSFPPKPLQDPNTWTVGNGNDPKTFGTWGAAGGAGDASNKMWGQKTEIGSWGESG
GGGVQRTTSISSWGDDGDAGGWDDQRRVTSGMGSMQAIVPPSPGMSGQTVVPNMVSQNITGLNVPVMGGADVTLWNESPKPGWNPGNVTALTRTKLDE
QWNKPPPNRTGWGDPTQDNVKVDDGTSIWAANVPKQLPPQVKQSGWGETAPQAQWNATVGAKPKAPAGLDEASWAMAQRAKQMPPKYMGSSNNPPT
QMRAKLLQELMDMGFRKEEAQNALITNNMNLKMALSDLKSSNGGMSRRDMDIDMFQSNGSQSRLSYMSGMGADDLTDIRPDQVPTFSSLQNTQFPNSQV
PNQPFMPNNSGMPPSGLNASSSSNNSSLQQKLMQKMQQQQAPPTPTLGPRGQMPPVGSTSNNQVPQQQQQQQQILAQLRQAVNNGFISPQLLNYQLPH
NILVLLQQLLQLQSALQTMVTKQQQMLQQARAAGNRSNPQLEQMPGIITNINQQIINLQKQLQQAQNNLFSSQKPPNISQPPSGLMSSHSGQQPISNIAENLD
ALSSDLANVALQSQSRLTTQWKASSDTSADGTTTPTTGNDENGENKAAGAKGMLQASSPNMNLIPGGLGMTGDKTWSSNMSATSSSNWPLSSNDSAGTNSQ
SDLKMPGSVSTSTSSILSGMPSGLTDVIPEFIPGKPWQGLTKNVEDDPHVTPGSIQLQRSLSVNRVHDDSLNNLDGHKLNSGNSWGVSSYKSDNSLGLSQLGSRP
PPPPVMLPGKSGGQQQWQSGFNRQASWPHSSNSAFTKVGGSNWNDRTAISSWILFKNLNQSVMESTFRSFCARPGSYNNFFYHVPMALVQYSSGELAMKA
QQQLNNQPLGSAIINAEFIGENDAQRLASQFPPPQPSVLPPMNTSPWSQAPPSALYQNSGRGDPWGNSMQSGQPMKQFGSEAWNLWDMNDHNSNPLLN
NILGGESM
**>Ccr4-Not**
MKKNMTDISIAEFMQESGYSCTVNVDECRNLLALCGAREITPVAVARILGMMARTPVGLGDHGQQQQGDRPDVPGVSTWNVDVFMQVVKEMNPHLNWRE
VVVNLDHPKFLVFGRKGLRLLVQAIFRGLHQEPFPIDQIYKHWSNTEGQLSFLMQALKHPDVLCLADHKGDTPTVVIDILKSAPNEDDREIATWKSLSLVETLLRLSD
CGHHNQVIELFKAPIAKCPDILVLALLQITSAVNRMKTEIISLLMPIFLGQHSNSAVILHYAWHHQGHSSSIRTLIMHSMAEWYMRGEPHDQVRLSRILDVAQDLKA
LSMLLNATPYAFVIDLACLASRREYLKLDKWLNDKITDHKEAFIQACVTFLKRRCPQLMGMPLKEEPPQARSQQLPPETVVTMLNCLRMWTQHLNPELAETINS
MLNNASIYLSKPRPPPPGAMPGIGPKQPPYPAQNMLDSLSNIGGVLNQAGPGVVGPPGTQNLPQAIANLTLGPPVPPGSPSKSFPPLSQQQQAAPGMNFNPM
MGNPMPSIGMPTSMPQNIGPGPNAVPGPGPVPRPPVNSLPSLGTATAPKIGMPMNQERLGNRTVQPGNPASDMSNIFPEMTQTFSKEVEDRANTYFQQIYN
QPPAPTMSIDEVLSMLKSFKDAQEKTNRDVFACMLRNLFEEYRFFPQYPDRELQITAQLFGGIIEQNIVTYWTLGIALRYILEALRKQPNSNMYLFGVASLERFKTKL
KEYPQYCQHLAQIPHFSQLPPHLIEYVEFGSRSQEPPHPQMPQGMLSEISGRLTPVSMFHGLVRPSMPLDGQNVPTSGSALSASALPVTATPSTPTPISAAPRGAT
FAKPSIATATNIDTLLAGQGNNDVMVPSEAIQDKVFFIFNNLSLANMNQKAEELKEVMGSDYVSWVAQYLVMKRASIEPNFHTLYSNFVDVLGIETLTSKVITETFR
NIKVLLRSDKGVANFSDRTLLKNLGHWLGMLTLAKCKPILTTQDLNLKALIYEAYQKGNQELLYVVPFTAKVLESCAKSKIFCKPNPWTLAIMNVLAELHQEPDLKLN
LKFEIEVLCKTLNIDISTELKATKYLRDPTRMATLEHQLSPVKPQEEPVPSAVAQPPAGLPMPPIPGMPPMHSLPPPPPNNISIPPPPPVIQLSTGPANTTLPLQPPKF
NIHDINTNTNLASFLQQYIVINEQVIYIQANVSLRNSVIGAIESTMQDFIVPITDRAIKVAVATTEQIVKKDFALDSDEIRMRTAAQAMGRNVCAGLALITSREPMVQ
QLKLAIQQALVSNIRGQQNASQLKTIEDAAVILANDNVEICANYIQKLAVERVGAEMDKKLATEYEVRKRRRQEGIRYYDPQVFTYQSERMPEQISLKVGSVTSQQ
LAVYDEFSRNIPGFQPLTDIVPPSGSRPTTSMSGQEELIHLYEKMVSEIESHMHLMANPTLFIPHLQQLMQAINVARNTLDKNTATQLVQKAVEGLTEQYAMTNM
ADGETLLRFRECHLLVLKTFAEHRAFGMTWIAKEVTRCLMQSREDNRYNLDVVELLIRNGFVHMPQYDVYLSSLMENGLNQIGINFTTQLMQRLCVEDKTRAPQ
QVICSESDFTNSIEALNMIGSRSRQTQPDSLPALIESLRVGGSSSSDPSSLMDRVPGSVTSMMQTGITQAREFDDPAGLHEKTEYLLREWVNMYHSPQAGRDSFK
AFNAFVQQMHTQGILKTDDLITRFFRLCTEMCVDLCYRALGEPGQSTTMIRAKCFHTLDAFVRLIALLVKHSGETNTVTKINLLNKVLGIMAGVLLQDHEVRYTEF
QQLPYHRIFIMLFIELNAPEPILEAINFQVLTAFCHVFHILRPAKAPGFAYAWLELISHRVFIGRLLALTPHQKGWGFYAQLLVDLFKFLGPFLRNADMTKPMQLLYKG
TLRVLLVLLHDFPEFLCDYHYTFCDVIPPNCIQMRNLILSAFPRTMRLPDPFTVNLVVEHLPDINKAPRILTNIVSLIQPASFKKDLDSYLKTRSPITFLSELKSNLQANP
QGSRYDIPLINALVLYVGTQAIPLLQNKGVSMSTIAPSSHMDIFQNLAVNLDTEGRYLFLTAIANQLRYPNSHTHYFSCTLLYLFRQANDQTIQEQITRVLLERLIVNRP
HPWGLLITFIELIKNPNFKFWEHEFVRCAPEIEKLFENVARSCYQQKQQTPQQGPQGMLREETQVD
**>Pan-2**
MDFPNLPSYRPPHTQDTQNMPPPPTSFPPQNMHRDQMGPNPVFPNTAIPPRMPPGYPPLGDPHMPPGYPMVPSQDQSMSMGYQSGPMQSPGLPPGYHP
HMGNPITSRPPQNLQQSFQNSNQGTNMSPQNFHQQPPQHHQRSNTYHSHPTQQHHLDFRQQQQRQQHQQHQHQRQHQQHQQHRQQHGALINEFHH
PNQTTPGEFHPQQNNLAQDFHALQQNQVVSDYHQDFGHHDQGLSHHDFRHSRDHQGMMGGYMDHNTLGMQALMGDPNLMNQMDGHMLGGGDAYS
QHHIMQMYGDPALQDLREGDYIPMGCQMMDKTANASITCVNFDGHQELLWAGNQRGYVASYYGTDFFKYSAFRVNFEEDVRMNLSFPQGVLSLTPSLLRGTL
RRGLSVFTFKDEHLHDMTCMMMRGDTSVLIAGHHKQVLEVDIIQNRLSNVYEVKDAGIAIFRSTDRYICGGDSVGKVTVYDAKTMRAEHTLEAHSSCLSDFDIHD
NLLITCGFSNRMNTLTPDRLLMVYDMRVMRAMAPIQVSIDPTFLRFVPTIPNHLIVVSQMGSFQLVETITGGAPNTRLSYPVDTHGAYLVAFDVSQTYQALGFGD
SSGNVHLYSASPNSLFNPYSEPTEFADPVEPYDPIDINDELAAYSGIPMPYPPQGPLVSDFPEHMMARTYRKPLPIDSEVLRTMKVYQNIGYAPNTNKQKRNLIPYN
LNDKGNKDRKSSIPESPMGRGDDPFMTVPKRYQRVEIKYSKLGVEDFDFRHYNKTNLAGLEPHIPNAYCNSMLQVLYYIEPLRCALLGHVCEREFCLSCELGFLFH

MLDLQKGNTCQATNFLRAFRTMPQVSALGLVLHDADESSGRVDFPRLIQSWQRFVLMQIHEEICVPVESVDGGVSDTKVATKGGEAEDKKGSKKKKGRKNLKEA
SAENSENSETKEGGTEGETED

**>Pan-3**

MQQQNNPIGNGSFRNGPVHGQAQEGIARDAMGTNGDQYFSATFNAMGIQDFPPFRGGPPRPRSSMGAPHPMAGKQYSSTSAMDQMPDDFAALNLAVGP
LHNKGAVNDFEPKSPGLPHSSSTSSFASYASMFGQNPSAPAWPFVPSSVAPSAHSPGSHNPAVFNAGVQNGPTTPKTSSIVPAFGPSQQNPVSTPSNATLGPPLF
GSSSQIPVATSTNPVSTNGLVYTSAISPIHSPGISPASSPLTTRRMQSPVTHSLRQTTIPQKSASSSLVQESVGGTTYFYHPDDFKPQIEGPQLTLPSFCAYPGMPSHIV
HLKMKTNVPQYFAPDELKMDILNRHVLTMMQVEPTSPNAVELPTEVDKYHNLCPLERTGAPKPSLGNLPSTTYKAVCTKDGLYYCLRRIHGFRLASTKVNSIMES
WKKLYHPNVVSLKEVFTTKNFNDNSIVFVYDYHPGAETLMSQHFSSHPHINGFNHHHHHHHHFSRDHGKNSSNARNMNNNLLPESLIWTYIVQLSSALRAIHA
ANLAARSLDPSRIIVSGKARIRINCIGILDIITPEGSTPPNINAYKQDDMTAMGKIILALACNNIMCVQRDNIQSAIDVVQRNYSPDLKNLICYLLQNTNRVRSVNDV
MPMIGARFFTQLDTATLRSDVLDYELAKEIENGRLFRILIKLGSINERPEYTTEPSWAETGDRFMLKLYRDYLFHQVDEAGLPWIDMAHVIQSLNKFDAGIPERVLLV
SRDENSMLVVSYSELKRCFQSCFQELVAQHNQGTTSFS

**>DGCR8**

MFLQKKKICFPFYHSILRSWRERRKHLGERNRRSRPELPPSTKLITCPIPGNPKADGNGDNVKKREFILNPTGKSYLCILHEYMQRTLKIQPLYVFKELENSKTPYGAT
VMINNIEYGTGYASSKKVAKQEAAKETLKVLIPDLFKKITDQEIKRNISDLSFFDDVKVTDPRVNELGNKVGQPSPFQLLLECLRRNFGMGNTQCEVSTKPLKNQKC
EFTINVGKHSATVVAKNKREGKQLAAQAILAKLHPHVPSWGSLLRLYGTSVEKPIQKPEEMNDVKSHVPNHSVLESLRQEMRKLHRQKEAIQSKGKIIISSKDLPCK
MSGVD

**>R2R2**

MSHNHMLSNQQQQHSLMNSGATASGGQQRTMIAQRGSQVLAQPKSAYQLQPASVSQLSHIQQQQLVNNFHSQKQMSPDISSTHQYNSAIGGLPSQNGNYQ
SLTSTSLHSNITLEGNALSVQHQHHQYPQTQQPQPPIQPPVPQHHYQSASVSQPQQPSQHHLHQSMQQQHHHLLQHQQQQIQIQQQPIPQHQSSSYQQHQ
QAQLQIQQHLQQRSLHQQQRFQKQQQVQPKQILQHPTSASKNVSSQIQQPSQPSTLQLQPQQQSKPLIINGQSGITDSKKESSDKVEVKLDDNEESEESKNNAT
EMLGLANTKEKTPMCLINELARFNKMSHQYTLVDEQGPAHKKTFYVKLKLGDEEYSASGESIKKAQHAAAAIALVETKHPHPPPKPARYSGSNSVDSGDENITPTV
ELNALAMKRGEAAIYKAIEPQQPPYYHQPGMDYRGIYSQRYHQYMRASRDPRYRGSGVLWPLRYHYPRMNRPFYVSLRVGHREFIGDGPTRQAARHNAAQKA
LRILKNLPVQHEAKKDQEQAEEESNDSLKSEISLVHEIALRRNMPVNFEVIREAGPPHMKNFITRCLVGEKVTEGEGNSKKTSKKKAAELMLEELRKLPPVPTPLYPR
PKSKIQLNKKKNRNLIKSELQQQKADPNYGVGINPISRLIQIMQAQKKKEPVYTLVAERGLPRRREFIMQVEVEDKNCTGTGPNKKLAKRSAAEAMLQILGYTKPSP
QPAKSAFKHPPSGEGNQSNGDKKVTFVEGDPPAESTEKPSGPTQSIQQRVPGLLHLPTSNSKLVMNSAASSSSSLTQKESLVNLASILKPNLRPEVQLRELCKALGQ
ELEIDDFTKKRQTGTEHITRITVGKEPSQNFHGSSNTLESSRDMAALDALKVLLAKVKDIHPGGDGPQVKKELLTRSSSGMKKDISK

**>Taf-11**

MTEMDGISPLLDVLSPLEVPTKTTVMTQHSLSKELIIETDISVSSTPSASPSPSPNPSLSPTPSSLQRDSTPVRLTASDKQASPKSSRSSTPEKRSSSERERSSDREKTPE
GGGAGSKEKAQSTEKAGPSGTPGPSGAAGPSQDKKRKLEMSEEEVKQAVKKHQKQEEERMKMQVLVSNFSEEQLNRYEMFRRATFPKAAIKRLMQNITGTSVS
QNVVIAMSGISKVFVGEVIETALDVMEGWGESGPLQPRHLREAVRLLKSRNMVFTNKTKKILF

**>Trbp-2**

MTVPPGKTPISYLQEYATKHAITPQYDLIANEGAVHEPTFIMRVTVGDNAVATGKGSSKKKAKHAAAQNALNILLGVTNGQEEIKEEPTVSTAPTPNKEEDVGNPIG
ELQEFTQKKLLKPPIYEFVTEQGPPHAREFICNIKLGKFTDKGTGRSKKTAKRMAAANMLAQLKALSQDKEAEKQLEDSDEDEEIPLGFEKSTFTGLKQSKGKAKPV
NPQSALEIQRFYEKVMIAGGKQIRGQGQTLTPPTNYCQMLQEIAEVQRFEVQYFDIKDDSAMGLNQCLVQLSTAPVAVCQGTGPIMDEAHANAAHNALHYLRI
MIK

**>Pir-1**

MAGIVTNFFRRTHNYVKRVMPPPDRWEQYCPLGKVIPGTSFIAFKVPLKETLLGQIEEKDRFSPKRLVQTLESEGLKLGGIIDLTYTLRYYDKAEFENLSIKHEKVFTP
GHEVPNYDVFHKFAKAVKTIDKKGLLIGVHCTHGVNRTGYLICRYMIEEMDMEPDTAITLFNEARGHDLERENYLDDLRKRKKGESTYDPNYQPKEDPAKPNHNG
GRGKGRWRHNRSDHPNWRQQNYEESNETKPLDGDSERLKESSNDFHGSFKFNRYGGRALDPGFYQEGYGRTHWGYDDMRHRGREYNAKQSNREYEDTRYS
DRGYDNTRGRGYDDARTSRREDDGARPSRREDDDDRSRGRDNDVPKWDNSNYKSKADINKHQESDTVDEKSGNVRKPQKRKSGEGSLVDSSVTTESHSCERK
KRKKSNKEKKV

**>Pact**

MTVPPGKTPISYLQEYATKHAITPQYDLIANEGAVHEPTFIMRVTVGDNAVATGKGSSKKKAKHAAAQNALNILLGVTNGQEEIKEEPTVSTAPTPNKEEDVGNPIG
ELQEFTQKKLLKPPIYEFVTEQGPPHAREFICNIKLGKFTDKGTGRSKKTAKRMAAANMLAQLKALSQDKEAEKQLEDSDEDEEIPLGFEKSTFTGLKQSKGKAKPV
NPQSALEIQRFYEKVMIAGGKQIRGQGQTLTPPTNYCQMLQEIAEVQRFEVQYFDIKDDSAMGLNQCLVQLSTAPVAVCQGTGPIMDEAHANAAHNALHYLRI
MIK

**>Ddx6**

MAKEEMRQNDTSNEQGWKSKLNLPAPDHRVRTSDVTNTKGNEFEDFCLKRDLLMGIFEKGWEKPSPIQEASIPIALTGRDILARAKNGTGKTGAYAIPCLEKVDSS
KDLVQALIIVPTRELALQTSQICIELSKHIGCRCMVTTGGTNLKDDIMRLTEPVHVIIATPGRILDLMNKNLVKIQNCGTLILDEADKLLSQDFKGMLDSIIGHLPKDRQ
IMLYSATFPLTVEAFMRKHLKNPYEINLMEELTLKGVTQYYAFVQEKQKVHCLNTLFSKLQINQSIIFCNSTQRVELLAKKITELGYSCYYIHAKMNQQHRNRVFHDF
RKGLCRNLVCSDLFTRGIDIQAVNVVINFDFPKHSETYLHRIGRSGRYGHLGVAINLITYDDRFALHKIEQELGTEIKPIPKTIDPSLYVAEAQQIDPDMEEDFYRQQQ
MQHNQQMQQRANSSHQQASRGPPPHQQQTVA

**>Eater**

MGVRGNLLGVTQSPNRSTTYVISKQAKISMSVSRIFEVTLPSEFSVLAKIKMPSKKPRGYFFVISDIQGRQQLAIYLGKRLKFQYLGYNYAFKMDIFENTWHSIAISVD
KNTVTLYADCEMVIKKRLRSKEEHLGTNLMMSIGPYFSQHGIPFEGEVEQLVFTSNPEVASQQCGLVVLDKKAANSFYEDSTQFRYTTVTAKPIVTSYAPLACSVWI-
MVGLVHLQRHLRTGVANKSAVLSCQHSQCSKLSPCFNLASPNEALLLGELPRMFPSLLQWWNLFTWQYLQM

**>Dcp-1**

MILNRKDLNNLVQPLTSQVEFHLNTPFLLFKKLSDDIEDGKFQMFCKQNTIQGVLHTTSG

**>Dcp-2**

MDIEVQKTTFGDESKSIPSYVLDDLSSRFIINSRPEEVTSIIRIFFLVENAYWFYLDFHRAENPDLKECSLKEFALNLISHCPQLKQYTLEFDKHFEAWKQYKRTIATCGAI
ILDQDLKHVLLVQSFSSKNSWGFPKGKINHNEAPEDCAAREVLEETGFDIKPYMDPQEYLEKFSNEQISRLYIVAGVPINSTFLPKTRREIRDIQWFPMDTLPTHRGD

QTLKEAVGQNLNLYTVTPFLRSLRIWIKNRSLEMSTMTSKQRNRQKQQKQFSQPSHGVYQEFSQLKKGKSAAKNLFTQKTSTSSVRYSKEIDVQEANVKQSKRAM
TVFQSLFGTKDGRPVADFKAPVVDTNSFHSKVWHDFVIDEDALMEGFPHEGTYCLPFGQQRELVAGK

**>Vig-1**

MDHQYGIAVNNKFALFLDEDEDPLEVLSRQEEQAKKKKDEGEKKPSKSKQKKTAVPETKAKVPEPPVVKKEEKTVPARQSERGGRGVGRSREPREIRDNDGEKRP
PRRQAPKENREIRVGDENVPPEFRERPESGFRRREDRGDREGGFGGERARGRGRGRGRGRGGERGGFGGPRGGFGERGERKREFDRHSGTDKTGVKPIEKKEG
GGSYNWGNFKDDLEEPAPQTETNEWANQPEPGTENPELNESAEGEQAPEEDPQPQEMTLDEWKALQTQNKPKAEFNIRQAGEGEDGSRWTKGREYHKKHE
EEDDEEEESEEEDEDDRHARGKHLVTDIRITFNDTPRRGRGRRGGRGGMERGGLERGGRGSRGGPRGMGGKPRESAPRFDDEADFPSLVKSAA

**>Zfp-2**

MGVHCAYGTCNSDSRYKDRPHMQGVKFYSFPNPQKDLEKCKRWVDACSREGFFVHSLSKHMFICSKHFVGGKGPTEKFPDPIPTNFTDTQLKKFSKHRKQPLVR
PTLQLPVATTSQCLDEENSFVNTGSVSVQGTLDNRKTEKVEDFKESYGCIKYEKMEDLSNEHNEKEKSFITNVDEQFVNVHKIQNFKEEYPLVKSENEEDSCNEEEKI
LTTDKVEEYGNIHKIEHFKLEYPLVKSENEEDSCYEQEKMFTKKDDDNVKVPKDGDFKEVCPPDTGNSNGINYVLNKEIYNGDCSIVKREREEDIDMEENILPTHLLE
ALTKSVNGEKQINCEKQTNCEKQTYSEIQANSEKQKNVVIQKNVMSLSPSSKNEEKTLSCDFCPAFFSHKSKLKRHRGLHTGETIYKCKNSPVALALKLCLVKRMQTK
EKPFKCEDCPSAFSLKTDLVRHQRLHTGEKPYTCDICPAAFPLKSSLVKHQNIHTGEKPYKCDVCLSDFALKSNLVRHQRVHNGDKPYKCEDCSAAFSCKSSLVSHQI
VHTGEKPYKCDICPAAFSLKSSLVSHQLVHTGEKPHKCDICPEAFTLKSSLVSHQLSHTGEKPYKCDLCPAAFTLKSSFCKHKKIHSGEKAHQCDNCSAAFIFKSRLVAH
QRMHTGEKPYKCDICVATFALSTSLVRHHKVHSGEKPYKCAICDTAFPLKASLIRHQRVHTGDKPYKCDICPAAFSCKSSIVAHQLVHTGIKNHNH

**>Xpo-5**

MKREWPQLWDNLFTDFTVLCQNGETQTELILQTLLRLTEDVVRFQNLPHSRRRELLQSLTSAMGSIFQFFLYTLNKNLKVYQSQSGKTSQKARKICQSVLDTLTAFV
DWVNISHITEGNLLPLLCSLLLDKNLCLRASECLLLIVGRKGKLSERKPLMILFTEEAMTVLLQAANNATEHITESTNFLFLKRLCEILLEIGKQLCTLWGSSEDTGQPQ
NFEMYLKALLAFTQHPSQSLRQMIFSMWLIFLRHPIASKDSIFQSVLPALLQCGTVCLHKVGFPSQHNSISCDYSRLEFDTDEEFNVILSTLRVSVVESVRTMTLMVP
TLTFSVASSWLKDLLNKPIEIGTGADADRGICNLSSPSFIAWDACSVFLEAVMSKLFLSDGDKPNVHEGIDLLHRVLAYQMQDPLILSAVLSCISGLFPFLNYTPQTLP
QVLAKIFDAVVFNLPGQTKSTRSQAVKNVRVHACSVLVKICKNYPDLLLPEFRHLYNSVKQLDSDREQLSQMEKIILIEALIIVSNQFHDFARQSAFIAEVIAPVKELW
SSEDFNKAFSTPECFMSYVGLDQAAVEPSSADTCGINRSHITYCINTILAVIKRSQWPEDFAVAQKGGFLFPADAGSVLRNPATPYICPLLDNLIILLKTTCCLFKTEYLQ
LRHSDFIRAYDLMDHDRLAILGIPPACVDNSDSLVYRHPLERMQNFISAVFEYGFHILGNASQCLGAEFYCAPGLSRVIISNVVVNFKLLPDFRARPFIRNFMKPFM
QWCPREQYATVAIPVLTILCPDILQ

**>Wht-1**

MDKRLDRSARLSRVEAVITEMGLIGCANNRIGDAGGGKKGISGGERKRLSFASEALTNPPIFFCDEPTSGLDTFMAQSIVSTLQKMASKGRVILCTIHQPSSELFSMF
DQILLMAEGRTAFMGTQQAAMEFFQKLDYPCPTNYNPADHYILTLAIVPGREEECRENTLAICNKFKETEECVKITRQTQELTETARHSNSDPLLNDLLAGDSRYDS
SWLVQFRYLFWRSWITTFRDVILFRVRILQTLTLAIVLGLVYLQTTVDQKGIMSINGAIFLLITNTSFSNMFAVVNSFPLELAIFLREYGTGLYRVDSYYLTKTLSEVPLFV
VISIIFTTLTYWMVGLYNSWDAYLIAVGILLLVANVAISLAYLI

**>Mrp-1**

MCLVLKILILERTRGVRTSGLLFLFWFIYLICEIIRLRSVIKVATSQGYVSPSIFIHYPCVLFALILSMFVDAKPEGSISSGDENPCPEKSSSFLSQLTFWWFTGMVVQGYK
RALVQTDLWSLNKEDTTEYVANAFYRKWNGTKEIIHTPVANGDVHETSFMYKEGSAILVPYGVGFEESQKPKRSLFHALVRTYIWFFLLAAIYKLIYDILVFVSPQLLK
LLIQFTSSDEHMWRGFFYASMLLLIALLQSVLLHQYFHATFILGMRLRSTIIAVIYKKTLRLSNTAKRSSTVGEIVNLMSVDAQRFMDLTTYLHTIWSGPFQICVALYFL
FQTLGPSALAGVGVMVLLIPLNAVLANKSRQYQVSQMRLKDNRIKLMNEILNGIKVLKLYAWEGSFQDKVLKIRNEELRVLKKAAYLNALSSFFWTCAPVLVSLSTF
AVYVLSSPDNILDAEKAFVSLALFNILRFPLSMVPNVITNIVQANVSLKRIQNFLNHPELDPYCIDKQDTEDGLAVRLEDASFSWEEGNITLTDINMKVEEGELVAVV
GAVGAGKSSLLAALLGELEKTSGHVSVKGSVSYVAQQAWIQNATVKDNILFNKPLDQTYYDSVVKACALTTDLEILPAGDLTEIGEKGINLSGGQKQRVALARAVYQ
DTDVYFLDDPLSAVDSHVGKHLFDEVIGPSGMLKGKTRILVTHGIGFLPQVDKIIVLVKGQVSEVGSFAELMKRNGAFAEFLRNYLTEELKDENSVAEDIDVLSKREEI
MSQIESMIDDTSALQRQVSKISERIRSDTEVSKGSNTSLPTTMTSPSSKRHDKSDIIENDEEKKLLPEEPTEKKGDKLVQAETAETGRVKMTVFMAYVRAVGAFLSVL
ILFFYILYNSASIYSNIWLSQWSNDARNPNISADEDQRNMRLGVYGALGVLQGIFVFITALMRLLGGVQATKILHSGLLANVVRSPMSFFDTTPSGRIVNRFSKDVD
TLDTVIPMIVGMFLMCVFQTFSTLLVITISTPMFLTCIVPLLIFYYFVQRFYVATSRQLKRLESISRSPIYSHFGETITGAMTIRAFGQEALFIKDSQTKVDNNQICYYPSIV
ANRWLAIRLEIVGNFIIFFASLFAVIGRDDLSPGIVGLSITYAMNVTQTLNWLVRMTCELETNIVAVERIKEYTETPTEAAWVIENNRPSPGWPDKGVVEFKDYKFRY
REGLDLVLKGINCTIQPGEKVGIIGRTGAGKSSLTLALFRIIEAAEGTITVDGINIATIGLHDLRSKLTIIPQDPVLFSGTLRMNLDPFEAHTDEQVWSALEHAHLKTFVS
GLEKGLLFECSEGGENLSVGQRQLVCLARALLRKTQVLILDEATAAVDLETDDLIQATIRSEFANCTVLTIAHRLNTIMDYTRILVLDAGRIKEFDDPKVLLQNPSSMFY
SMAKDAGLA

**>Abt-1**

MDVIYALIWAPDPYATFCADKYIKYLELPPNVPEDEISGLVCDTNWTSAVDQMTQPVKNIVERITEIGDLFALNLTRTYNVENDWIDMVDYTERIFDLFNSGDFMK
MDFTLGYKAVFDQMNFTKLGDGFDVLFGFFKEININDVEKVGNLTLKILEKLDSTYGSTSGWHTIRKYIVLFDAFLDIEINMTQEFENSKSLLDIVTSYPPEIRQIITLM
AKAYPEFVTAFKNILLDPGRLIDKFLSDGFQAPDCKVHFVTDYTLLTSDSAMHKLEKYLCELNYQALGQEFQNTWPIVNKFSEKLTLILNGSDFDTAVNWEQLAIKTN
KFLDLLGYLGNAPLFSDGFDFSPFNLTNIDLTWQEFYTAAETLKYFDMEKFQNVAIKVQELALKLLVNETSSDVSLVLYSYMYVTHHVLKLANTELEYFNGSTEVVMS
EYLWSSELNKLLAVLSKSPDINAIGVATIQRLILSPGEITIPDNFEDLCTDVVLFSKVFAVEASPLDPAVIQKTVCDLNLDFQLILDQVQNNRPSVHEFVSSMKLLASNF
TIDQIRVDPSVITKEYYQMSSLIKDVVFKPPEFKLTPNQNWMNLTLYTQEWEKLSQKLNILGQDLQNSSKLSDWESSVAKMLLTTVEKIPGAEKALGYIDGILTLLER
QLNTIEGGTKGLKNYTNANLILELLSSSPEAVQTILYTVMSEPVKTTRWGVAFQSWAIFCSTPASDIMTVPSGLNFNISLFLEKVCRIDIEELGKELTRYEGIDKLERLFT
YGPDTSVNVTDVQKKLDSFIRAMTDYIHNPSDVTDNSLEWLFNFTIWEEVGSRINVWSHNVGAVYSSPDKIAGMALEIIQIIFGNMSKDYTEIYEKIVGVSDIILGQ
VLNILNSTRLSDSFHDLPSLHTIVDLLEDLPELYETALYTSMYHPERITAKGSDFRSFEVFCSNNPQEIFSNAPGSTFDIKSWFTRFCAINITKMVDELGNYSVAKDIDKII
SGPYTEPVKISSVIDKVEKIIEKFNKGIDVQEPLFDSKVWQQVMDHVNTWLWPYNAFTEFLRDMPTRWMKAVVAYFENNPISVNVFGYIDAVLDIIINQLDETQN
GMQGLQNYTNFNMILNLVHSAPEAIQTILYTVMSEPIKTTLWGQAFKSWSTFCNTPANDILTVPPGLNFNVSLFLANVCKVDIEELGQEMSRYQGIDRLNALFTN
GTTGYVNSSVVVEKVDKLFTTLSDIIDNKVEMQTNVALSRLFNTTIWEEIGDRIKVWANSSEKIYLSPDNIVGMTLESLEVFFGNMGQSYKGIFEKAIGVTDIIAGQIL
EILNSTSLNDTLHDLPNLQTMVGLVENLPELFETALYTSMYYPEKITAKVSDFTSFEVFCSNKPEDIFTLAPNSRFNIKSWFTNLCTINVTKLVEELGDYKAAMDFKSIIS
GSSTVKISSVIDKVEKIIDRVNQGVDIKGQILDQDIWKHVMDNVNSWIWPYNTFTDFLKDAPMRVMKSVIAYFENTPGSNKVLRYIDAVLDLISNRLDNAHTNLTE
AVKEFPTLRTIIELVQQSPEMIETVVYTVLTQPIKTASWVNAFQSFSDFCNTDVSEILTVAPGSSFNVEGFIKKVCSINIDNIATEITSFEGFDRLRRIDSENDTSAINATT
LLEKADLIFHSLDYYYDHPLSIITLDLPLFNETVWNNMGQRIEKWAGTSEQVYLNAKTISRMFELMFGHAISMTSELKDAYDKAVSITSVLVDEVSMVINGTSLEEVY
TIPSMRQLIKLANKAPEVFETLLYTTMFHSEKVARRALSLVSFEKFCSVNPTEIFTVPPDSSFDITAWVNDLCELNFTVVETELANYRTVKHIENIVSGVGYINTSLSSV

VDKIDKLVQQIQQGFNLNDHFLNGTVWTQVISRLERWVGNYTIYDSPLMNFDGIFDLIEQLIDTRPGVEEELKPFMYVGVILDNLLDKMITLENKTHFGAGEIVEG
WTAMQKLLNFIQRPDVLDIIISSATSEHFLSLFLNTSSLTDFCAPNTPVTNYLVAPKAGVDLKQFKSAFCDIDFENFLNHFDDFFDTKKLERVINGTEKFNWTKLQDK
VRRVTHLINDKWIQNPPEFDLPSHLTNVTYWTSLFEGSIKSLGQRLQTKEQIENIIHQLKPLLELDGFKQMGIVLNSVLDILNENLAALQNTTLTLGNAVTKIPALREF
FDALGLDSGTLESLLMAPVTNLTKLAEMFLSTNMEGEFCTTNKWKEIIHLPDSFNTSSLFYAVCDGNVTPLLDNLKHVYNLQTMIDALNDTSSHLQDWDQSVDKIF
KMVDYIKILTENPPQISTDSTLEKLKASYTNTTQLWKMLYAFSALHQAFINDSVFTGPFEGVFRVSNVLVNFLDEMVARVKIQNGRLDLATLFKDVPEFVKIVNGILD
MNPDPLTGLFAVQLKPTEVDKFIDIGQNSTELSQLSCSQSKFREIFDVRADVDITSLLSALCAIDFGPIAEKLRETFKINTIGDDVINAWNSGSEFDIVNFMTKMEAIID
KITNLTKVNSIFFNDHDLGNVLRINATRLVEELKNQYTNIQDAFTQDYVSLSASILNSLLSNMQNETEIKTLTLSLQFAQVYLHSFNTFLRSVQDTPISLVKLLNNTEVG
SIINPILNDPHILYALVNSTLNIKELDVLLKHPFPVEALCGEQLWSAIEVPTNELEPFLKLQKSICSANVSLIMWQTIMLQAKGYQLYKQLDILKEEIAQGHSSLNVSAVG
NEMKDFVSLMTTIINQYIDGSRSITDLLDVKGFQDILDSVQRTIAQKLINSAKNWSVDIAELVLPHIMDQTSLLSVAKTVNTVNIFMNVVISKLQGIRNGTISSSTLFT
NADGLANLIEAYIRVGKNIAQSWIDGEFDVDKIQKLFSNGTTQLEILCGQTDGIAKNLTSLASASQTIKDLSTVICSAVTESVLKDIQSFVDYNAIINETEKIWRANQQ
TPDFTEFANNVEEFTNIIKDLASKSLVVSSNLSSSFGFTSTLDALKNIATDPKLIFRMLKLLGIVVEAPLRKEQTVVQVLSGLNTFAIMPVVNILEGLRAKGLTLQSIGSN
PETLLGVLSTMVDFDVQYSVMETVIIKSQLNSFISNRDNGLCNSTNYEDVLLSRGLAQVELPVPWKDIYSIMCTLTPDADVLLQKLKDLGLTKIQIALVLEKIMPVGQI
ENLPQMYDHSFGWIDLMSSLEQLSDLATQGQLNQTKVYSMKNAWEAVKDIVFDKSLKSLLSYLNLVEGEAEPNDSWVKFKQFLRFFSTTLNFIDGNMAKFAQGK
VIELKDILPDSDRVASLLENVFGGSAAELLTTSINPTMFYRITMTDSWDEICHGNKSIATYFNFPYGTDIDAVKNNLCDAVTSRSTSFQQLLSLFDPQGIIKELDMLIH
GEYDASAHNDTVWDALYTSVMNVIHTAESLQNVQIDFSSVDGWLTPILNRLHILIEYDSFRITGMCNSLVHYLNNTEGYKAARQPLTMVITGLNLLTDVTKLVPELD
DLACLLVSKTGIDVIDVMSRVSDLGFWDAIGKAIDDFSDPPKTLDCSAPVLDILHLMESINASLRNDGLDVPKMSQCFQQAGEHIRVMLTSFGDTILFVKDLLSLIQ
NPALQRVLHDPNVLPIVDFILGTINANERVIISVSDLLQEKINVTDFLVNTLGLSQLVVDSFLNATVNNDWAFFLHEPLNLIVDILCDPAKLSKIISLPPESPLDVTNISSV
MCNANVTMAADYISTMSATVGNITNLISTSTVHVMDVLMDLIPAIADVLVKFTDIIQFSTQAFTSFDISDIKNNVDLIDKLMKGNGLSDLANSVHKILDTLSKVLPSN
SRTDMVMRDVRRILDGLLGLDIIKGSFLEEFQVKDLIKDSENMKKYLINQFGFTDLIAQEIMDAVFSSRVIVGVVEKDQPTCKDVMKMLIIMNTTTAIVQDVNNAI
CALNSSQVEQLLKDLTPELDIGVFIAKYVANTGKAFLTSANITSDELNEMIEKLDKGLGNLQKAADLMEQKKSGKLILEAFSNVSASSMISLDQLTPSICGRHVTDAFS
FPAAPGFIVSSGDDIKSQLQKEIDGNEFDLPNEFCMQLYEEIRSSSIGSIVWAYVKPIMRGKILYTPDNEMTRAILAKANSTFAGLEELRRVAAIWAEQSSNLNSLIDL
ANDTEDIKEALQNDFIQGVVEATAGIKSDAILTSLESLQSGKFDLSQIKNLKTAAEIIANYSSCILTNRFQPVASESEMITEAYKLTQNKTFFAGIVFLNADDSSNAQSSV
NRKRRQANKDLFPKHIGYKIRMDLDNVMDTTWIKSPIFKMTSESNFIEDLRYLRGFLYIQDMLDQAIIDVHKNEKLTSPGLNLRQMPFPCHHQDSFIFFLGAYLIPV
MMTFVFLASLGVATHNLVYDREDGQEENLHVIGMVKGLNVIAWLISTLLLMGIVCIIIAVMLKYTQIFIYSELSIMFLYLLDFCFSSVMLLYMVSSFFTRTTMAILFVLII
YIMTYLPYVIIIGLEVSTEVRVMEFWHKILASLCSTTAFCFGSLRLARLEENGIGIQWSNIDERVSNELSMSWVCFMMLIDSAIYFLVGWYVRNVKPGQYGIPEPLYF
PFMPSYWSSCFRSDKKGLAMADRQPNTQGSSFETPPEGYKVGIAIENLTKVYSNKKKAIDRLSISFYENQITTLLGHNGAAKTTTMKIICGVLKPTSGHVFINDQKIG
QSHINNLGVCPQQNSLFYYMTVKEHMRFYAGVKGARDTTNTAKEIKKLLNDVELWHARKTPVGDLSYGMKRRLCVALAFVGGSKTIILDEPTSGVDPHARKNIW
NLITRNRPGRTILLSTHHLDEADFLSDRIAVMHQGKLLCCGSPSFLKWSFGGGFNLTVVKKDQINMQKSEQGEQAESQEVADLRSKSVNNAVLAFIQNYCPDAKLI
EQVGSDLTFNIPKDPQKMTVPFYQFFNQLDLHQEELNFASYGLSDTTLEEVFLKLTGDADDAADSDIIDPVVTSRHRTAPLEFDDLEGAQTGKGYSTHGVDTSQRR
KGSHLLFAQIGALITKRFNHYRRNWRILISALILPMIFFLCAIGFSSILPNESEPAELVLDSSIYGPKMYSFYQDSVNNMMSNKFVDRLTNSDVGYGTACMKDWGQK
YFKESTCVATTPWYNYSTPMFMVGCRNARQIFTELPVPYSVIEKNVQPNDFIQDLNSWNVLSYLLRTFDEYKEKRYGGWSFERSNDGDSQNVNPYVWFNSKGY
HAMPSFYNSLTNTILRAQLPPEENPNEYGITAVNHPIKLGRAPLTIKNLQGDAADAGLSLAIVVAFSFIPCGIILYIINERVMKERQLQNISGIGVFTYWFVAYIWDLCM
YAITLGIAVVILAIFKTDSFYLRDNLAGFAAIAILYGWAIIPCLYCLSHLFNTGSTAYLVSFCLNLFLALCTVISLLVLQLFRDSTGVLEAYKVCKYLYLIFPQYSLGQGLMDM
ATNTVIYKLFRRFDDDRYVSPFSVDVLGWKLLALAIEGVFFFILTIILDVISRPALSLPPRHQSPDIYVEDDEDVIREKERIIRGNTNDMLVVDDLSKVYRRKLRKFLPVN
HISFGVREGECFGLLGVNGAGKTTTFRMLTGDLKPAGGNITLRGQRMSLNERSFGQNVGYCPQEGGLDEYLTGEEMLYFHSRLRGFNSDQTKKLVTDLLAKLGLK
QYAKRSVHTYSGGTKRKLALAVALLGDPSIL

**>Vha-16**

MSTYNMDAPPYSAFFGVMGVTSAISFCALGAAYGTAKSGTGISAMAVMKPELIMKSVIPVVMAGIIAIYGLVVAVVITQDMPTKDYTLYKSFLHLGAGLSVGLSGM
AAGFAIGIVGDAGVRGTAQQPRLFVGMILILIFAEVLGLYGLIVALILSVK

**>VhaSFD**

MYSLINNPKAHLILVTHLIEPPSRHPHAPFSTHH

**>Chc-1**

MFTHYDRAHIAQLCEKAGLLQRALEHYTDLYDVKRAVVHTHLLNPEWLVTYFGTLSVDDSLECLKAMLTSNIRQNLQVCVQIASKYHEQLTTESLIELFESFKSYEGL
FYFLGSIVNFSQDSEVHFKYIQAACKTGQIKEVERICRESNCYDPERVKNFLKEAKLTDQLPLIIVCDRFDFVHDLVLYLYRNNLQKYIEIYVQKVNPSRLPVVVGGLLD
VDCAEDIIKQLILVVRGQFSTDELVEEVEKRNRLKLLLPWLETRVLEGVNEAATHNALAKIYIDSNNNPERFLRENQFYDSRTVGKYCEKRDPHLACVAYERGLCDEE
LIKVCNENSLFKSEARYLVRRRDPDLWATVLSEENEFKRQLIDQVVQTALSETQDPEEISVAVKAFMTADLPNELIELLEKIVLETSVFSDHRNLQNLLILTAIKADRTRV
MEYINRLDNYDAPDIANIAITNELYEEAFAIFKKFEVNTSAIQVLIDNVNNLDRAYEFAERCNDPAVWSQLARAQLNQGMVKEAIDSYFKADDPSQYMEVVAVASK
SENWEDLVKFLQMARKKARETFVETELVYAYAKTNRLADMEEFVSGPNHANITQVADRCFDNKMYDAAKLLYNNVSNYARLAITLVHLGEYQGAVDSARKANST
KTWKEVCFACVDNGEFRLAQMCGLHIVVHADELEELINYYQDRGYFEELIQLLEAALGLERAHMGMFTELAILYSKYKPEKMREHLELFWSRVNIPKVLRAAEQA
HLWPELVFLYDKYDEFDNAILTIMAHPTEAWREPQFKDIITKVANIELYYKAIQFYLDFKPMLLNDLLLVLTPRLDHTRAVNFFQKVNQIPLVKPYLRSVQKNNNKAIN
EAINNLFIEEEDYQGLRTSIDAYDNFDNIALAQRLEKHELIEFRRIAAYLYKGNNRWKQSVDLCKKDRLFKDAMSYAGESRQIEIAEDLISWFLENGNHECFAACLFQC
YDLLRPDVILELAWRHNIMDFAMPYMIQVMREYTSKVDKLEQSESVRSEEEQKAEERPIVLTEPQLMLTAGPMGMPPQPPYGYIPGPMPGAPMPGYGYSM

**>Ap-50/Ap-2**

MIGGLFIYNHKGEVLISRVFRDDIGRNAVDAFRVNVIHARQQVRSPVTNIARTSFFHIKRSNIWVAAVTKQNVNAAMVFEFLHKMCEVMAAYFGKISEENIKNNF
VLIYEILDEILDFGYPQNTDVGILKTFITQQGVKSQTKEETAQITSQVTGQIGWRREGIKYRRNELFLDVLESVNLLMSPQGQVLSAHVAGRIVMKSYLSGMPECKFG
INDKVLMESKGKSSLDDTARGGVTAGKTSIAIDDCQFHQCVKLSKFETEHSISFIPPDGEFELMRYRTTKDISLPFRVIPLVREVGRQKMEVKVVVKSNFKPSLLAQK
VEVRIPTPLNTSGVQVICMKGKAKYKASENAIVWKIKRMGGMKECQLSAEIELLNTSDKGKKWTRPPISMNFEVPFAPSGFKVRYLKVFESKMNYSDHDVIKWV
RYIGRSGLYETRC

**>Arl-1**

MGGLLSYFSALFGSKERRILILGLDGAGKTTILYRLQVGEVVTTIPTIGFNVETVTYKNLKFQVWDLGGQTSIRPYWRCYYSNTDAIIYVVDSMDRDRIGISKQELVS
MLEEDELKKAVLVVFANKQDIEGAMTVTEVANSLGLPALKNRKYQIFKTSAIKGEGLDEAMEWLTNTLVQAK

**>ninaC**

MYMHAMKADNPPHIFAVADQSYQMMMHNKHCQCIVISGESGAGKTESANLLVQQLTQLGKAPNRTLEDRILQVNPLMEAFGNAKTVINDNSSRFGKYLEMFF
TASSGTVVGAKITEYLLEKSRVIHQAVGEQNFHIFYYLHDGLMSSDRQAEYHLKPHTVYKYIAEYTERPIEISSISINRVKFRAIEHCFEIIGFKKEEVSSVYGILVAILQTG
NIEFVAKDSGYGGDACVVSNTDLISIVSDLLGLDYTDLLECLTTTGMVAKGEVIIRDNSVQESMDARDAMAKALYGRLFSWIVNRISSLLKPSHRDSQDDQFYMVG
LLDIFGFENFKSNSFEQLCINIANEQIQYYFNQHIFAWELEEYKNEQIDAAEVAYVDNRPILDMFLAKPVGLLSLLDEESHFPKATDTTLVEKFHQNIKSSIYSRPKSNEL
SFGIDHYAGKVEYEAQGFLEKNRDRLPVEITNIMRLSENTVVKSLFQTPLTKTGNLQSGSLHSSSRNSSRGGSPMPTSPGVTIATYSSKGSSMGGSRIYNHLVSIYIEL
KLFKVQP

**>Egh**

MDILSMAQLLLYKSLQRITLLNCPRILQHLVRPLSLVSMRHVKHFLAVLLLVSLICLSVHVHWIVCKPNFSAIEHYRVSVGILLCLIQISVVFSLPFFIFNFLGVVLINVFTQ
RPKLQRSPLLGPFLCFRVVTKGLYPELVNENMKLNLQVCRKVGLHNYIFEVVTDIPINLKSTKDSREILVPKNYKTSSGSLFKARALQYALEPEVDILSPGDWIVHLDEE
TLLTEEAVIGIINFTGAGEHAFGQGVITYGSGKIINWVTTLADSIRVGVDYGCLRFTLGILHKPVFSWKGSFIVADAEAEKKISFDHGPEASIAEDCFFACVAFSKGYSF
GFVEGEMLEKSTFTLKDFIQQRRRWMQGILLTALSAKIPLRYKLGPVLMSFSGCTLPANLLMVPISLIWPLPSSPIITVTYSFILGTILYLFILGTLQTFSVHRYGIWKCILL
VLATMMCGLVACILENIASILVFWFPRSSVASFYVVKKDIPNDCVKGI

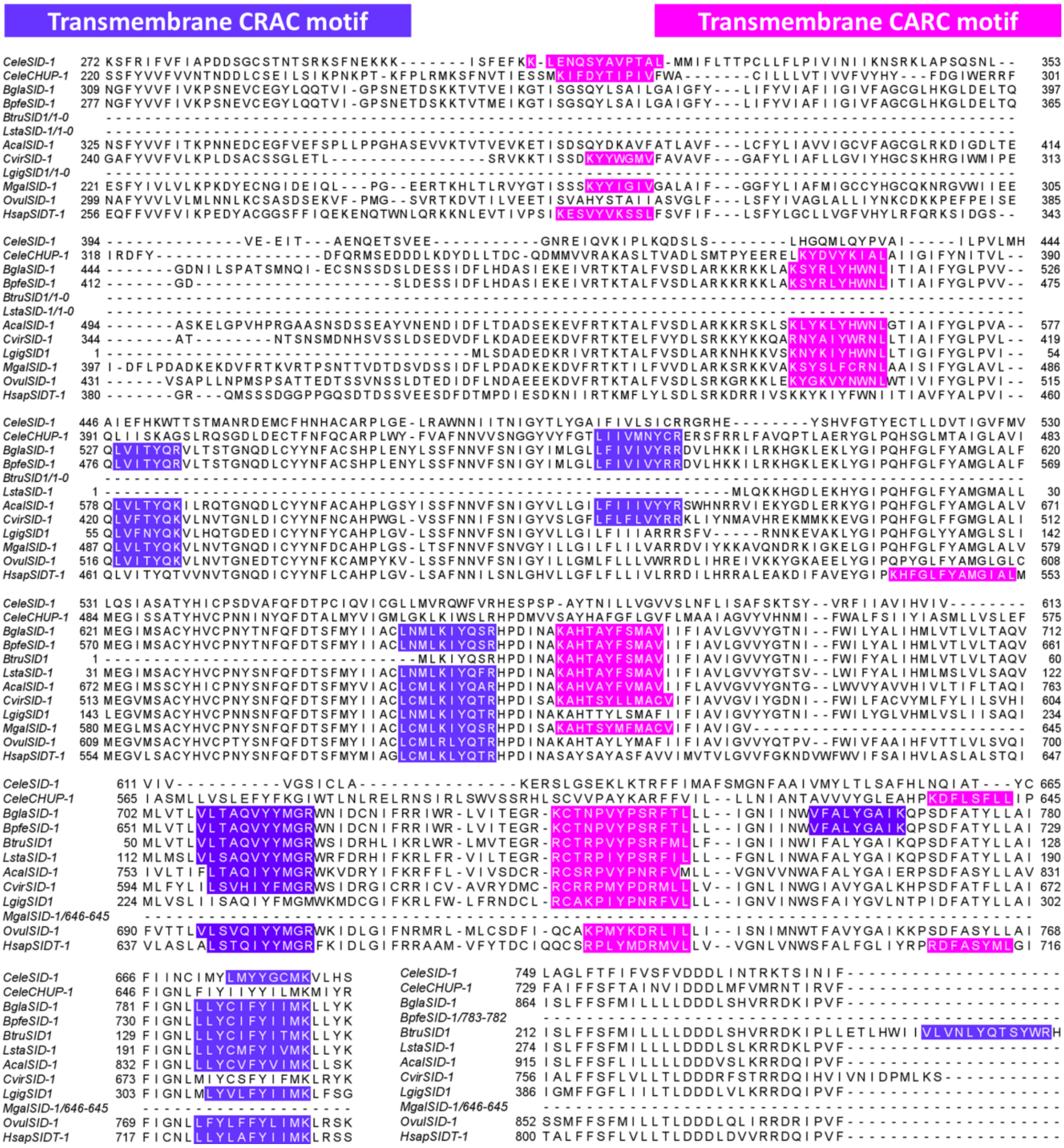


**Figure 1.** Multiple alignments of SID-1/CHUP-1 sequences of *C. elegans*, *H. sapiens*, *Biomphalaria* spp., *Bu. truncatus*, *L. stagnalis*, and other selected molluscs. Component residues of the CRAC and CARC motifs are highlighted in purple and pink, respectively. “-” indicates absence of an amino acid residue.

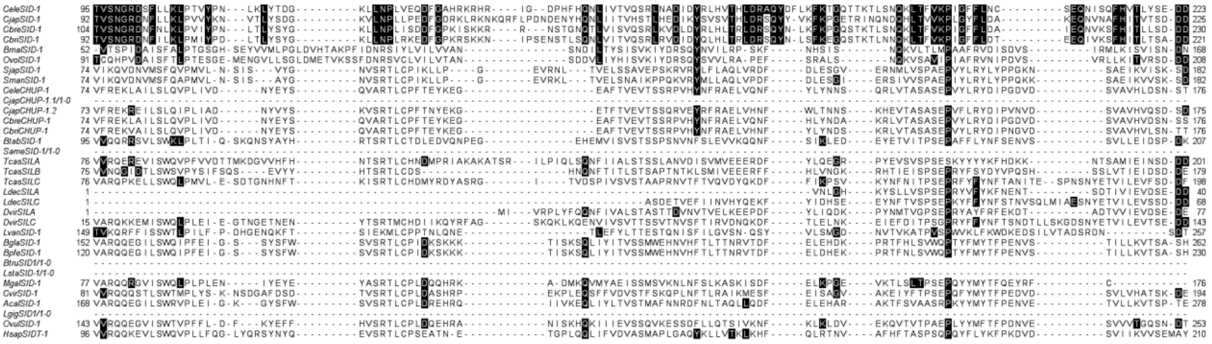

**Figure 2.** Alignment of partial extracellular N-terminal domain sequences of *C. elegans* SID-1 (amino acid T95–D223, containing the conserved RNAi uptake motifs) (Whangbo et al., 2017), with other SID-1/CHUP-1 peptide sequences in nematodes (*Cjap, Cbre, Cbri, Bmal, Ovol*), trematodes (*Sjap, Sman*), insects (*Btab, Same, Tcas, Ldec, Dvir*), crustaceans (*Lvan*), molluscs (*Bgla, Bpfe, Btru, Lsta, Mgal, Cvir, Acal, Lgig, Ovul*) and mammals (*Hsap*). All sequences were obtained as follows: *Caenorhabditis elegans* (*Cele*SID-1: CE30331, *Cele*CHUP-1: CE26397 – WormBase); *C. japonica* (*Cjap*SID-1: JA60059, *Cjap*CHUP-1.1: JA41534, *Cjap*CHUP-1.2: JA52591 – WormBase); *C. brenneri*, *Cbre*SID-1: CN02065, *Cbre*CHUP-1: CN03240 – WormBase); *C. briggsae* (*Cbri*SID-1: CBP22114, *Cbri*CHUP-1: CBP03753 – WormBase); *Brugia malayi*, *Bmal*SID-1 (BM45574 – WormBase); *Onchocerca volvulus*, *Ovol*SID-1 (OVP05450 – WormBase); *Schistosoma japonicum*, *Sjap*SID-1 (BAH22347 – GenBank); *S. mansoni*, *Sman*SID-1 (XP_018649472.1 – NCBI RefSeq); *Bemisia tabaci*, *Btab*SID-1 (XP_018910555.1 – NCBI RefSeq); *Schistocerca americana*, SameSID-1 (AY879097 – GenBank); *Tribolium castaneum* (*Tcas*SILA: EF688527, *Tcas*SILB: EF688528, TcasSILC: EF688529 – GenBank); *Leptinotarsa decemlineata* (*Ldec*SILA: AKE50508.1, *Ldec*SILC: AKE50509.1 – GenBank); *Diabrotica virgifera* (*Dvir*SILA, *Dvir*SILB – (Miyata et al., 2014)); *Litopenaeus vanname*i, *Lvan*SID-1 (ADK25179.1 – GenBank). Refer to the previous pages of this supplementary material for other accession numbers.

**REFERENCES**


**Miyata K, Ramaseshadri P, Zhang Y, Segers G, Bolognesi R and Tomoyasu Y** (2014). Establishing an in vivo assay system to identify components involved in environmental RNA interference in the western corn rootworm. *PLoS One*, **9**(7), e101661. doi: 10.1371/journal.pone.0101661

**Whangbo JS, Weisman AS, Chae J and Hunter CP** (2017). SID-1 domains important for dsRNA import in *Caenorhabditis elegans*. *G3 (Bethesda)*, **7**(12), 3887-3899. doi: 10.1534/g3.117.300308
